# BATOseq-PE: A public batoid DNA library with the first molecular insights into the cryptic diversity, genetic health, and management of Peruvian marine rays

**DOI:** 10.64898/2026.09.15.751127

**Authors:** Alan Marín, Luis E. Santos-Rojas, Renato Gozzer-Wuest, Angel Yon-Utrilla, Santiago Vigo-López, Stefany Rojas-Perea, Claudio Villegas-Llerena, Solange R. Paredes-Moscosso, Edgar A. López-Landavery, Lorenzo E. Reyes-Flores, Eliana Zelada-Mázmela

**Affiliations:** Laboratorio de Genética, Fisiología y Reproducción, Facultad de Ciencias, Universidad Nacional del Santa, Chimbote, Perú; Innovations for Ocean Action Foundation (I4OA), Lima, Peru; The Manta Trust, Catemwood House, Norwood Lane, Corscombe, Dorset DT2 0NT, UK; Biomolecules Laboratory, Faculty of Health Sciences, Universidad Peruana de Ciencias Aplicadas (UPC), Lima 15023, Perú; Escuela de Biología, Facultad de Ciencias de la Salud, Universidad Peruana de Ciencias Aplicadas (UPC), Lima 15023, Perú

**Keywords:** Chondrichthyes, Cytochrome c oxidase subunit I, DNA barcoding, Eastern South Pacific Ocean, Manta rays, Mobulids

## Abstract

Batoids (rays, skates, and relatives) are among the most threatened cartilaginous fishes globally, yet the molecular diversity of Peruvian taxa remains poorly evaluated. Furthermore, regional barcode sequences for high-conservation-priority taxa, such as sawfish, manta, and devil rays, are often restricted to private records, hindering essential phylogeographic and population structure assessments. To bridge this gap, we developed BATOseq-PE, the first public genetic reference library for Peruvian marine batoids, covering 14 species sequenced *in vitro* from local specimens, complemented by a novel *in silico* assembled mitogenome for *Notoraja martinezi*. This open-access repository contributes 73 novel COI barcodes, providing robust taxonomic resolution across 15 molecular operational taxonomic units (MOTUs) and Barcode Index Numbers (BINs). Notably, we provide the first Peruvian public records for high-priority species such as *Mobula munkiana*, *Mobula thurstoni*, *Myliobatis peruviana*, *Rostroraja velezi*, and the first global public barcodes for *N*. *martinezi* and *Zapteryx xyster*. By coupling BATOseq-PE with global datasets, we conducted the first multi-taxa screening for delimitation and phylogeographic assessments on Peruvian rays, revealing a cryptic lineage within *R. velezi* and genetic connectivity patterns suggesting female philopatry in *Mobula mobular*. Finally, by integrating nucleotide diversity into a Genetic Diversity Risk Indicator (GDRI) alongside ecological and fishery data, we identified an alarming trend where heavy fishing pressure overlaps with extinction risk and depleted genetic diversity, allowing us to suggest data-driven, tailored management strategies. BATOseq-PE represents a valuable resource to accelerate taxonomic and population assessments, ultimately fostering ray conservation through the integration of genetic frameworks into fishery management.

## 1. Introduction

The infraclass Batoidea (rays, skates, guitarfishes, and sawfishes) is a diverse group of cartilaginous fishes that primarily inhabit marine ecosystems worldwide, with fewer species found in freshwater habitats (Last et al. 2016b). This group comprises over 730 species, categorized across four extant orders and 26 families (WoRMS database, as of June 2026; Froese and Pauly 2026). Although intrinsic chondrichthyan traits, such as low fecundity and slow growth, render batoids highly vulnerable to overfishing, most species still support substantial artisanal fisheries globally (Last et al. 2016b; Barrowclift et al. 2023; FAO 2024). In addition, most batoid species are benthic or demersal, relying heavily on the seabed for foraging and shelter; consequently, they are often caught accidentally as commercial bottom-trawling bycatch (Last et al. 2016b). Therefore, the cumulative impacts of their biological vulnerability, targeted fishing, bycatch, and habitat degradation have severely depleted multiple batoid populations, underscoring the critical need for global conservation frameworks (Dulvy et al. 2021).

Peru is a major fishing nation with a high diversity of batoid species (Chirichigno and Cornejo 2001; Marín et al. 2018, 2026; Campos-León et al. 2026). This richness is driven by the confluence of two main currents: the Northern Humboldt Current and the North Equatorial Counter Current, which create a highly productive transition zone that supports diverse taxa (Velez-Zuazo et al. 2024; Campos-León et al. 2026). Consequently, up to 37 marine ray species are reported to inhabit the Peruvian Sea (Campos-León et al. 2026), including one of the world’s largest populations of the flagship conservation species, the giant manta ray, *Mobula birostris* (Harty et al. 2022). With about 25 batoid species holding commercial value (Zavalaga et al. 2021), four main taxonomic groups account for the vast majority of historical Peruvian batoid landings. These include eagle rays (*Myliobatis* spp.) and devil rays (*Mobula* spp.) (hereafter abbreviated as *My*. spp. and *Mo*. spp., respectively), guitarfishes (*Pseudobatos* spp.), and the diamond stingray, *Hypanus dipterurus* (González-Pestana et al. 2022), which is provisionally listed herein as *H. brevis* following Marín et al. (2026).

DNA-based methods are widely utilized in batoid research for accurate species identification, the clarification of taxonomic boundaries, and the evaluation of population structure (Vargas-Caro et al. 2017; Sales et al. 2019; Marín 2025, 2026; Cunha et al. 2026; Rodrigues-Filho et al. 2026). During the last two decades, the application of genetic markers has led to major taxonomic revisions of global batoid diversity, directly impacting the management and conservation of numerous species around the world (Naylor et al. 2012b; Last et al. 2016b). As a result, genetic studies have reshaped multiple taxonomic tiers, such as families (e.g., Aetobatidae, Arhynchobatidae), genera (e.g., *Hypanus*, *Mobula*, *Rostroraja*), and species (e.g., *Aetobatus ocellatus*, *Mo. birostris*) (White et al. 2010, 2018; Naylor et al. 2012b; Poortvliet et al. 2015; Last et al. 2016a; White and Naylor 2016).

While contemporary chondrichthyan systematics heavily favor the NADH dehydrogenase subunit 2 (ND2) marker due to its higher mutation rate (Naylor et al. 2012a, b), cytochrome c oxidase subunit I gene (COI) sequences remain the most abundant locus in public repositories (Kottillil et al. 2023). Furthermore, molecular studies of chondrichthyan taxa from geographic regions with low sequence representativeness, such as the Eastern South Pacific Ocean (ESPO), rely primarily on COI barcodes (Velez-Zuazo et al. 2015, 2021; Marín et al. 2018, 2022, 2026; Biffi et al. 2020; Alfaro-Cordova et al. 2023; Dufflocq et al. 2026; Marín 2026). In addition, the COI’s nucleotide diversity (*π*) has proven useful in revealing population declines and consequently it can be used as a proxy for the global conservation status of fish species (Petit-Marty et al. 2021, 2022), including batoids (Ferragut-Perello et al. 2023, 2026).

Recently, the COI-*π* metric was integrated into the FishBase database as part of its Genetic Diversity Risk Indicator (GDRI; available at FishBase). The GDRI is an open-access tool that uses this metric to assess the genetic health of fish populations, classifying them into three conservation risk categories: low, medium, and high. This tool is particularly valuable for monitoring range-restricted species, which face heightened vulnerability to various threats such as genetic erosion (Hobbs et al. 2013). Applying such genetic indicators could prove vital for Peru’s batoid diversity, given that over one-third of these species are regionally endemic taxa (Campos-León et al. 2026). Commercially exploited species such as *H. brevis*, *My. chilensis*, *My. peruviana*, and *Urotrygon chilensis* occur exclusively within the ESPO or the Humboldt Current Large Marine Ecosystem (Last et al. 2016b, Zavalaga et al. 2021; Ehemann et al. 2024a; Marín et al. 2026). Yet, despite their vulnerability and the heavy, unregulated exploitation, local molecular initiatives targeting endemic batoids remain scarce.

Regrettably, the stock boundaries, reproductive connectivity, and spatial demographic patterns of Peruvian batoid populations remain poorly characterized. Currently, regional molecular data remain fragmented and primarily limited to species-level DNA barcoding for seafood traceability and species diversity assessments (Marín et al. 2018; Rosas et al. 2018; Biffi et al. 2020; Cabanillas-Torpoco et al. 2020; Velez-Zuazo et al. 2021; Siccha-Ramirez et al. 2022; Alfaro-Shigueto et al. 2025; Zavala et al. 2025). Furthermore, a significant share of regional molecular records for key conservation species, such as for the sawfish *Pristis pristis* (e.g., BOLD ID PMFSH636-20) and *Mo. birostris* (e.g., BOLD IDs DBPD001-21 to DBPD005-21), are still kept as private within BOLD, constraining their integration into subsequent research. Additionally, to date, only two baseline studies have quantified genetic metrics including haplotype number (*H*), haplotype diversity (*H_d_*), and nucleotide diversity (*π*), in local populations of devil rays (Alfaro-Cordova et al. 2023) and diamond stingrays (Marín et al. 2026).

To date, no molecular study has focused comprehensively on the entire marine batoid fauna of Peru. Local genetic studies involving batoids have been restricted either to multi-species fish identifications (e.g., Marín et al. 2018; Siccha-Ramirez et al. 2022; Alfaro-Shigueto et al. 2025; Zavala et al. 2025) or to narrow taxonomic groups, such as sawfish (Cabanillas-Torpoco et al. 2020) and mobulid rays (Alfaro-Cordova et al. 2023). Hence, there is a critical research gap and urgent need for a dedicated, comprehensive research effort assessing the current molecular knowledge of Peru’s batoid species. Such an investigation effort is essential to advance future research in conservation genetics, phylogenetics, population genetic structure, and taxonomy of Peruvian batoids. Moreover, a dedicated molecular review will help identify batoid species vulnerable to different threats including localized losses of genetic diversity, severe genetic bottlenecks, and cryptic diversity (Petean et al. 2024; Marín et al. 2026; Rodrigues-Filho et al. 2026).

Given the lack of studies focused solely on the molecular identification of batoid diversity and the limitations of private databases, this study aimed to expand the regional COI reference library by introducing a public, dedicated genetic repository for Peruvian batoids. Novel sequences were obtained from specimens collected from different commercial venues across the Peruvian coast. Additionally, the first complete mitogenome for the barbedwire-tailed skate, *Notoraja martinezi*, was assembled and characterized using public genomic reads, providing its first global public COI reference. All novel genetic data were integrated into public databases to maximize geographic and taxonomic coverage. To evaluate phylogeographic and species boundaries, robust molecular reconstructions and delimitation analyses were conducted. Finally, the COI-*π* metric was integrated with conservation status, endemism metrics, and landing statistics, to establish the first regional prioritization framework for identifying species requiring immediate research, management, and conservation actions.

## 2. Materials and methods

### Ethics statement

Tissue samples utilized in this research were obtained exclusively from deceased specimens available via different seafood facilities. Therefore, no live animals were sacrificed or harmed for the purposes of this study. Field collection did not require any special permits as no live or protected fauna were targeted.

### Sampling and tissue collection

To maximize taxonomic coverage, we conducted a non-probabilistic, opportunistic, and intensive sampling strategy targeting all available batoid diversity during each survey day. Most samplings were strategically conducted in northern Peru (Fig. 1), a region harboring the vast majority of the nation’s batoid diversity, characterized by high landings and strong demand for regional gastronomy (González-Pestana et al. 2022, 2024; Rojas-Perea et al. 2025; Marín et al. 2026). Thus, artisanal fish landing sites in the Tumbes region alongside two key wholesale fish markets in the Piura (José Olaya) and Lambayeque regions (ECOMPHISA Santa Rosa) were surveyed. Additional sampling sites comprised landing sites, artisanal fishing ports, and local markets from the Ancash and Ica region regions.

**Fig. 1.**
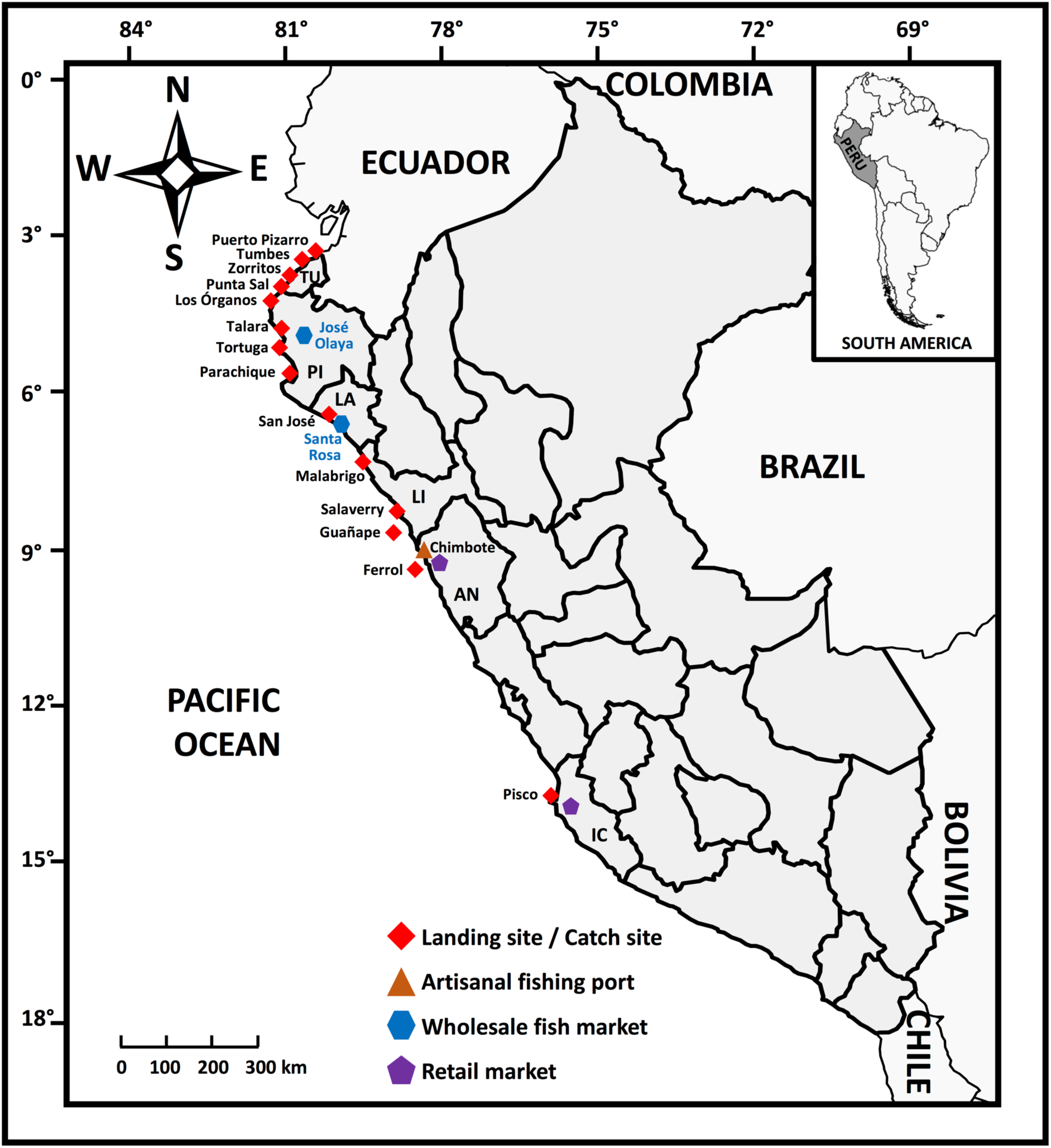
Sampling locations for batoid specimens along the Peruvian coast. Commercial venues include landing and catch sites, artisanal fishing ports, wholesale fish markets, and retail markets. Peruvian region codes indicate Tumbes (TU), Piura (PI), Lambayeque (LA), La Libertad (LI), Ancash (AN), and Ica (IC)

A total of 75 batoid samples were collected over a nine-month period (November 2025 to August 2026). This dataset comprised 73 tissue samples obtained from whole specimens or partial bodies of fresh organisms. Two additional processed commercial products of dried guitarfish meat, locally known as “*chinguirito*”, were also collected to assess seafood label accuracy. We interviewed each fisher and vendor to record the reported catch site for each collected sample (Table 1). Additional collected data included commercial labels, product presentation, and retail price per kg.

**Table 1.**
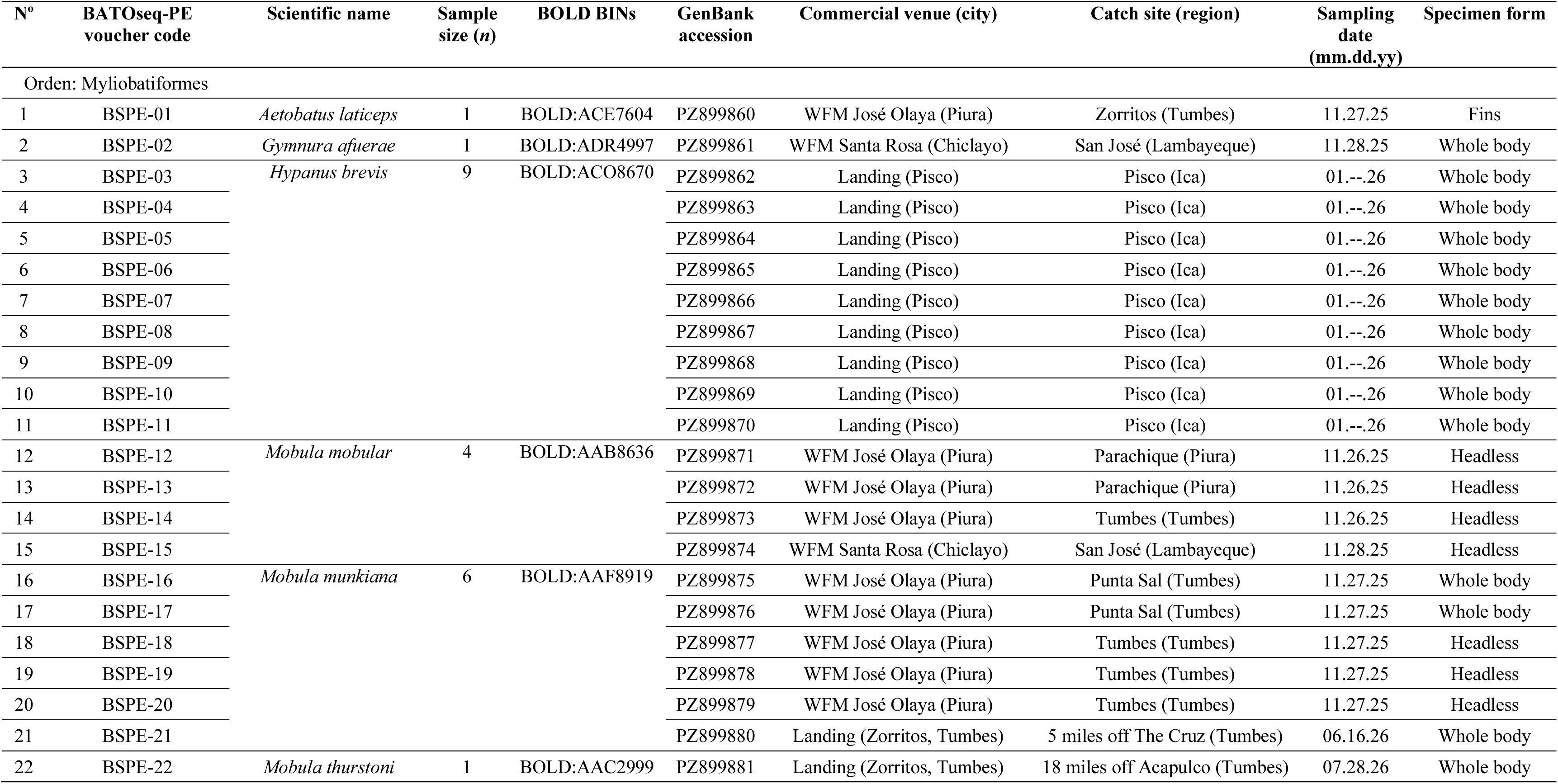
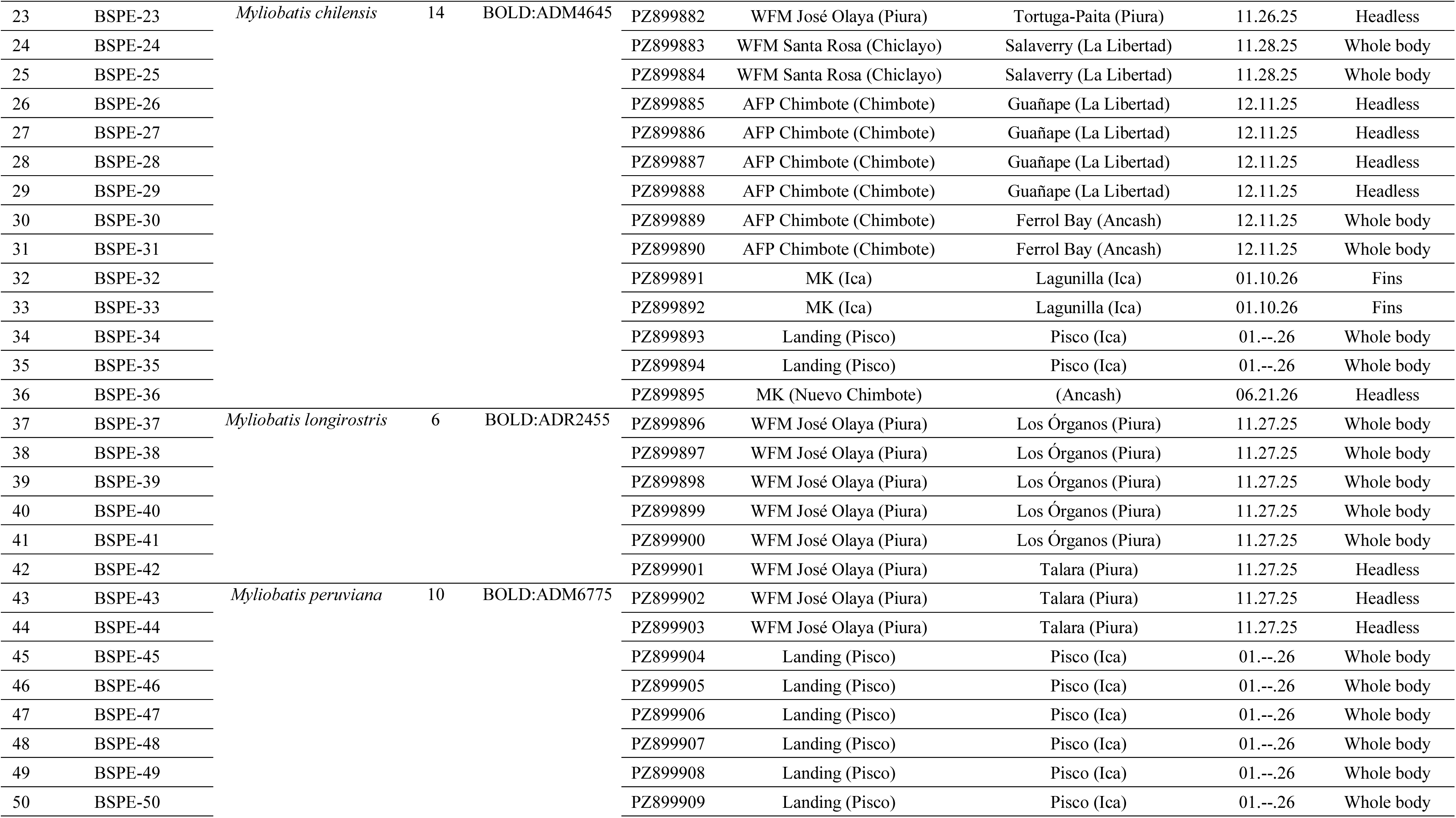
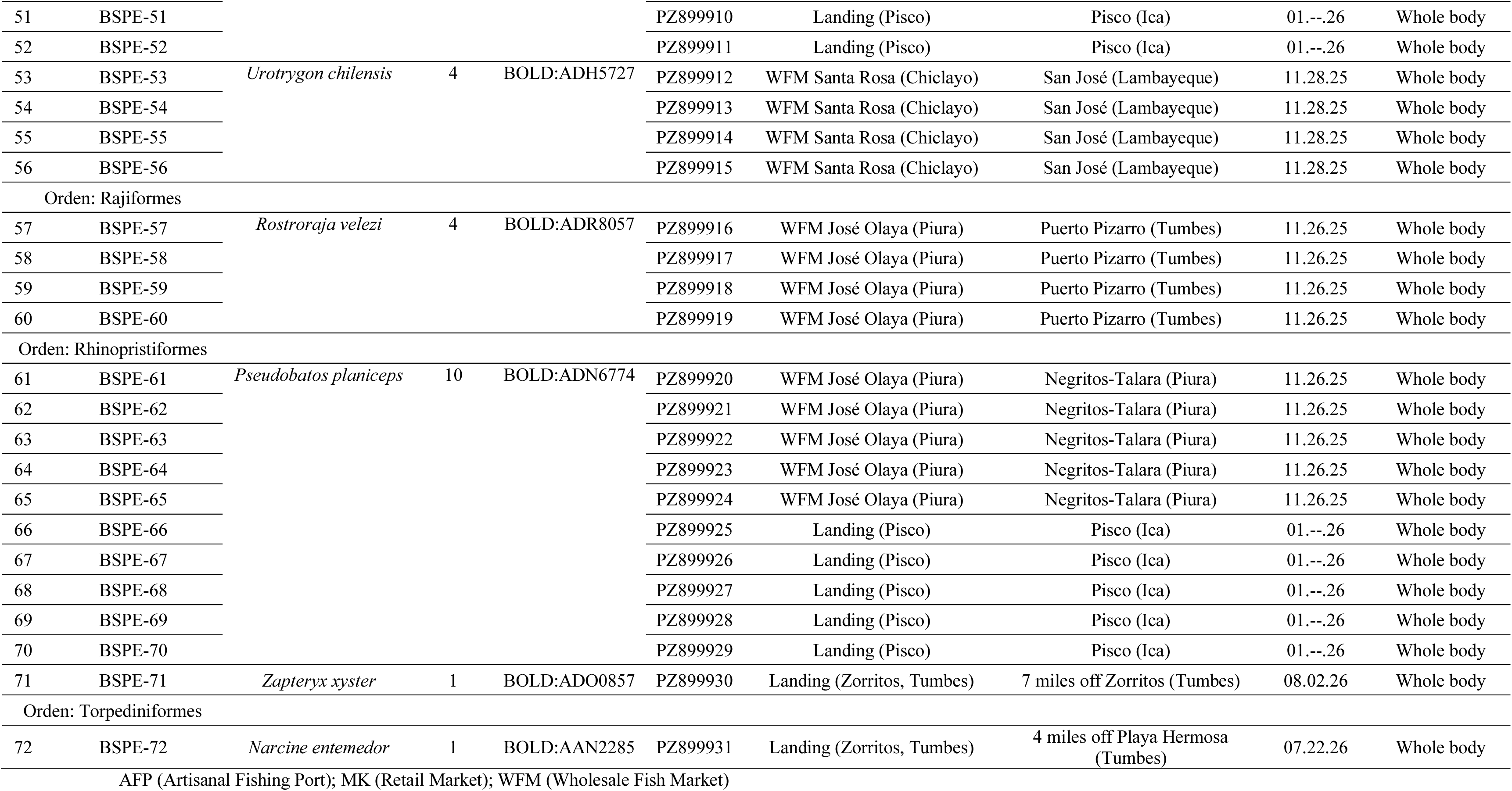
Metadata of batoid specimens collected from different Peruvian locations and commercial venues. Voucher codes, BOLD Barcode Index Numbers (BINs), GenBank accession numbers, commercial venues, geographic provenance, and sampling dates are listed

The majority of samples were collected from intact, whole individuals (Fig. 2) where tissue sampling and diagnostic digital photographs were taken *in situ*. Exceptionally, four specimens (identified as *Mo. munkiana*, *Mo. thurstoni*, *Narcine entemedor*, and *Zapteryx xyster*) were purchased as whole animals. For these four individuals, morphometric parameters including total weight, total length, and disc measurements were recorded alongside tissue sampling and digital photographs, although the physical carcasses could not be preserved due to storage constraints. For all collected samples, muscular or fin tissue fragments (∼1–2 cm³) were surgically removed using sterile scalpel blades, placed in 1.5 mL microtubes containing 96% ethanol, transported to the laboratory, and stored at –15 °C. All tissue samples were deposited in the fish genetic collection of the Laboratory of Genetics, Physiology, and Reproduction (LGPR) of the National University of Santa (Ancash, Peru) under unique catalog codes linked to their respective genetic sequences.

**Fig. 2.**
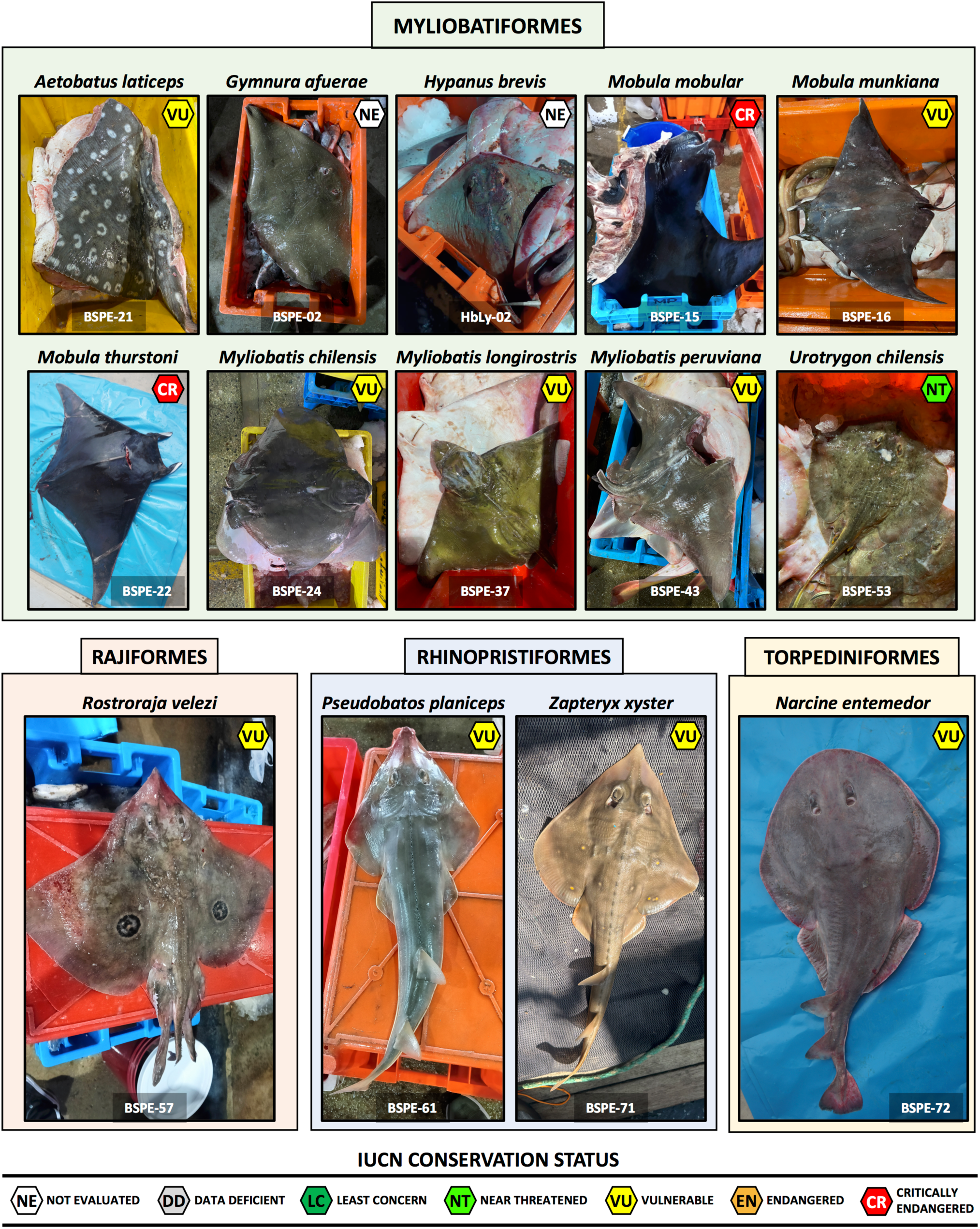
Representative photographic records of the 14 batoid species sampled from Peruvian commercial venues. Specimens are organized under their respective taxonomic orders: Myliobatiformes, Rajiformes, Rhinopristiformes, and Torpediniformes. The individual BATOseq-PE voucher code is provided at the bottom of each panel. The photograph for *Hypanus brevis* (HbLy-02) represents the single specimen external to the BATOseq-PE reference library and was taken during the field sampling of Marín et al. (2026)

### Taxonomic identification

For intact whole specimens, morphological identifications were performed using complete organisms or digital photographs in conjunction with a taxonomic key for Peruvian marine fishes (Chirichigno and Velez 1998) and specialized regional field guides and inventories (Zavalaga et al. 2021; Siccha-Ramirez et al. 2022). For heavily processed catches, digital vouchers were limited to isolated diagnostic fragments (e.g. spotted patterns in *A. laticeps*).

### DNA extraction, PCR amplification, and sequencing

The molecular experiments were carried out entirely at the LGPR. Genomic DNA was isolated with the CTAB method following Zuccarello and Lokhorst (2005), with minor modifications described in Marín et al. (2026). A partial COI fragment was amplified with four different universal primer sets designed by Folmer et al. (1994), Ward et al. (2005), and Ivanova et al. (2007). Primer sequences, PCR mix composition, and the amplification protocols used in target taxa are shown in File S1.

Bidirectional Sanger sequencing was performed using the same PCR primers, with the protocol and equipment described in Marín et al. (2026). Electropherograms were manually checked using MEGA7 (Kumar et al. 2016). Primer hybridization regions were removed from both strands and final consensus sequences were obtained by aligning complementary forward and reverse sequences. To rule out the amplification of nuclear pseudogenes (NUMTs), we applied a final quality control step to all consensus DNA sequences by translating them into their corresponding amino acid sequences and checking for premature stop codons or frameshift mutations.

### Species identity determination

The assembled consensus DNA sequences were compared against reference libraries in the BOLD Systems Barcode ID engine version 5 (https://id.boldsystems.org/), searching the Animal Species-Level Library, which encompasses both public and private barcode records. Sequences were automatically clustered into Barcode Index Numbers (BINs) within the BOLD database, which utilizes the Refined Single Linkage (RESL) algorithm to delineate Molecular Operational Taxonomic Units (MOTUs). A sequence similarity threshold of ≥ 99% was used for definitive species-level taxonomic assignments. Additionally, to ensure diagnostic robustness, all taxonomic designations were cross-verified using the Basic Local Alignment Search Tool BLASTn algorithm (https://blast.ncbi.nlm.nih.gov/) against the nucleotide database from the National Center for Biotechnology Information (NCBI). Finally, current accepted taxonomic classification and scientific names were verified using the Eschmeyer’s Catalog of Fishes (https://researcharchive.calacademy.org/research/ichthyology/catalog/fishc-atmain.asp) and the WoRMS database (https://www.marinespecies.org/).

### Genetic distances and barcoding gap assessments

To evaluate the taxonomic resolution of our BATOseq-PE library and assess the existence of a definitive barcoding gap among the analyzed Peruvian rays, intra- and interspecific pairwise genetic distances were estimated using the Kimura 2-parameter (K2P) distance model (Kimura 1980). Analyses were executed across two parallel frameworks including local and extended datasets. The local BATOseq-PE library dataset consisted strictly of the regional sequences generated *in vitro* by this study. The expanded contextual dataset combined sequences from the BATOseq-PE library, the novel *N. martinezi* COI barcode assembled *in silico*, and public records sampled in Peru and abroad (when regional sequences were not available) to encompass a total of 28 Peruvian batoid species. A total of 72 genetic barcodes for the local baseline and 113 sequences for the expanded dataset were multialigned and trimmed using MEGA7, resulting in alignments of 548 bp and 517 bp, respectively.

The alignments were subsequently imported into R (R for Mac OS X GUI version 4.2.3) (R Core Team 2024), where pairwise genetic distances were calculated via the Analysis of Phylogenetics and Evolution (ape) package, using the K2P model. The results for both datasets were visualized and compared using customized genomic distance histograms.

### Taxonomic coverage and public sequence abundance in repositories

We screened both the BOLD and GenBank repositories for all publicly available COI barcodes of the 37 Peruvian marine batoid species listed in the most recent taxonomic inventory of regional chondrichthyan diversity by Campos-León et al. (2026). To eliminate potential bias arising from misidentifications or outdated nomenclature, a rigorous taxonomic reconciliation process was implemented. Thus, sequences assigned to the Atlantic *A. narinari* from Pacific waters were reclassified as *A. laticeps* (White et al. 2010; Last et al. 2016b). Similarly, *Hypanus dipterurus* records from the Eastern North Pacific were disregarded as this species is restricted to the Northern Hemisphere (Marín et al. 2026). Also, records assigned to the genus *Urotrygon* from the ESPO were treated under the single verified southern lineage *U. chilensis* (Ehemann et al. 2024a). Additionally, all *Mo. japanica* records were assigned to its senior synonym and valid taxon *Mo. mobular* (Last et al. 2016b; White et al. 2018). Furthermore, public records within the genera *Mobula*, *Pseudobatos*, and *Tetronarce* were audited to incorporate earlier sequences labeled under their obsolete junior synonyms *Manta*, *Rhinobatos*, and *Torpedo*, respectively (Last et al. 2016b; White et al. 2018).

Finally, to account for critical geographical variations in distribution ranges, sampling location metadata from selected records were cross-verified with current databases (FishBase, IUCN, WoRMS) and relevant literature reporting updated geographic boundaries (Zavalaga et al. 2021; Roque-Sanchez and Donayre-Salazar 2025; Campos-León et al. 2026). All retrieved COI barcodes were stratified by geographic origin to distinguish between the four ESPO nations (Colombia, Chile, Ecuador, and Peru) and an “others” category for all remaining countries.

### Mining of public genomic data for Eastern South Pacific batoids

To expand the taxonomic coverage of genetic barcodes for Peruvian batoids, we searched the Sequence Read Archive (SRA) database (https://www.ncbi.nlm.nih.gov/sra) for available genomic data from ESPO batoids. We identified short-read sequences (SRS) data derived from whole-genome sequencing (WGS) experiments for two species missing public COI references within both the BOLD and GenBank datasets. The first taxon, *N. martinezi*, was represented by public genomic libraries from a specimen collected off the Pacific coast of Costa Rica (BioSample SAMN48911751, Run SRR33849666). These data were selected for downstream bioinformatics processing via the Galaxy webserver (https://usegalaxy.eu) to assemble the first complete mitogenome for this species. The second taxon detected was the whitesnout guitarfish, *P. leucorhynchus*, which featured two WGS datasets from museum specimens (BioSamples SAMN45906730 and SAMN45906740; Runs SRR31776815 and SRR31776813, respectively). However, due to the extremely low sequencing depth of these historical DNA libraries, subsequent bioinformatics efforts failed to recover either the mitogenome or diagnostic COI barcode sequence for this species.

### Mitogenome assembly and annotation for *Notoraja martinezi*

High-throughput SRS data deposited under GenBank accession SRR33849666 were downloaded using the Faster Download and Extract Reads in FASTQ tool version 3.1.1 (Leinonen et al. 2010). Raw FASTQ reads were subsequently subjected to quality filtering and adapter trimming using Trim Galore! version 0.6.10 (Krueger 2021). The filtered reads were assembled *de novo* with GetOrganelle version 1.7.7.1 (Jin et al. 2020), enforcing a maximum threshold of 20 million reads to optimize assembly efficiency. Lastly, the resulting circular consensus sequence was annotated using the MITOS2 pipeline version 2.1.10 (Donath et al. 2019) with default parameters for vertebrate mitochondrial codes.

Finally, the species identity of BioSample SAMN48911751 was evaluated by comparing the 5’ region of its COI sequence against the reference sequences deposited in both the BOLD and the GenBank databases.

### Phylogenetic analysis

To infer the phylogenetic relationships of Peruvian marine batoids and confirm species identity, we combined our newly COI sequences (generated *in vitro* and *in silico*) with conspecific reference sequences retrieved from public BOLD and GenBank databases. Whenever possible, we prioritized the inclusion of samples from Peru. However, to cover the taxonomic gap generated by the lack of regional reference sequences, we included barcodes from eight species collected in foreign countries. These included *Discopyge tschudii*, *Gurgesiella furvescens*, *Rajella nigerrima*, and *Tetronarce tremens* from Chile; *U. pardalis* from Costa Rica; *G. crebripunctata* and *Mo. birostris* from Mexico; and *Mo. tarapacana* from Colombia.

Additionally, three freshwater species of the family Potamotrygonidae (*Heliotrygon rosai*, *Paratrygon ajereba*, and *Potamotrygon motoro*) and three foreign auxiliary myliobatiform taxa (*Plesiobatis daviesi*, *Urolophus cruciatus*, and *U. paucimaculatus*) were added to provide topological stability during the phylogenetic reconstruction. However, these species were excluded from the downstream diversity analysis and richness estimations of the regional marine dataset.

DNA sequences were aligned using ClustalW algorithm (Thompson et al. 1994) implemented in MEGA7 (Kumar et al. 2016). The final aligned dataset (597 bp) consisted of 149 barcodes including 72 novel sequences from the BATOseq-PE library, the novel record for *N. martinezi* assembled herein, and 76 reference sequences retrieved from the BOLD or GenBank databases. The outgroups comprised two Holocephali taxa (*Callorhinchus callorynchus* and *C. milii*). Additional outgroup species included two primitive Squalimorphi sharks (*Hexanchus griseus* and *Squalus acanthias*) and two modern Galeomorphi sharks (*Heterodontus zebra* and *Sphyrna zygaena*).

For the phylogenetic tree construction, a maximum likelihood (ML) approach was conducted with IQ-TREE version 2.4 (Minh et al. 2020) via the Galaxy webserver. The analyses were partitioned by codon position with best substitution models determined with ModelFinder (Kalyaanamoorthy et al. 2017) and statistical node support conducted with 1,000 replicates of the Ultrafast Bootstrap (UFBoot2) (Hoang et al. 2018). The final phylogenetic consensus tree was visualized in FigTree version 1.4.4 (http://tree.bio.ed.ac.uk/software/figtree/) and edited using Inkscape (https://inkscape.org).

### Species delimitation analyses

To maintain analytical efficiency and focus on unresolved taxa, we excluded lineages whose species boundaries have recently been resolved using Peruvian samples, specifically: *Gymnura* (Ehemann et al. 2024b; Gales et al. 2024), *Hypanus* (Marín et al. 2026), *Narcine* (Rodrigues-Filho et al. 2026), *Rhinoptera* (Cunha et al. 2026), and *Urotrygon* (Ehemann et al. 2024a). Additionally, species from the genus *Myliobatis* were intentionally excluded from our delimitation analyses to respect and prevent overlapping with an ongoing, independent taxonomic project currently focusing on this group. We selected species whose pairwise intraspecific genetic distances exceeded a K2P distance threshold of 0.8%. This inclusive baseline is justified by the narrow boundary zones of batoid genetic variation documented by Crobe et al. (2021) and Petean et al. (2024).

Five different algorithms comprising distance-, tree-, and coalescent-based approaches were employed for species delimitation analyses. These include 1) the network-based Refined Single Linkage (RESL) method; 2) the Assemble Species by Automatic Partitioning (ASAP) method; 3) the Bayesian Poisson Tree Processes (bPTP) approach; the 4) the PTP-ML model; and 5) the General Mixed Yule Coalescent (GMYC) model. The distance- and network-based RESL method is implemented in the BOLD database to automatically cluster sequences into MOTUs termed BINs, based on a pre-defined sequence divergence threshold (Ratnasingham and Hebert 2013). To complement this, the distance-based ASAP approach (Puillandre et al. 2021) was conducted using the Spart explorer interface (https://spartexplorer.mnhn.fr/delimitation) under the K2P substitution model and the transition/transversion rate ratio set to 2. Two tree-based models (bPTP and PTP-ML) (Zhang et al. 2013) were conducted in the bPTP server (https://species.h-its.org/ptp/) using default parameters (100,000 MCM generations; thinning: 100; burn-in: 0.1). The non-ultrameric phylogenetic tree needed for these analyses was obtained via IQ-TREE with the same parameters presented in the “*Phylogenetics analysis*” section.

Finally, the GMYC algorithm (Fujisawa and Barraclough 2013) was conducted via the GMYC web server (https://species.h-its.org/gmyc/) under default settings. The ultrameric phylogenetic tree required for this approach was constructed with BEAST version 1.1 (Drummond and Rambaut 2007) under a Bayesian framework. The COI dataset was partitioned by codon position, with each partition assigned to its optimal evolutionary substitution model as determined by ModelFinder (Kalyaanamoorthy et al. 2017). The Markov chain Monte Carlo (MCMC) simulation in BEAST was run for 10,000,000 generations, with parameter logs and tree topologies sampled every 1,000 generations. Convergence and stationarity of the chains were evaluated using Tracer version 1.7.2 (Rambaut et al. 2018), ensuring that the Effective Sample Size (ESS) exceeded 200 for all estimated parameters. After discarding 10% of sampled trees as burn-in, the maximum clade credibility (MCC) tree was annotated and compiled using TreeAnnotator (Suchard et al. 2018).

### Haplotype network, genetic diversity, and population structure

For the phylogeographic assessments, we selected those taxa with publicly available regional (Peru) and international (Eastern Pacific nations) sequences, enabling us to evaluate genetic variation across two or more populations. Conversely, two batoid species, *A. laticeps* and *H. brevis*, were intentionally excluded from these analyses; their phylogeography, haplotype networks, and population structures have been recently and comprehensively investigated by Marín et al. (2026) and Vinueza-Guarderas et al. (2026), respectively. Similarly, to avoid overlap with ongoing studies, *Myliobatis* taxa were excluded from our phylogeographic network analyses.

The relationships among haplotypes of selected species were examined using median-joining networks constructed in HaplowebMaker (Spöri and Flot 2020). Genetic indexes (*H*, *H_d_*, and *π*) were calculated using the ape package implemented in R. To detect departures from selective neutrality and reconstruct past demographic history, we performed Fu’s *F_S_* and Tajima’s *D* neutrality tests in Arlequin version 3.5 (Excoffier and Lischer 2010). Significant thresholds were determined via 10,000 coalescent simulations, using critical values of P < 0.02 for *F_S_* and P < 0.05 for *D*.

Genetic divergence within the selected ray samples was quantified via Analysis of Molecular Variance (AMOVA) in Arlequin. To avoid sampling artifacts, AMOVA was restricted to populations with a sample size of at least 10 individuals (*n* ≥ 10). Pairwise fixation indices (variation among groups: Φ*_CT_*; among populations within groups: Φ*_SC_*; and among all populations: Φ*_ST_*) were estimated based on the K2P distance model (Kimura 1980).

### Conservation status and national regulatory framework

The conservation status of Peruvian batoid species was retrieved from The International Union for Conservation of Nature (IUCN) Red List of Threatened Species (https://www.iucnredlist.org). National regulatory laws regarding batoid fisheries were screened from the legal regulations and documentation search server of the Ministry of Production (PRODUCE) (https://www.gob.pe/institucion/produce/normas-legales).

### Exploratory description of the batoid fishery data and trends

The database of chondrichthyan landings from 1996 to 2024, provided by the Peruvian Marine Research Institute (IMARPE) and curated by Marín et al. (2026), was used to retrieve information on batoid catches at the finest available scale. In total, 16 categories of species were included in the database, 12 at the scientific name level and 4 at the genus level. Also, two species were labeled with outdated scientific names, which were updated before analysis (see subsection “*Taxonomic coverage and public sequence abundance in repositories*”). The database was used to describe the consistency of species-level information across 29 years of data, as well as the magnitude of fishing activity by gear type and zone. In addition, a time series of total batoid landings per year was estimated and fitted to a simple linear regression.

Then, a yearly count of records was estimated to detect, by simple inspection, any changes in the amount of data collected by year. In this regard, the number of records in the IMARPE database shown a highly significant increase from 2015 onwards (P-value ∼ 0.000), passing from 1,257 records per year in 1996-2014 (SD = 255 records, min = 453, max = 1,783) to doubling this average from 2015 onwards (mean = 2,580, SD = 645, min = 1,599, max = 3,575). Thus, year 2015 was considered a break point of the time series metrics, so non-parametric pairwise comparisons were done to compare the average of the main metrics (total catch and the number of records, split by fishing gears) of the time series before and from 2015 onwards, performing the Wilcoxon Mann-Whitney test, and applying the Benjamini-Hochberg (FDR) correction for multiple comparisons where appropriate.

A second database of landings for *Myliobatis* spp. and *Mobula* spp., covering the period from 1996 to 2025, was provided by IMARPE. This database, organized as monthly landings by latitude-longitude coordinates and fishing gear, was obtained in response to a request for information made in accordance with Peruvian transparency regulations. The database was used to explore the fishing dynamics of these two groups of species. Coordinates were organized into grid cells measuring 0.2 by 0.2 degrees in latitude and longitude; consequently, the reported fishing volumes were aggregated by cell. To facilitate comparisons before and after the breakpoint, the data of each cell were summarized into two periods: Period 1) average annual landings from 1996 to 2014, and Period 2) average annual landings from 2015 to 2025. Cells with data on both periods were classified as potential traditional fishing zones, and the ones that only appeared in Period 2 were considered potential new fishing zones.

### Species prioritization based on a multi-criteria decision analysis (MCDA) framework

We developed a multi-criteria decision analysis (MCDA) framework to establish genetic research priority for the Peruvian batoid diversity. The MCDA was based on two-tiered hierarchical system. Tier 1 comprises the core score integrating ecological and anthropogenic risk parameters following Equation 1:

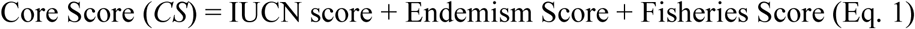

Where variables were parametrized based on the established scoring rubric presented in Table S1. Briefly, the *CS* spans from a minimum baseline score of 3 (defining species listed as LC, DD, or NE in the IUCN, with non-commercial value, and non-endemic to the ESPO) to a maximum possible score of 11 (assigned to species listed as CR in the IUCN, ESPO restricted, and under intense fisheries pressure). Following calculation, species were classified into three discrete baseline Tier 1 priority ranks: High Priority (9 ≤ *CS* ≤ 11); Medium Priority (6 ≤ CS ≤ 8); and Low Priority (3 ≤ CS ≤ 5).

Subsequently, a genetic modifier Tier 2 was applied to integrate evolutionary resilience into the scoring framework. We followed the global Elasmobranchii genetic diversity risk thresholds hosted on FishBase’s GDRI tool. The GDRI metric is based on the COI-*π*, using the statistical frameworks established by Petit-Marty et al. (2022) and Ferragut-Perrello et al. (2026). These frameworks define three genetic diversity levels: low (COI-*π* ≤ 0.0021), medium (0.0021 < COI-*π* ≤ 0.0025), and high (COI-*π* > 0.0025). This Tier 2 modifier prioritized localized COI-*π* derived strictly from samples collected in Peru, including the barcodes generated by the BATOseq-PE library alongside those from public repositories (BOLD and GenBank). The only exceptions were *Mo. birostris*, *Mo. munkiana*, *Mo. tarapacana*, and *Mo. thurstoni*, whose diversity metrics were sourced from Alfaro-Cordova et al. (2023). Sequences from outside the ESPO were excluded to avoid diversity inflation driven by regional structure. Due to the limited availability of Peruvian COI barcode sequences for batoids, a minimum threshold of five sequences (*n* ≥ 5) was established to calculate the *π* metrics while mitigating potential bias from small sample sizes.

For species exhibiting low local genetic diversity (COI-*π* ≤ 0.0021), their *CS* rank result was upgraded by one full priority category (e.g., shifting from Low Priority to Medium Priority). Conversely, if genetic diversity was deemed adequate (COI-*π* > 0.0021), the *CS* rank was retained. Finally, a data gap penalization, assigned to a “missing data” category, was applied for species represented by fewer than five COI barcodes in public repositories. Consequently, the final *CS* of these species was appended with a selective “High Priority (data gap)” flag to formally denote them as immediate targets for genetic research.

## 3. Results

### Primer performance, PCR amplification, and sequencing

Notably, the primer cocktail from Ivanova et al. (2007) yielded high-quality, species-specific sequences in some *My*. *chilensis* specimens where Fish1 primers (Ward et al. 2005) produced noisy chromatograms or amplified bacterial DNA. Similarly, universal Folmer primers successfully amplified and sequenced the guitarfishes *P. planiceps* and *Z. xyster*, whereas COI-1 oligos (FF2d/FR1d; Ivanova et al. 2007) optimized amplification for *R. velezi*. Across these three taxa, the Fish1 primer set failed during PCR or produced low-quality DNA sequences. Nevertheless, the Fish1 primers successfully amplified and identified distinct batoid samples including *A*. *laticeps*, *H. brevis*, *Mobula* spp., *Myliobatis* spp., and *U. chilensis*. The primer sets used for the amplification of each taxonomic group are detailed in Supplementary File S1.

A total of 75 samples were effectively amplified, representing a successful amplification rate of 100%. Of these, 74 amplicons (98.7%) yielded high-quality electropherograms after Sanger sequencing and were assigned to various chondrichthyan taxa. The remaining sample amplified bacterial DNA (i.e., *Vibrio* sp.) and consequently was excluded from further analyses. No signs of premature stop codons or frameshift mutations were detected in the newly generated sequences, indicating the total absence of NUMTs.

### DNA barcoding analyses and species diversity

The batoid sequences generated *in vitro* herein (*n =* 72) were submitted to the GenBank database under accession numbers PZ899860 to PZ899931. These novel COI barcodes successfully identified all samples to the species level, yielding a 99.69%-100% sequence identity match in the BOLD database. In total, 14 batoid species were identified, each assigned to a unique BIN (Table 1), distributed across 10 genera, 10 families, and four orders. When supplemented with the *in silico* assembled mitogenome of *N. martinezi*, our reference library encompasses a total of 15 species, clustered into 15 MOTUs and assigned to 15 BINs. This taxonomic representation accounts for 40.5% of the total marine batoid species richness in Peru (Campos-León et al. 2026).

The dominant commercial batoid group identified in our regional samples corresponded to eagle rays in the genus *Myliobatis* (*n* = 30, 41.7%), encompassing *My. chilensis* (*n* = 14), *My. peruviana* (*n* = 10), and *My. longirostris* (*n* = 6). This was followed by the Pacific guitarfish (*P. planiceps*) and the Peruvian diamond stingray (*H. brevis*) with 10 and nine specimens, respectively; whereas the Munk’s devil ray (*Mo. munkiana*) was represented by six samples. The spinytail devil ray (*Mo. mobular*), the Velez ray (*R. velezi*), and the blotched stingray (*U. chilensis*), were each represented by four specimens. The species with the lowest number of authenticated samples (*n* = 1) were the Pacific eagle ray (*A. laticeps*), the butterfly ray (*G. afuerae*), the smoothtail mobula (*Mo. thurstoni*), the giant electric ray (*N. entemedor*), and the witch guitarfish (*Z. xyster*). The specific GenBank accession numbers assigned to each analyzed specimen are provided in Table 1.

Finally, two processed samples labeled as “guitarfish *chinguirito*” were identified as the blue shark, *Prionace glauca* (GenBank accessions PZ901813 and PZ901814). Consequently, these mislabeled shark samples were excluded from our reference sequence library. Instead, these findings were integrated into the assessment of mislabeling and commercial names presented in Table S2.

### Genetic distances and barcoding gap

For the local BATOseq-PE library dataset, intraspecific distances ranged from 0 to 0.0055 (mean = 0.0003 ± 0.0009 SD), whereas within-genus interspecific distances were significantly higher, ranging from 0.0615 to 0.1211 (mean = 0.0856 ± 0.0221 SD). This clear separation established a well-defined barcoding gap (Fig. 3A).

**Fig. 3.**
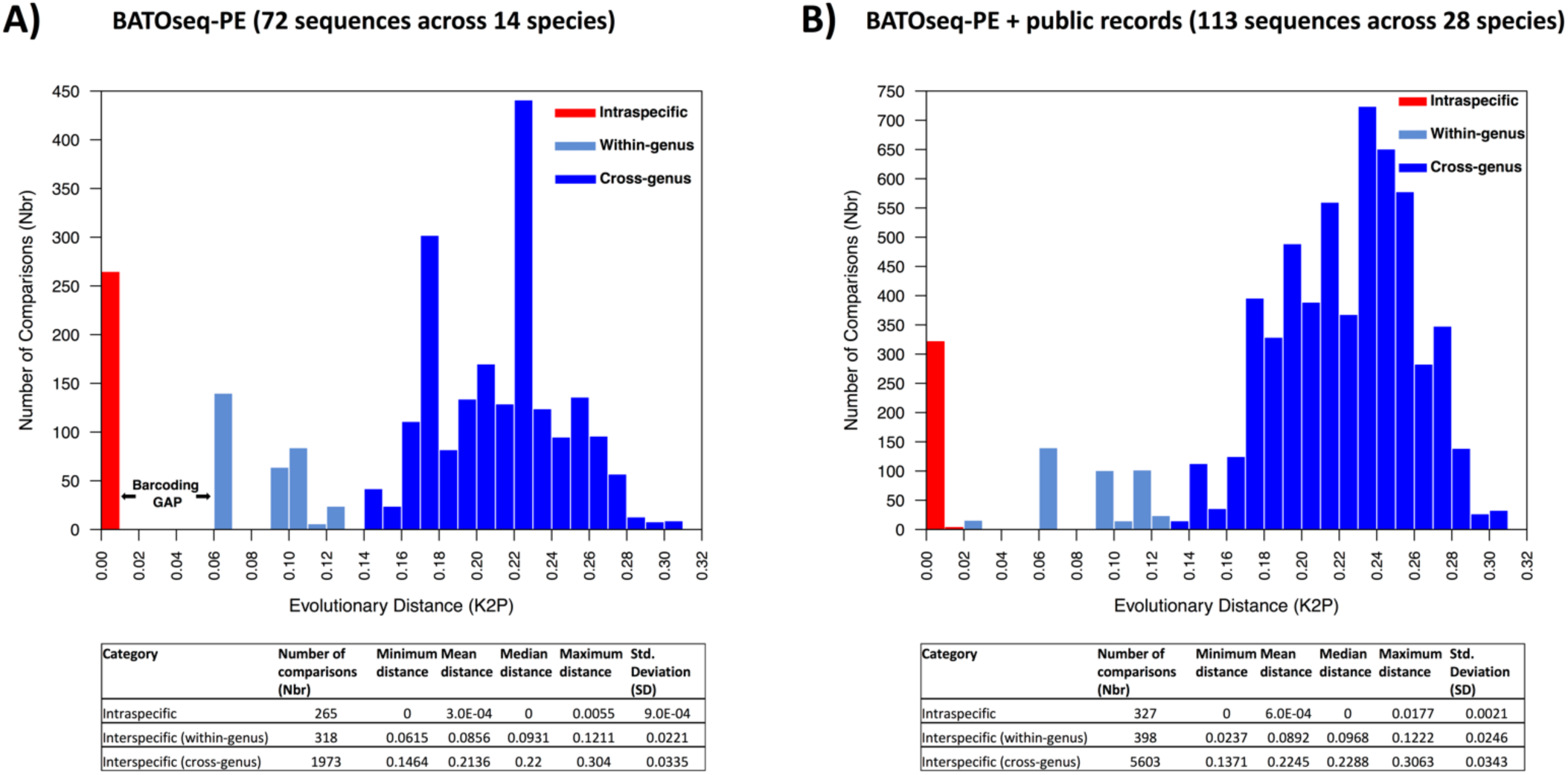
Distribution of pairwise genetic distances (Kimura 2-parameter model) for Peruvian marine rays. (A) Analysis of the local BATOseq-PE reference library, comprising 72 newly generated sequences across 14 species. (B) Analysis of the expanded global contextual dataset, combining local sequences with 113 public records across 28 species from Peru and abroad. Frequency distributions are shown for intraspecific (red), interspecific within-genus (light blue), and interspecific cross-genus (dark blue) genetic comparisons. Vertical bars represent the total number of pairwise comparisons (Nbr) sorted into bins of evolutionary distance (K2P). Standard summary metrics for each taxonomic comparison tier are provided in the embedded data tables below each histogram

Conversely, the inclusion of public records in the expanded dataset led to an increase in the maximum intraspecific divergence up to 0.0177 (mean = 0.0006 ± 0.0021 SD) (Fig. 3B). This high intraspecific value was driven entirely by a deep genetic split within *R. velezi*, resulting from the comparison between our four core Peruvian sequences (BSPE-57 to BSPE-60) and a public record from Costa Rica (BOLD ID RDFCA219-05). Concurrently, the minimum interspecific congeneric distance decreased to 0.0237 (mean = 0.0892 ± 0.0246 SD), resulting from the pairwise comparisons between *G. afuerae* sequences from Peru and *G. crebripunctata* from Mexico. The foreign *G. crebripunctata* sequences were included to assess the efficacy of COI barcoding identification for this putative Peruvian taxon. Consequently, due to the deep intra-species divergence within *R. velezi* and the low interspecific distance between *G. afuerae* and *G. crebripunctata*, the barcoding gap completely collapsed in the expanded global dataset.

### Taxonomic coverage, barcode sequence availability, and current status of the reference library for Peruvian batoids

As of July 31, 2026, following the completion of our global assessment of COI records for the 37 Peruvian marine batoid lineages, a total of 759 public records were verified across both the BOLD and GenBank databases (Fig. 4B, C). This dataset included 642 sequences (84.58%) for Myliobatiformes, 49 sequences (6.46%) for Torpediniformes, 40 records (5.27%) for Rhinopristiformes, and 28 records (3.69%) for Rajiformes.

**Fig. 4.**
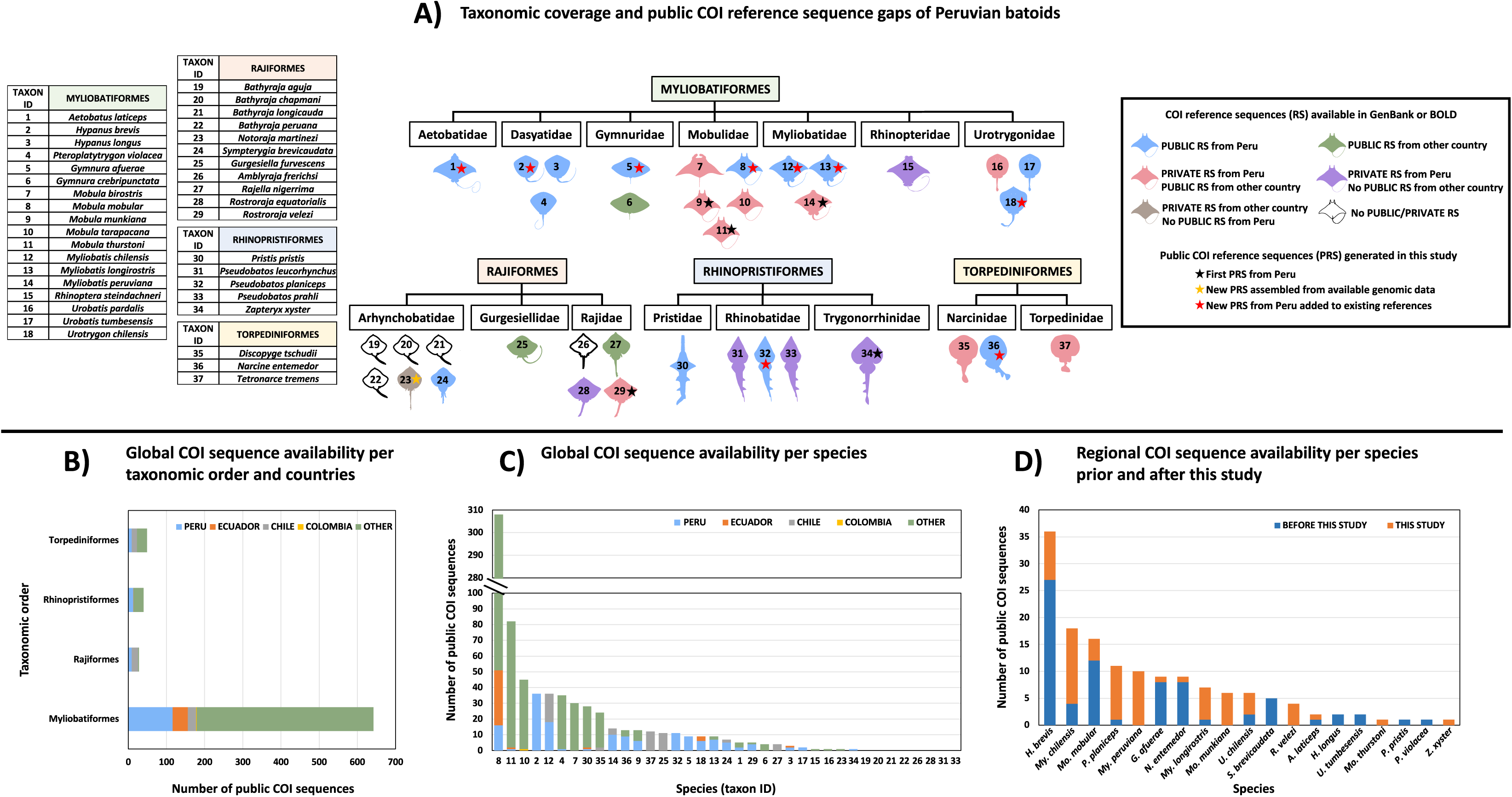
Taxonomic coverage, regional reference gaps, and global availability of the cytochrome c oxidase subunit I gene (COI) sequences across 37 Peruvian batoid species. (A) Schematic representation of taxonomic coverage across four batoid orders. Colored geometric icons indicate the baseline availability of COI reference sequences (RS) from Peru and other countries, while stars highlight public reference sequences (PRS) generated or updated in this study. Silhouettes for the genera *Aetobatus*, *Mobula*, *Myliobatis*, *Narcine*, *Pristis*, *Pteroplatytrygon*, *Rhinoptera*, *Urobatis*, and *Urotrygon* were retrieved from www.phylopic.org under the CC0 1.0 Universal Public Domain Dedication. (B) Cumulative abundance of public COI sequences partitioned by taxonomic order and country of origin (Peru, Ecuador, Chile, Colombia, and other nations). (C) Total global public COI sequence availability distributed by species. (D) Comparative ranking of regional COI sequence availability per species before (blue) and after (orange) the implementation of this study

When grouped by geographic provenance, non-ESPO nations account for the vast majority of the data, contributing 68.12% (517 sequences) of the global library. Within the ESPO region, Peru leads with the highest number of public references, representing 19.37% (147 sequences) of the total dataset, which includes the 72 novel sequences from the BATOseq-PE library alongside the newly assembled COI barcode for *N. martinezi*. This was followed by Chile (6.98%, 53 sequences), Ecuador (5.4%, 51 sequences), and Colombia (0.13%, 1 sequence) (Fig. 4B, C).

When grouped by species (Fig. 4C), three mobulid species accounted for the largest number of global sequences: *Mo. mobular* (*N* = 308; non-ESPO *n* = 257; Ecuador *n* = 35; Peru *n* = 16), *Mo. thurstoni* (*N* = 82; non-ESPO *n* = 80; Ecuador *n* = 1; Peru *n* = 1), and *Mo. tarapacana* (*N* = 45; non-ESPO *n* = 44; Colombia *n* = 1). Notably, no public COI reference sequences were available for *Mo. birostris* from the ESPO, despite Peru and Ecuador hosting the world’s largest population of this iconic, endangered species (Harty et al. 2022).

Prior to this study, the global COI reference library for Peruvian batoids was represented by 32 species, with five taxa lacking genetic barcodes entirely. Categorized by data availability status, this comprised 26 species with publicly available barcodes globally and six species whose sequences were retained entirely as private records. When categorized by geographical sampling origin, the library encompassed 28 species derived from Peruvian specimens (14 as public and 14 as private records) and four represented exclusively by foreign samples (three as public and one as private). Notably, nine of the 14 species with private Peruvian records already possess public barcodes of foreign origin.

At the regional level, before this research, the existing public reference data for Peruvian batoids was limited to 75 records comprising 14 species. By generating 72 new COI barcodes, the present research increased the public reference dataset by 96%, integrating novel public regional reference sequences for five batoid species: *Mo. munkiana*, *Mo. thurstoni*, *My. peruviana*, *Rostroraja velezi*, and *Z. xyster* (Fig. 4D). While the four former species already possessed public records from foreign localities, our new COI barcode for *Z. xyster* represents a novel global public record.

Although we detected a single public BOLD record for *Z. xyster* from Mexico (BOLD ID CBPM073-11), a reanalysis of its sequence placed it within the BIN BOLD:ACE7604, with 100% sequence identity to reference records of *A. laticeps*. Similarly, three Peruvian private records (FMCT011-18, FMCT749-19, and PMFSH677-20) labeled as *G. crebripunctata* exhibited a 100% identity match to our *G. afuerae* specimen BSPE-02 (GenBank PZ899861), which were assigned to the unique BIN BOLD:ADR4997. Another potentially misidentified entry correspond to the private regional record PMVTB106-20 (from Tumbes), which was labeled as *Urobatis halleri* but shared a 98.3% identity match and the same BIN (BOLD:AEE2435) with the *U. pardalis* record BMAR2855-22 (from Costa Rica). Regarding *A. laticeps,* while Vinueza et al. (2026) published COI sequences from specimens collected in four nations including Peru (GenBank PX289121–PX289193), these records lack voucher and country data, preventing the inclusion of their Peruvian barcodes in our comparative analyses.

Furthermore, the first complete mitogenome for *N. martinezi* assembled herein comprises its first global public COI barcode. Consequently, this study increases the global public reference library by two taxa, bringing the total from 26 to 28 species (Fig. 5). Notably, a total of five species still completely lack genetic barcodes for any marker (*Amblyraja frerichsi*, *Bathyraja aguja*, *B. chapmani*, *B. longicauda*, and *B. peruana*), rendering them currently unidentifiable via DNA barcoding.

**Fig. 5.**
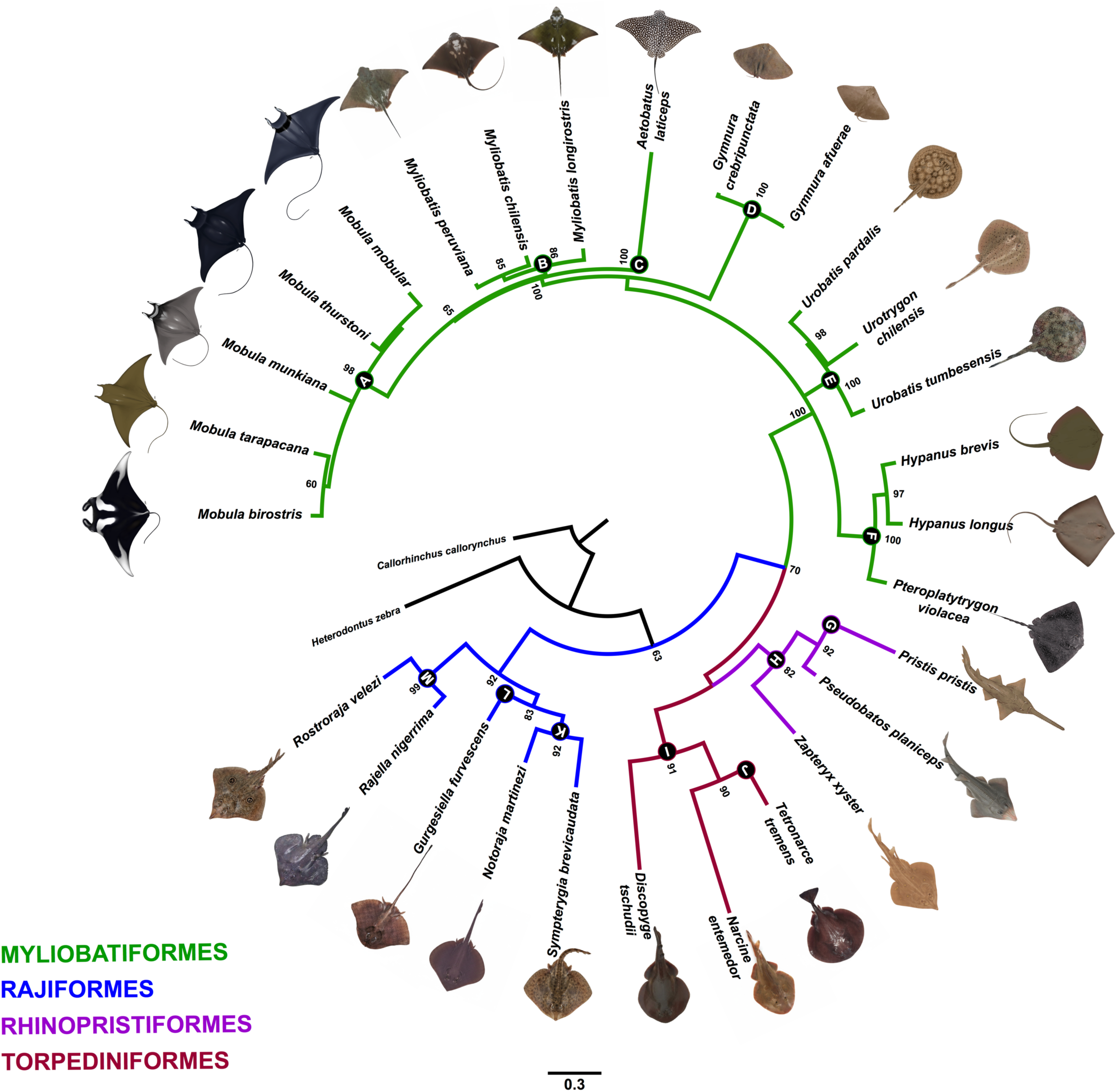
Maximum likelihood (ML) phylogenetic representation of public cytochrome c oxidase subunit I gene (COI) records for Peruvian batoids (28 species). The dataset includes 19 species directly generated from Peruvian records (14 from the BATOseq-PE library), the Costa Rican barcode for *Notoraja martinezi* assembled herein, and eight species integrated from public international barcodes. An additional nine species known to occur in Peru (bringing the regional total to 37 species) could not be included due to a complete lack of public records. This phylogenetic representation is presented solely to visually summarize taxonomic diversity rather than infer deep evolutionary relationships. Nodes display ultrafast bootstrap values (UFBoot2 ≥ 50). The topology is constrained to reflect the recovered monophyly of the genus *Gymnura*, which was artifactually placed within Myliobatidae without the inclusion of two auxiliary taxa: *Plesiobatis* and *Urolophus*. Major clades are color-coded by taxonomic order: Myliobatiformes (green), Rajiformes (blue), Rhinopristiformes (purple), and Torpediniformes (maroon). Capital letters on the tree nodes denote their respective family-level groupings: A (Mobulidae), B (Myliobatidae), C (Aetobatidae), D (Gymnuridae), E (Urotrygonidae), F (Dasyatidae), G (Pristidae), H (Rhinobatidae), I (Narcinidae), J (Torpedinidae), K (Arhynchobatidae), L (Gurgesiellidae), and M (Rajidae). Graphic illustrations for the five *Mobula* species were generously provided by the *Instituto Público de Investigación de Acuicultura y Pesca* (IPIAP) and AMAREA (Guerrero et al. 2026). Outgroups include the plownose chimaera, *Callorhinchus callorhynchus*, and the zebra bullhead shark, *Heterodontus zebra*

### Mitogenome assembly for *Notoraja martinezi*

The final circular consensus mitogenome sequence for *N. martinezi* was 16,700 bp in length (Fig. S1 from Supplementary Appendix S1). This nucleotide sequence was submitted to the GenBank database under accession number BK075715. Complete specimen metadata and assembly statistical metrics are provided in Table S3 from Supplementary Appendix S1.

Genetic characterization revealed that the mitogenome for *N. martinezi* exhibits a typical marine vertebrate configuration, containing 13 protein-coding genes, two ribosomal RNA genes, and 22 transfer RNA genes (see Table S4 in Supplementary Appendix S1).

Species identification analysis in BOLD revealed a 100% identity match between our query COI sequence from *N. martinezi* and three private records (MOP513-12, MOP737-12, MOP777-12) labeled as *N. martinezi* within BIN BOLD:ABY1401. The resulting phylogenetic tree file generated in BOLD is shown in Fig. S2 from Supplementary Appendix S1. These outputs confirmed that the specimen BioSample SAMN48911751 belongs to *N. martinezi*.

### Phylogenetic analysis

The ML phylogenetic reconstruction is presented in Fig. 6. The best-fit partitioning models according to ModelFinder were SYM+I+G4, F81+F+I, and GTR+F+G4 for the first, second, and third codon positions, respectively. The four primary batoid lineages (Myliobatiformes, Rajiformes, Rhinopristiformes, and Torpediniformes) were recovered as discrete monophyletic groups with moderate to strong nodal support. The order Rajiformes was recovered as the most basal sister clade to all other batomorphs, receiving strong nodal support (UFBoot2 = 98). The second major lineage was formed by the middle-tier sister relationship between the Rhinopristiformes and Torpediniformes (UFBoot2 = 50), whereas the Myliobatiformes was recovered as the most highly derived crown group (UFBoot2 = 86). These broad phylogenies align with the molecular consensus established by multilocus data (Aschliman et al. 2012) and complete genome surveys (Brownstein and Near 2026).

**Fig. 6.**
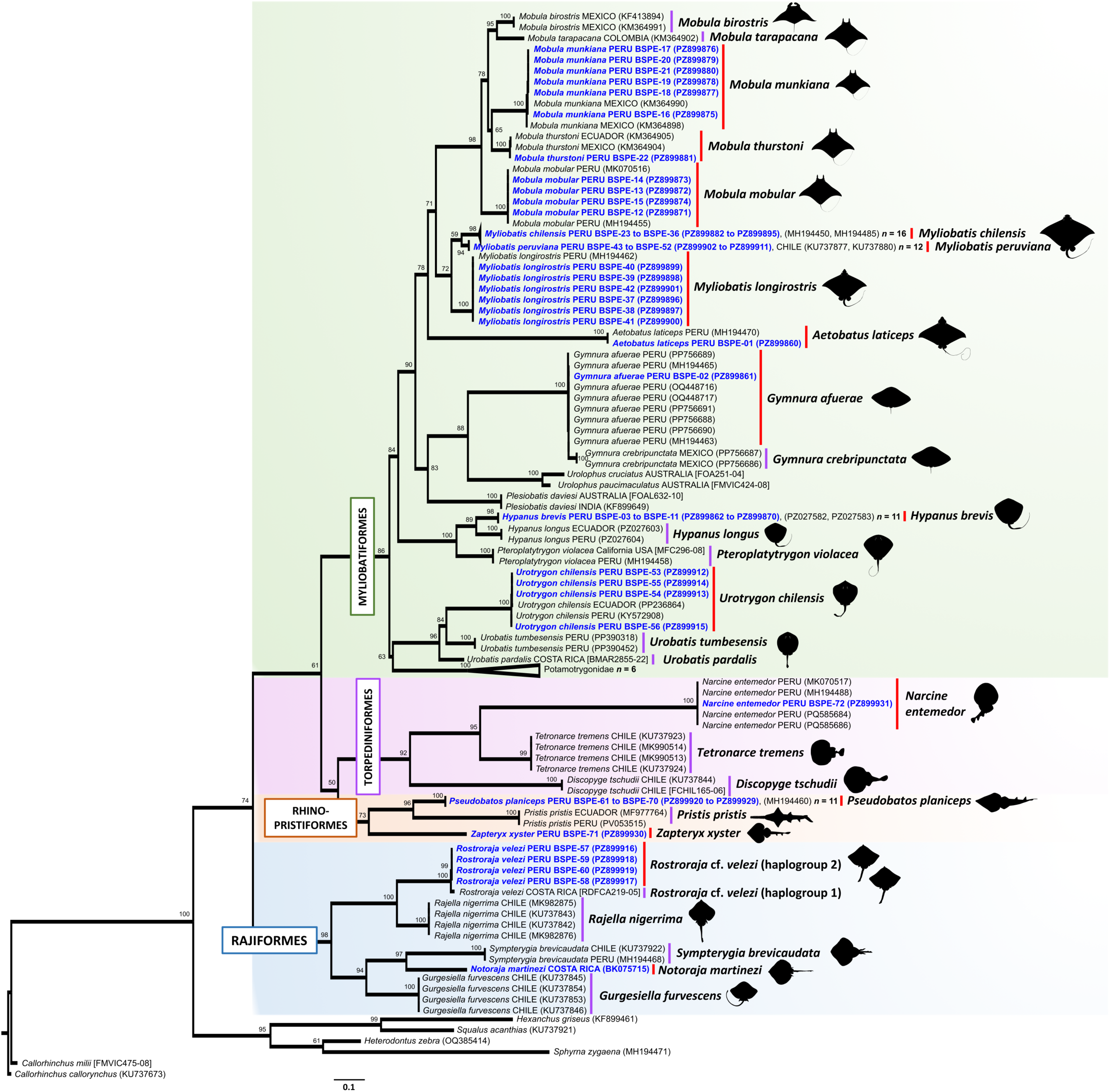
Maximum likelihood (ML) tree assessing the barcoding utility and phylogenetic relationships of batoid species based on cytochrome c oxidase subunit I gene (COI) sequences from the newly generated BATOseq-PE reference library alongside public reference records. Nodes display ultrafast bootstrap values (UFBoot2 ≥ 50). Sequences generated in this study are highlighted in bold blue, while reference sequences from public repositories are shown in regular black text. Molecular Operational Taxonomic Units (MOTUs) are indicated by red bars (originated from our novel barcodes) or violet bars (originated from reference sequences) on the right side of the tree. Sample information includes scientific name, sampling nation, BATOseqPE genetic voucher code (where applicable), GenBank accession number (in parentheses), and BOLD ID (in brackets). Clades containing more than 10 homologous sequences from target taxa were collapsed for visual clarity. Divergent reference taxa from genera *Plesiobatis, Urolophus*, and freshwater taxa (Potamotrygonidae) were selectively integrated to enhance topological stability. Outgroups include four sharks (*Heterodontus zebra*, *Hexanchus griseus*, *Sphyrna zygaena*, and *Squalus acanthias*) and two chimaeroid species (*Callorhinchus callorhynchus* and *C. milii*). Silhouettes for the genera *Aetobatus*, *Mobula*, *Myliobatis*, *Narcine*, *Pristis*, *Pteroplatytrygon*, *Rhinoptera*, *Urobatis*, and *Urotrygon* were retrieved from www.phylopic.org under the CC0 1.0 Universal Public Domain Dedication

Within the Myliobatiformes, family-level monophyly was consistently supported; however, during preliminary reconstructions, certain deep inter-family relationships remained unstable or contradictory to traditional systematic views. A phylogenetic artifact, where initially placed *Gymnura* deep within *Myliobatis* as the sister group of *My*. *longirostris*, was successfully resolved by including two auxiliary taxa: *Plesiobatis* and *Urolophus*. Because these two genera are more closely related to *Gymnura* (Aschliman et al. 2012; Naylor et al. 2012a), their inclusion caused *Gymnura* to cluster with them in a distinct monophyletic clade (UFBoot2 = 83) entirely outside *Myliobatis*.

Nevertheless, our barcoding analysis exhibited 100% accuracy in species discrimination for the 149 sequences included in the phylogenetic dataset. Excluding the auxiliary groups (Potamotrygonidae, *Plesiobatis*, and *Urolophus*), a total of 28 marine batoid morphospecies were successfully segregated into 29 distinct MOTUs, revealing cryptic divergence within *R. velezi*. Of these, 15 MOTUs originate from our novel COI sequences. These findings reinforce the robust utility of COI for the molecular identification of Peruvian marine batoids.

### Species delimitation analyses

*Rostroraja velezi* was the only taxon whose intraspecific pairwise genetic distance (K2P = 1.4%–1.6%, Fig. 7B) exceeded the threshold established herein (K2P ≥ 0.8) for species delimitation analysis. Consequently, we constructed a dataset comprising our four novel sequences (BSPE-57 to BSPE-60) alongside the only available public record for this species (BOLD ID RDFCA219-05, from Costa Rica). Additional congeneric barcodes included *R. alba* (*n* = 7) and *R. eglanteria* (*n*= 6), while rajiform allies comprised *B. spinicauda* (*n* = 5), *Raja asterias* (*n* = 7), *R. clavata* (*n* = 5), *R. maderensis* (*n* = 3), and *R. straeleni* (*n* = 5). The Chilean torpedo, *T. tremens* (*n* = 4), was used as an outgroup (see Table S5).

**Fig. 7.**
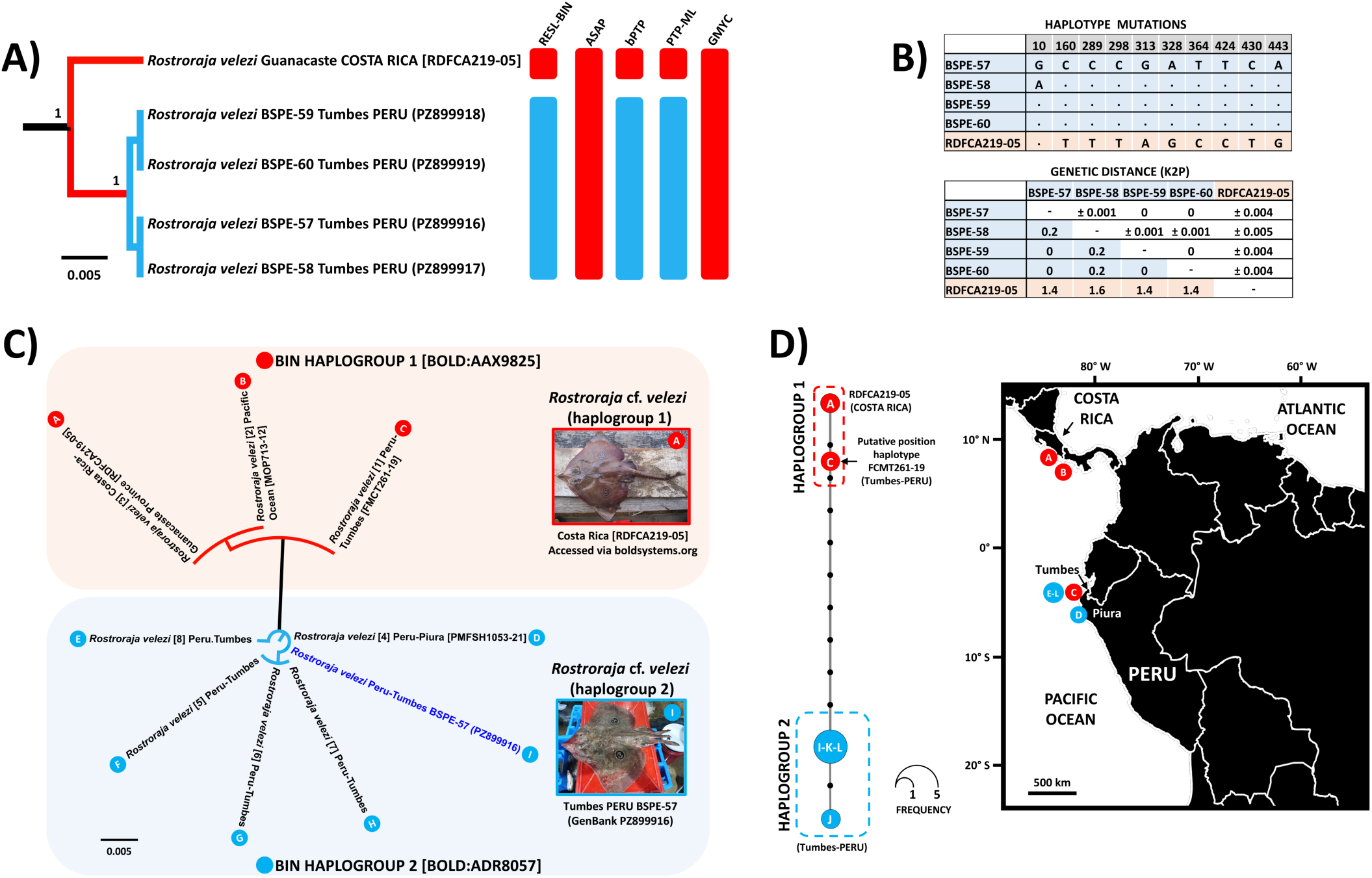
Species delimitation, genetic distance, and geographic distribution of *Rostroraja velezi* lineages based on partial cytochrome c oxidase subunit I gene (COI) sequences. (A) Species delimitation analysis. Ultrametric phylogenetic tree alongside multi-method species delimitation analyses (RESL-BIN, ASAP, bPTP, PTP-ML, and GMYC). Colored vertical bars indicate the molecular operational taxonomic units (MOTUs) recovered across methods. (B) Haplotype mutations and genetic distance matrices. The upper table shows the diagnostic nucleotide mutations across variable alignment positions identifying the different haplotypes of *Rostroraja velezi* from Costa Rica and Peru. The lower table displays the percentage pairwise genetic distances (Kimura 2-parameter model). (C) Distance-based identification tree natively generated within the BOLD Systems platform using the query COI sequence from our specimen BSPE-57 (highlighted in blue text). Sequences assigned to Barcode Index Number (BIN) BOLD:AAX9825 are indicated by red circles (labeled A to C), while those belonging to BIN BOLD:ADR8057 are denoted by blue circles (labeled D to I); corresponding codes are mapped onto the geographic distribution panel. Except for the public reference sequence from Costa Rica [RDFCA219-05], all other BOLD sequences (labeled B to H) represent private records. (D) Haplotype network and geographic distribution. Median-joining network constructed using our novel sequences from Tumbes (labeled I: BSPE-57; J: BSPE-58; K: BSPE-59; and L: BSPE-60) alongside the public Costa Rican reference sequence (labeled A). The red circle labeled C indicates the putative network position inferred for the private sample FCMT261-19 (Tumbes-Peru). Sampling localities on the map are color-coded to correspond with recovered molecular lineages: red markers represent haplogroup 1 (BIN BOLD:AAX9825; Costa Rica, Pacific Ocean, and Tumbes, Peru), while blue markers denote haplogroup 2 (BIN BOLD:ADR8057; Tumbes and Piura, Peru)

BOLD’s RESL algorithm assigned *R. velezi* sequences into two distinct BINs. The first BIN BOLD:AAX9825 (haplogroup 1) clustered three BOLD sequences from the Eastern Pacific: the public record from Costa Rica (*n* = 1) and two private records from the Pacific Ocean (*n* = 1, MOP713-12), and Tumbes, Peru (*n* = 1, FMCT261-19). Conversely, BIN BOLD:ADR8057 (haplogroup 2) comprised nine Peruvian sequences, including our four core specimens from Tumbes alongside five regional private records from BOLD: four from Tumbes (FMCT258-19, FMCT259-19, FMCT260-19, FMCT370-19) and one from Piura (PMFSH1053-21) (Table 1, Fig. 7).

Two additional tree-based algorithms, PTP-ML and bPTP, strongly supported the existence of two distinct MOTUs within the *R. velezi* clade, mirroring the dual-BIN structure (Fig. 7A). These methods delimited the Costa Rican specimen as an independent evolutionary unit (support = 0.97) and our four Peruvian individuals from Tumbes as a distinct lineage (support = 0.96), further supported by a low model transition value (0.03) at their shared ancestral node. In contrast, the more conservative global parameters of the ASAP and GMYC algorithms lumped all sequences into a single MOTU.

The phylogenetic tree generated via the BOLD ID engine (Fig. 7C) and the haplotype network analysis (Fig. 7D) confirmed this dual structure. The three sequences from the Eastern Pacific (FMCT261-19, MOP713-12, and RDFCA219-05), belonging to BIN BOLD:AAX9825 (haplogroup 1), were recovered in a unique clade. Conversely, our four core specimens from Tumbes clustered into a well-defined group within BIN BOLD:ADR8057 (haplogroup 2), displaying 100% sequence identity with specific regional private records. Specifically, three core specimens (BSPE-57, BSPE-59, and BSPE-60) exhibited a 100% match with the private record from Piura (PMFSH1053-21), whereas BSPE-58 matched perfectly with a private Tumbes record (FMCT370-19).

In the haplotype network, haplogroup 1, which comprises the public sequence from Costa Rica and putatively two associated private records (FMCT261-19 and MOP713-12), was separated by up to 10 mutational steps from our four novel Peruvian sequences (K2P: 1.4% to 1.6%; Fig. 7B, D). The maximum genetic distance of 1.6% detected between the Costa Rican haplotype and our Peruvian specimen BSPE-58 (Fig. 7B) far exceeds the maximum intraspecific variation reported for numerous batoid lineages (Crobe et al. 2021; Petean et al. 2024). The private record MOP713-12 (origin labeled as “Pacific Ocean”) likely originates from Central America, as the “MOP” dataset belongs to the Smithsonian Institution’s regional collection associated with that region. Conversely, haplogroup 2 is composed exclusively of Peruvian samples, clustering our four novel barcodes from Tumbes with the putative anchoring of five regional private barcodes (four from Tumbes and one from Piura).

Most species delimitation methods conducted herein strongly support the existence of an evolutionary cryptic divergence within *R. velezi*. Crucially, the co-occurrence of both lineages in Tumbes provides additional biological evidence of reproductive isolation rather than geographic variation. Therefore, we recognize that the two MOTUs revealed in this research represent two distinct unconfirmed candidate species, designated as *Rostroraja* cf. *velezi* (haplogroup 1) and *Rostroraja* cf. *velezi* (haplogroup 2), pending ancient DNA validation of the historical holotype.

### Phylogeographic and population structure assessments

Based on the species selection criteria outlined in the “*Materials and methods*” section, four batoid species (*Mo. mobular*, *Mo. munkiana*, *Sympterygia brevicaudata*, and *U. chilensis*) were selected for phylogeographic assessments. Notably, a relatively larger sample size across distinct populations was available only for *Mo. mobular* (Ecuador: *n* = 23; Mexico: *n* = 21; Peru: *n* = 13), enabling a more detailed assessment of both its phylogeography and population structure. On the other hand, *Mo. munkiana*, *S. brevicaudata*, and *U. chilensis* exhibited limited sample sizes, ranging from merely 3 to 6 individuals per population. Consequently, the phylogeographic evaluations for these three taxa were restricted to haplotype networks, and their outputs must be considered preliminary and interpreted with caution.

The median-joining haplotype network for both *Mo. mobular* (Fig. 8A) and *Mo. munkiana* (Fig. 8B) revealed a pattern of high connectivity across the Eastern Pacific Ocean. Although the molecular diversity metrics for *Mo. munkiana* may be constrained by small sample sizes (Mexico: *n* = 5, Peru: *n* = 6), the detection of a shared haplotype between Mexico and Peru is consistent with historical or ongoing genetic connectivity. For *Mo. mobular*, the phylogeographic analysis revealed a southward latitudinal decrease in genetic diversity. The Mexican population exhibited the highest levels of molecular variation (*H_d_* = 0.4238, *π* = 0.000870), followed by intermediate values in Ecuador (*H_d_* = 0.3913, *π* = 0.000783), and the lowest baseline records in Peru (*H_d_* = 0.1538, *π* = 0.000282). The corresponding haplotype network displayed a classic star-like topology, dominated by a single, high-frequency central ancestral node shared among all sampled nations, with unique, low-frequency satellite haplotypes separated by a single mutational step. These derivative haplotypes branched off predominantly from Mexican and Ecuadorian lineages, while the Peruvian populations were almost entirely fixed within the shared ancestral haplotype.

**Fig. 8.**
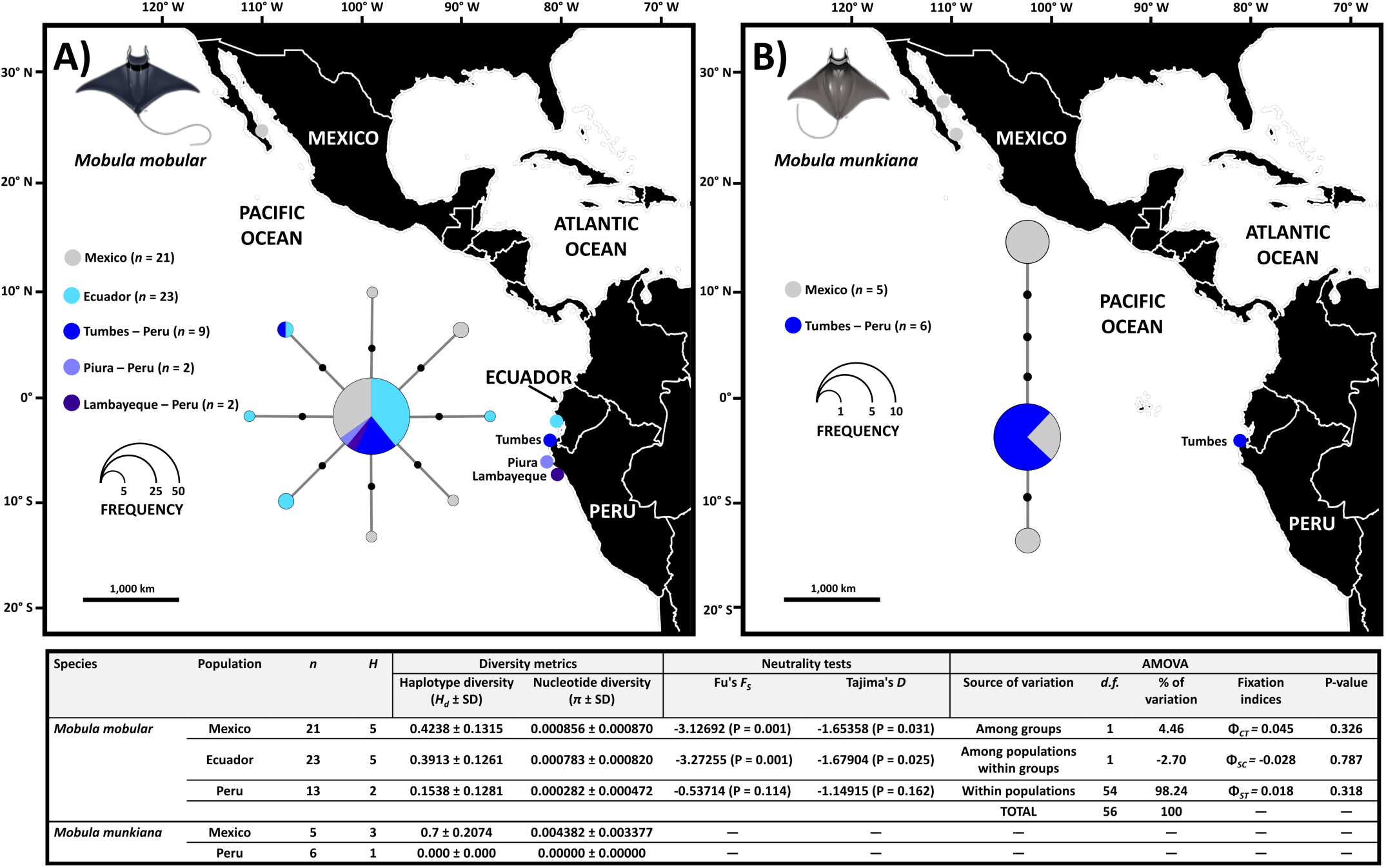
Haplotype network and spatial genetic diversity of *Mobula mobular* and *Mobula munkiana* in the Eastern Tropical Pacific. (A) Median-joining haplotype network for *Mo. mobular* across Mexico, Ecuador, and northern Peru (Tumbes, Piura, and Lambayeque). (B) Median-joining haplotype network for *Mo. munkiana* across Mexico and Tumbes, Peru. Circle sizes are proportional to total haplotype frequencies, and transverse black dots along the connecting lines indicate single mutational steps. Colored pie charts represent the relative proportion of individuals from each geographic region contributing to that specific haplotype. The table in the bottom panel displays population genetic parameters, neutrality tests for *Mo. mobular* and *Mo. munkiana*, and Analysis of Molecular Variance (AMOVA) for *Mo. mobular*, based on cytochrome c oxidase subunit I gene (COI) barcodes. The bottom-right section details the hierarchical AMOVA results for *Mo. mobular*, partitioning genetic variance among regional groups from the Northern population (Mexico) vs. the Southern populations (Ecuador and Peru), among populations within groups, and within populations. Graphic illustrations for *Mo. mobular* and *Mo. munkiana* were generously provided by the *Instituto Público de Investigación de Acuicultura y Pesca* (IPIAP) and AMAREA (Guerrero et al. 2026)

Neutrality tests for both the Mexican and Ecuadorian populations of *Mo. mobular* yielded significantly negative values for *F_S_* (see embedded lower panel in Fig. 8), denoting a significant excess of rare alleles and a signature of demographic expansion following a historical bottleneck. Conversely, the non-significant negative values observed in Peru likely reflect a combination of lower regional sample size and localized genetic fixation. Finally, the hierarchical AMOVA test, which evaluated the regional groups of North (Mexico) versus South (Ecuador and Peru), revealed a complete absence of regional genetic structure. Instead, the vast majority of molecular variance (98.24%) occurred within populations, yielding a low and non-significant global pairwise fixation index (Φ*_ST_* = 0.018, P = 0.318).

S*ympterygia brevicaudata* exhibits a low-to-moderate *H_d_* and low *π*, characterized by a dominant northern Peruvian core and a uniform, fixed Chilean haplotype (Fig. 9A). The high standard deviations surrounding the Peruvian diversity metrics are a mathematical consequence of small sample sizes rather than biological instability. Similarly, *U. chilensis* displays a star-like network topology defined by high *H_d_* and low *π* across both Peruvian and Ecuadorian groups (Fig. 9B). In both species, the combination of moderate/high *H_d_* and low *π* is a classic signature of historically recent population expansion following a bottleneck or founder event (Schlichta et al. 2025). The shallow mutational divergence across both networks indicates sustained historical connectivity, suggesting that the Humboldt Current system and its northern equatorial transition zones may act as an effective corridor facilitating gene flow.

**Fig. 9.**
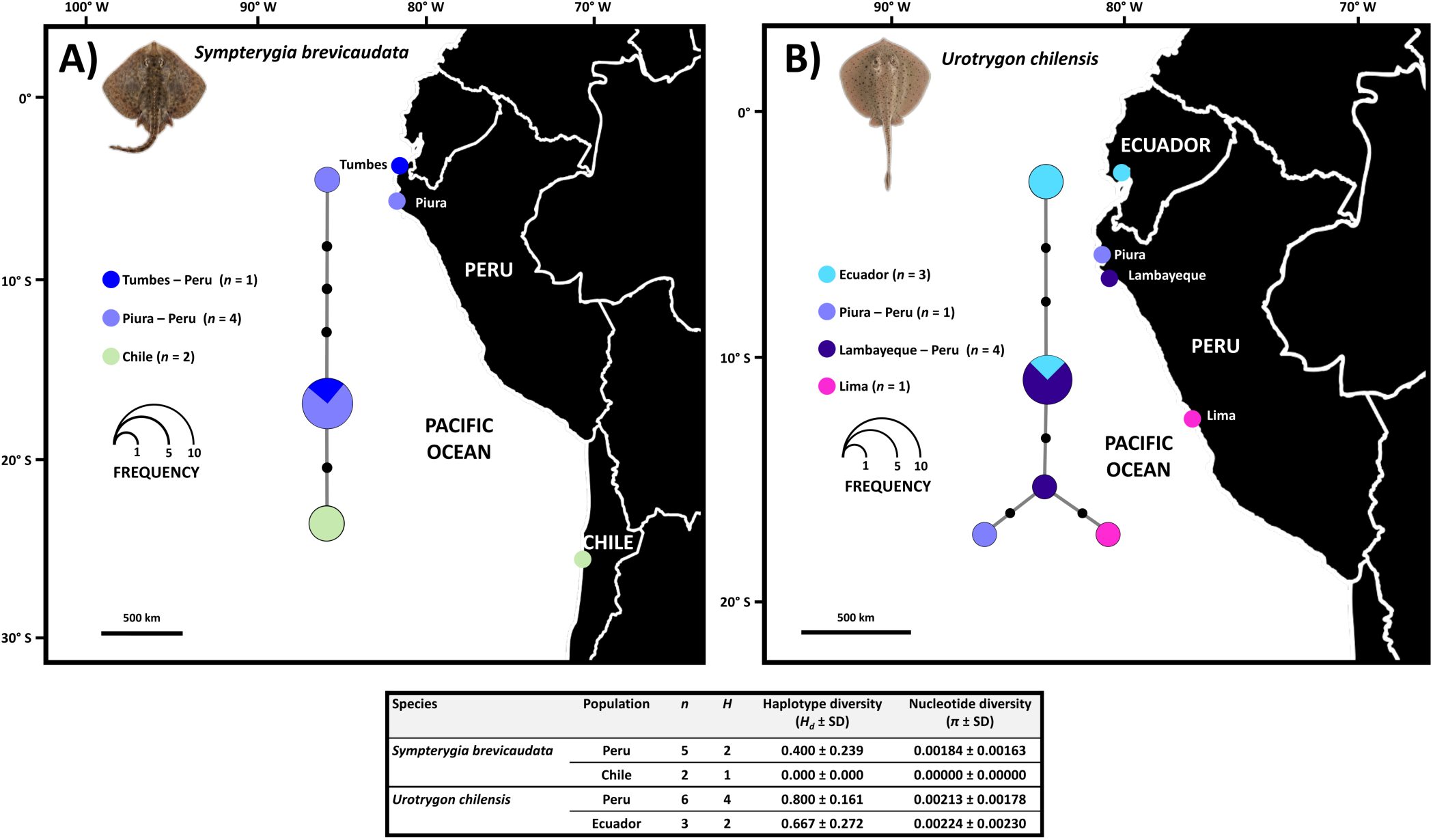
Phylogeography, haplotype networks, and genetic diversity of *Sympterygia brevicaudata* and *Urotrygon chilensis* along the Eastern South Pacific. (A) Phylogeography of *S. brevicaudata*, displaying the geographic sampling locations and a median-joining haplotype network based on cytochrome c oxidase subunit I gene (COI) sequences from Peru (Tumbes and Piura) and Chile. (B) Phylogeography of *U. chilensis*, displaying the geographic sampling locations and a median-joining haplotype network from Ecuador and Peru (Piura, Lambayeque, and Lima). In both networks, circle sizes are proportional to haplotype frequency, and black dots along the connecting lines represent mutational steps between haplotypes. The table below shows genetic diversity metrics for *S. brevicaudata* and *U. chilensis*, including sample size (*n*), number of haplotypes (*H*), haplotype diversity (*H_d_*), and nucleotide diversity (*π*)

### Current knowledge of Peruvian marine batoid diversity

Earlier comprehensive checklists of Peruvian marine fishes by Chirichigno and Cornejo (2001) and chondrichthyans by Cornejo et al. (2015) recognized 46 and 43 nominal batoid species, respectively. Following stringent taxonomic filtration, revised geographical distribution records, verified genetic entries, and the addition of new species (i.e., *B. chapmani*) and novel regional records (i.e., *A. frerichsi* and *N. martinezi*), the most recent checklist of Peruvian chondrichthyan diversity recognized 37 marine batoid species (Campos-León et al. 2026). These updated taxa are grouped across 22 genera, 15 families, and four orders, showcasing a comprehensive baseline for the region. Most notably, this consolidation includes the formal lumping of six formerly recognized *Urotrygon* species (*U. aspidura*, *U. caudispinosus*, *U. chilensis*, *U. munda*, *U. peruanus*, and *U. serrula*) into a single taxon, *U. chilensis* (Ehemann et al. 2024a).

Twelve species (32.4%) are endemic to the ESPO, among which *B. chapmani* currently represents the sole micro-endemic species exclusive to Peru (Ebert et al. 2022). Notably, 28 species have already been genetically barcoded in Peruvian waters, as revealed by our screening of both public and private regional molecular records (Fig. 4). Although 10 species have not yet been verified by DNA barcoding locally, the taxonomic validity of most deep-sea or conservative taxa has been thoroughly established through rigorous morphological assessments; e.g., *A. frerichsi*, *B. aguja*, *B. chapmani*, *B. longicauda*, *B. peruana*, and *N. martinezi* (Ebert et al. 2022; Zavalaga et al. 2024).

#### IUCN conservation status and national regulatory framework of Peruvian batoids

According to the conservation status assessments conducted by the IUCN, the vast majority of Peruvian batoid taxa (20 species, 54.0%) are classified within threatened categories: four species (10.8%) are Critically Endangered (CR), one (2.7%) is Endangered (EN), and 15 (40.6%) are Vulnerable (VU). Conversely, the remaining 17 species comprise four (10.8%) listed as Near Threatened (NT), 10 (27.0%) as Least Concern (LC), and three (8.1%) as Not Evaluated (NE) (Fig. S3). It should be noted that the NE category comprises three ESPO-restricted taxa: *B. chapmani*, *G. afuerae*, and *H. brevis*. While the former species is a relatively recent discovery (Ebert et al. 2022), the latter two taxa have been recently split from their Eastern North Pacific geminates (*G. crebripunctata* and *H. dipterurus*, respectively) (Ehemann et al. 2024b; Marín et al. 2026) and are currently awaiting official conservation assessments.

When segregated by taxonomic order, the composition of IUCN categories varies significantly. Within the Myliobatiformes (*n* = 18), three species each (16.7%) are CR and NT, one (5.6%) is EN, seven (38.8%) are VU, and two species each (11.1%) are LC and NE. Within Rajiformes (*n* = 11), three species (27.3%) are classified as VU, one (9.1%) as NT, six (54.5%) as LC, and one (9.1%) as NE. The Rhinopristiformes (*n* = 5) exhibit a high proportion of extinction risk, with one species (20.0%) listed as CR and four (80.0%) as VU. Finally, within the Torpediniformes (*n* = 3), one species (33.3%) is designated as VU, while two species (66.7%) are classified as LC.

The current regulatory framework for Peruvian marine batoids is heavily fragmented and inherently weak, lacking species-specific conservation and management measures (e.g., minimum landing size regulations and seasonal reproductive closures). Instead, management relies exclusively on a gillnet mesh size restriction of 200-330 mm (R.M. N° 209-2001-PE; PRODUCE 2001), which loosely groups eagle rays (*Myliobatis* spp.) and the Pacific cownose ray (*R. steindachneri*). The National Plan of Action for the Conservation and Management of Sharks, Rays, and Allied Species (PAN-Tiburón), approved via D.S. N° 002-2014-PRODUCE (PRODUCE 2014), establishes the broad guidelines for national chondrichthyan management. Explicit *de jure* capture bans are strictly limited to the oceanic manta ray, *Mo. birostris* (R.M. N° 441-2015-PRODUCE; PRODUCE 2016) and the largetooth sawfish, *P. pristis* (R.M. N° 056-2020-PRODUCE; PRODUCE 2020).

Conservation and management measures for mobulids, which are the most developed within this framework, began with a strict ban on the harvest and trade of *Mo. birostris* (R.M. N° 441-2015-PRODUCE; PRODUCE 2016). This regulation was subsequently amended by R.M. N° 000362-2024-PRODUCE (PRODUCE 2024a) to expand its scope into a *de jure* prohibition covering the entire genus *Mobula*. Despite these explicit bans, their current *de facto* commercial availability (as corroborated during our fieldwork) is sustained by a rolling legal loophole. First established under R.M. N° 00411-2024-PRODUCE (PRODUCE 2024b) and subsequently revised on several occasions, this loophole was most recently re-issued in substantially similar terms under R.M. N° 00152-2026-PRODUCE (PRODUCE 2026). This resolution allows commercial landings of non-*birostris* mobulids under bycatch status, through December 2026.

### Exploratory description of batoid fishery data and trends

IMARPE database reported up to 16 species categories, divided into i) 12 single species identified by their scientific names and ii) four groups of congeneric species identified at the genus level (Fig. 10A). In the recent history of the batoid fishery (2015-2024), 1,289 tons·year^-1^ were reported as landed, with two myliobatiform genera at the top of the landing list: the *Myliobatis* spp. and *Mobula* spp. (Fig. 11). The first group accounts for 60.1% of the total recorded batoid landing volumes for the period 2015-2024 and may comprise three distinct species: *My. chilensis, My. longirostris,* and *My. peruviana*. However, it was not possible to determine whether all three species are included or which species predominates in the records. The second group, which may mask information from up to five species (*Mo. birostris, Mo. mobular, Mo. munkiana, Mo. tarapacana,* and *Mo. thurstoni*), represented 29.4% of all 2015-2024 volumes.

**Fig. 10.**
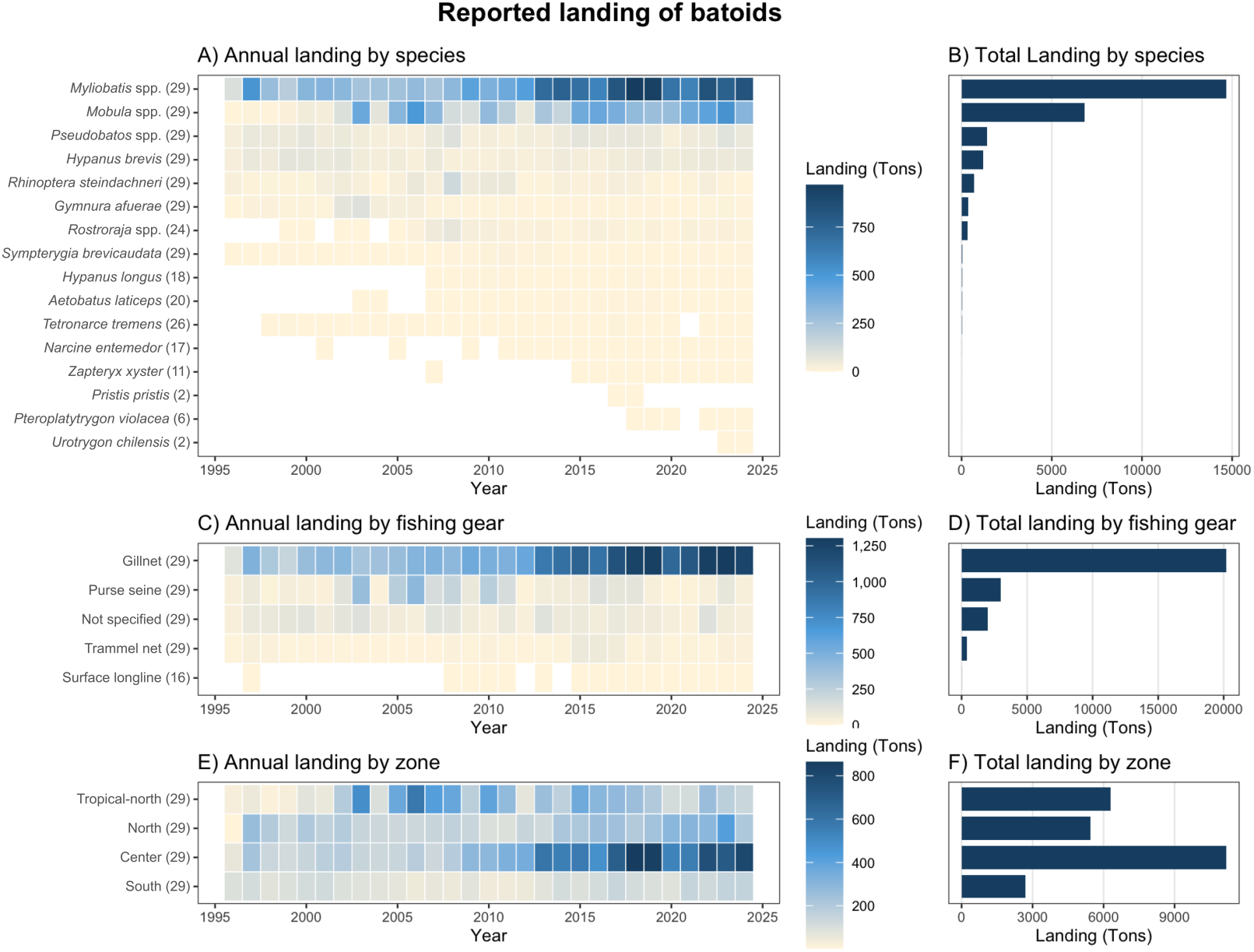
Annual and cumulative catch volumes of batoids from 1995 to 2024, categorized by species (A and B), fishing gear (C and D), and latitudinal zones (E and F)

**Fig. 11.**
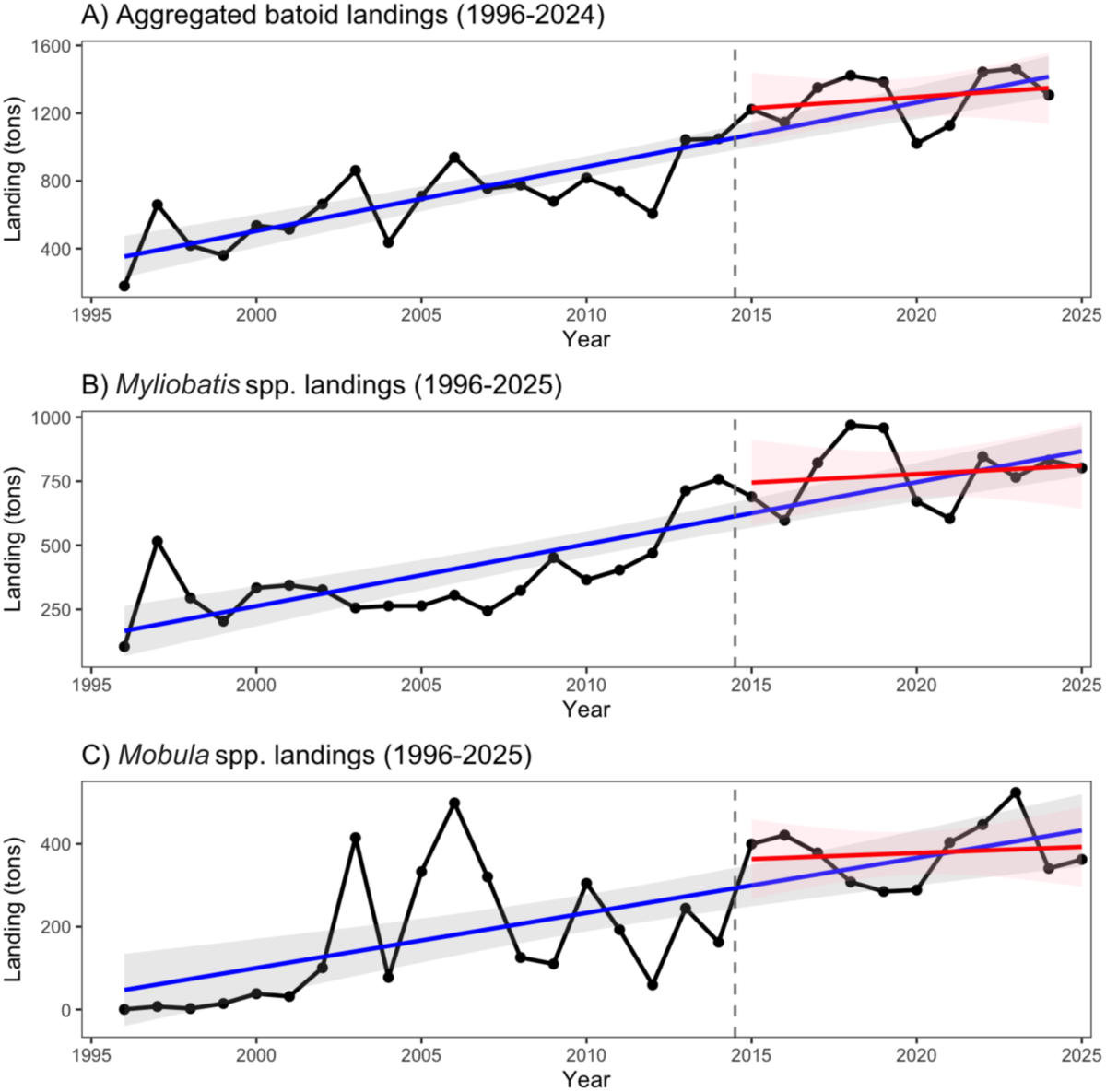
Time series of the landings of the aggregated batoid species (A), *Myliobatis* spp. (B), and *Mobula* spp. (C). Grey dashed line represents the division between Period 1 (1995-2014) and Period 2 (2015 onwards)

Notably, 88.9% of batoid landings were recorded with gillnets (Fig. 10C, D), while 52.9% were reported landed in coastal areas from Central Peru (Fig. 10F), where the magnitude of the landings has been shown to increase in recent years (Fig. 10E). Other relevant areas are the North and the Tropical North, accounting for 20.6% and 17.0%, respectively. A detailed analysis of the historical trends, fishing gear contributions, and expansion associated with the *Myliobatis* and *Mobula* fisheries is provided in Supplementary Appendix S2; Fig. S4, and Fig. S5.

After *Myliobatis* and *Mobula*, *Pseudobatos* spp. (mostly *P. planiceps*) and *H. brevis* are the main regional taxa contributing consistently to Peruvian batoid fisheries. The guitarfishes represent the third largest group in cumulative landing biomass, exhibiting continuous annual landings throughout the entire study frame (1995–2025). *Hypanus brevis*, the fourth most relevant species by volume, is characterized by an exceptionally persistent historical presence with stable, uninterrupted annual records across the three decades (Fig. 10A, B; see Marín et al. 2026). Other secondary taxa, such as *R. steindachneri* and *G. afuerae*, displayed lower but steady volumes across historical landings. Conversely, structurally rare or highly depleted taxa, including *P. violacea*, *U. chilensis*, and the CR *P. pristis* (which has been strictly banned since 2020) exhibited marginal, heavily fragmented landings restricted to very few years, likely confirming their rare, incidental nature within artisanal fisheries.

### MCDA research priority classification

By integrating data from IUCN extinction risk profiles, regional endemism records, and landing statistics (Tier 1 metrics) with molecular diversity values (Tier 2 metrics), we constructed the first comprehensive research priority framework for Peruvian batoids (Table 2). A fully automated Excel file containing the MCDA framework is presented as Appendix S3. Baseline demographic and fisheries indicators revealed critical vulnerabilities across the 37 evaluated species: over half (54.0%, *n* = 20) are threatened with extinction according to the IUCN Red List, 32.4% (*n* = 12) are regional endemics, and 29.7.1% (*n* = 11) face intense fisheries pressure.

**Table 2.**
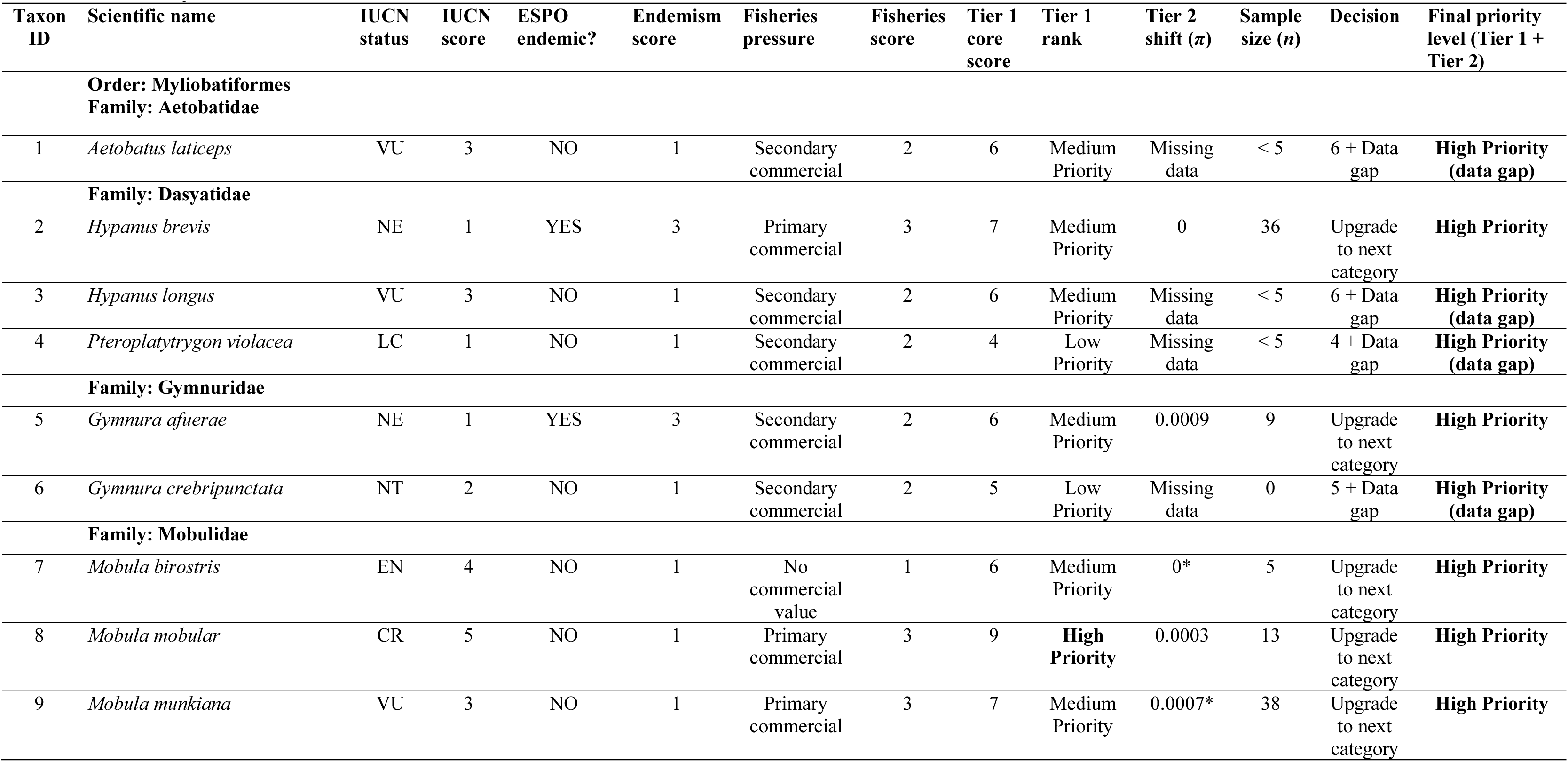
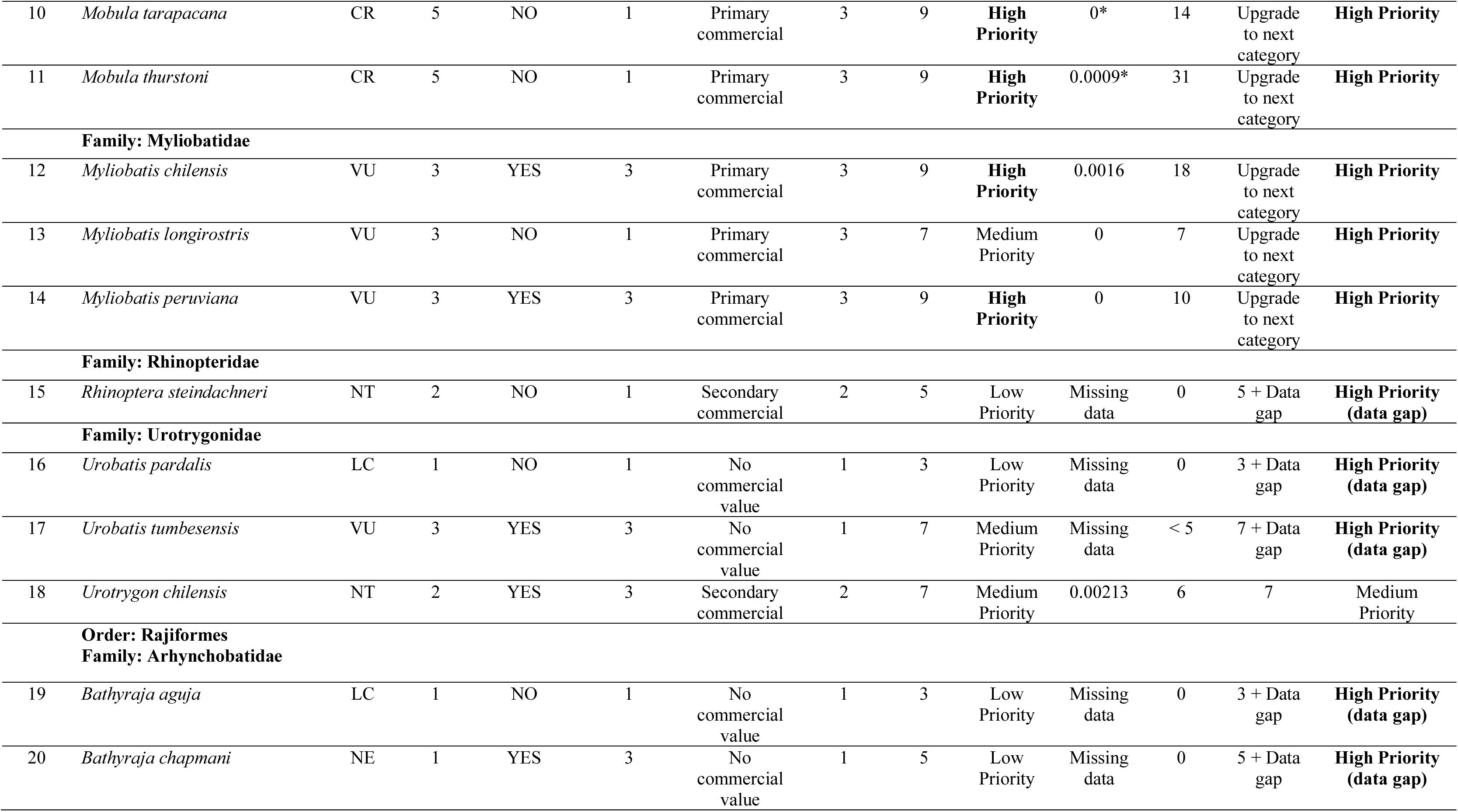
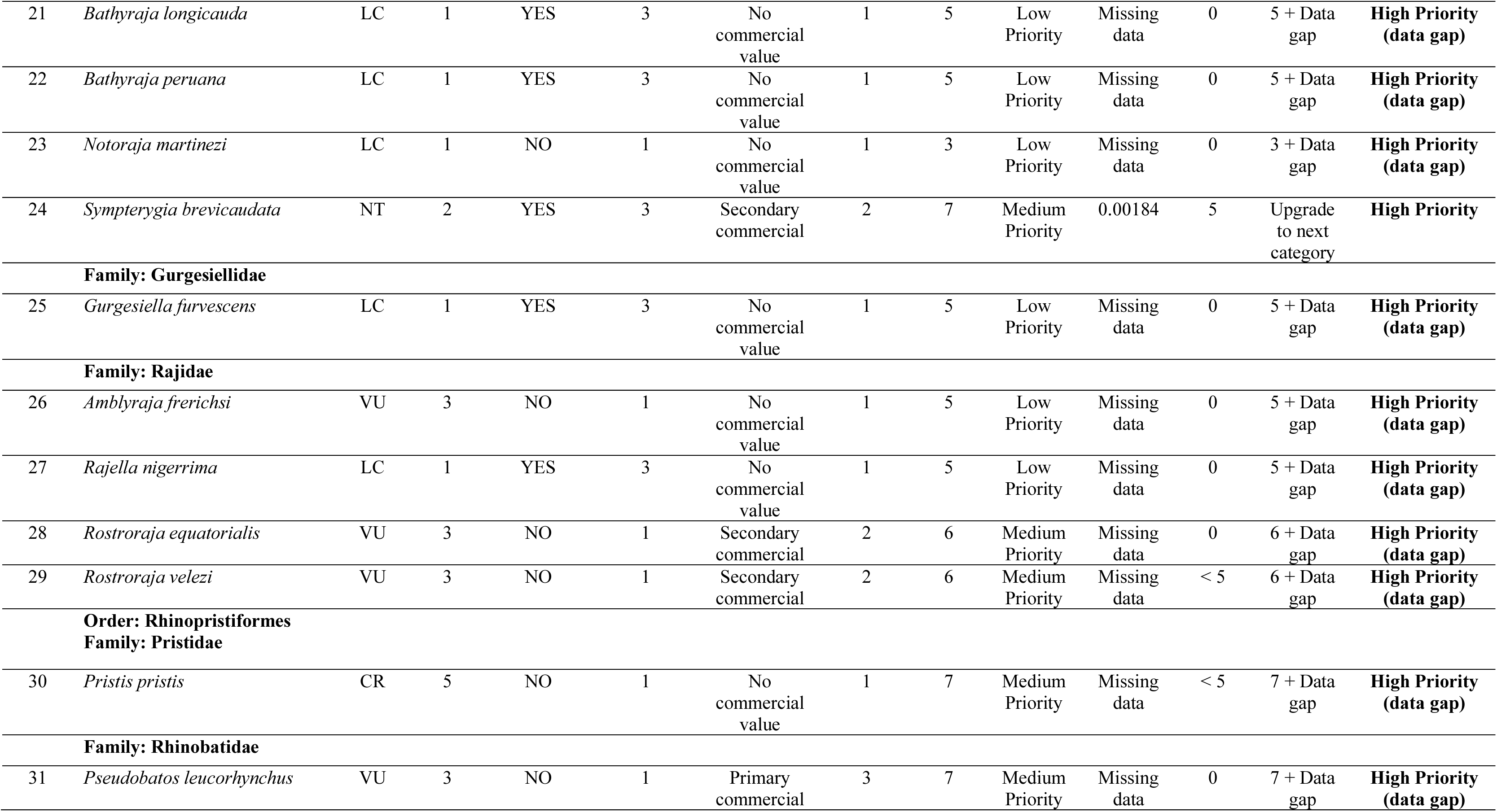
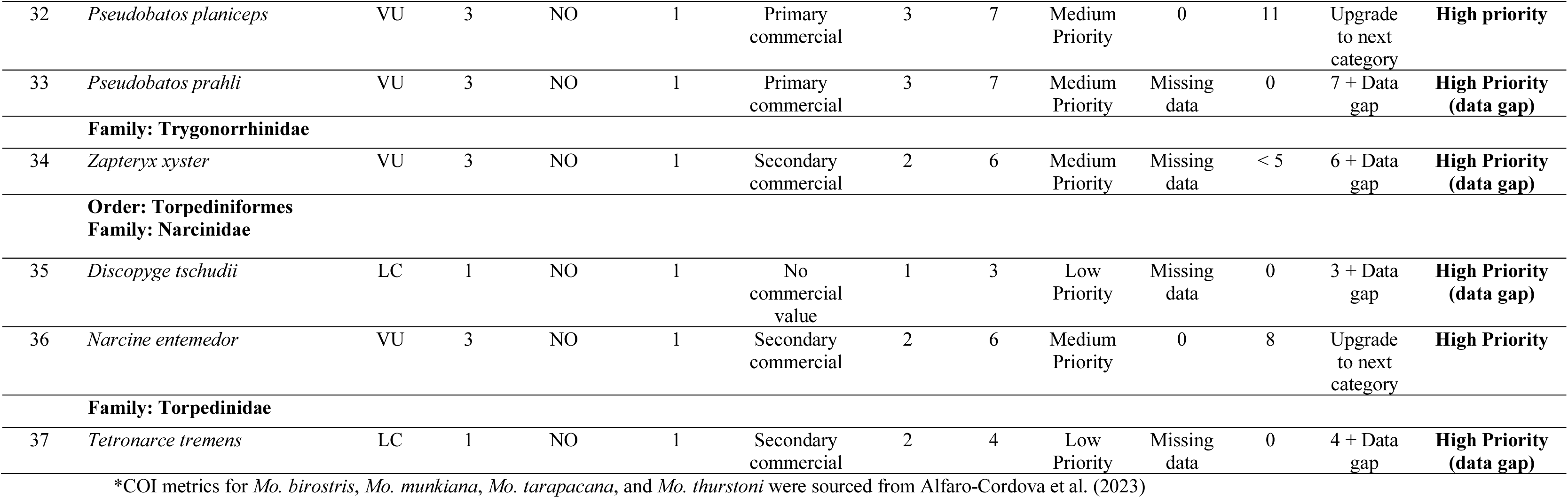
Multi-Criteria Decision Analysis (MCDA) framework for setting research priorities of Peruvian batoids. Tier 1 comprises IUCN conservation status, Eastern South Pacific Ocean (ESPO) endemism, and fisheries pressure. Tier 2 score is based on the nucleotide diversity (*π*) of the cytochrome c oxidase subunit I gene (COI). Sample size (*n*) corresponds to the number of DNA sequences analyzed. For a detailed scoring rubric, parameters, and conditional thresholds, refer to Table S1

Regarding molecular data, only 14 species (37.8%) met our minimum analytical threshold of at least five homologous sequences (*n* ≥ 5) required to calculate the *π* metrics. Within this sequenced cohort, *π* ranged from zero (*H. brevis*, *Mo. birostris, Mo. tarapacana*, *My. longirostris*, *My. peruviana*, *N. entemedor*, and *P. planiceps*) to a maximum value of 0.00213 (*U. chilensis*) (Fig. S6). Under the GDRI framework, 13 of the evaluated species fell below the established cutoff for low diversity (COI-*π* ≤ 0.0021), while *U. chilensis* was assigned the medium genetic diversity level category (0.0021 < COI-*π* ≤ 0.0025). Consequently, this critically constrained genetic diversity triggered a Tier 2 priority upgrade for eight species, elevating them from Medium to High Priority. Conversely, because its molecular metrics sat within the medium-risk tier, *U. chilensis* retained its original Medium Priority status designated by the Tier 1 baseline score. The remaining five species (*Mo. mobular*, *Mo. tarapacana*, *Mo. thurstoni*, *My. chilensis*, and *My. peruviana*) had already been assigned a High Priority status based on their cumulative Tier 1 baseline scores (IUCN status + regional endemism + fisheries pressure).

For the remaining 23 unsequenced taxa (62.2%), the absolute absence (*n* = 0) available records or lower than five sequenced specimens (*n* < 5) triggered the precautionary designation “High Priority (data gap)”, formally categorizing these species as immediate, urgent targets for exploratory genetic research. Notably, this MCDA framework generates the first baseline genetic diversity indexes (*H*, *H_d_*, and *π*) for 8 regional batoids: *G. afuerae* (*n* = 9), *My. chilensis* (*n* = 18), *My. longirostris* (*n* = 7), *My. peruviana* (*n* = 10), *N. entemedor* (*n* = 8), *P. planiceps* (*n* = 11), *S. brevicaudata* (*n* = 5), *U. chilensis* (*n* = 6). A comprehensive summary of all molecular diversity metrics for Peruviand batoid species with accessible COI entries is compiled in Table S6.

## 4. Discussion

During the Golden Age of Natural History, Peru emerged as a key region for marine taxonomy, serving as the type locality for numerous lineages (Tschudi, 1846; Garman, 1880). Notably, nearly one-third (*n* = 11) of the nation’s 37 marine batoid species possess a Peruvian type locality (see the timeline of regional taxonomic discoveries in Fig. S7). This regional baseline began in 1846 when Austrian ichthyologist Johann Jakob Heckel described the first Peruvian marine batoid, the apron ray *Discopyge tschudii* (Tschudi, 1846). Paradoxically, no public COI barcodes from Peruvian specimens exist for this founding species. Subsequent foundational descriptions were added by American naturalist Samuel Garman, including *H. brevis* (Garman, 1880), *P. planiceps* (Garman, 1880), and *My. peruviana* (Garman, 1913). In the 20th century, pioneering work by Peruvian ichthyologist Norma Chirichigno enriched this record with the discoveries of *R. velezi* (Chirichigno, 1973) and *U. tumbesensis* (Chirichigno & McEachran, 1979). Today, with at least 37 marine batoid species (Campos-León et al. 2026), Peru accounts for roughly 5% of global batoid diversity, underscoring its critical role in implementing robust conservation and genetic frameworks to safeguard these historical lineages.

Preserving this rich evolutionary heritage requires modern molecular frameworks to urgently complement traditional conservation and management strategies. In this context, the addition of 72 newly generated sequences across 14 batoid species from regional specimens significantly enriches the open-access molecular database, effectively doubling the prior public library. Nevertheless, a substantial portion of local batoid diversity remains hidden in private databases or completely unrepresented in genetic repositories. By successfully adding barcodes for key species such as *Mo. munkiana*, *Mo. thurstoni*, *My. peruviana*, *R. velezi*, and *Z. xyster*, this research expands the regional public records to encompass 19 species. These expanded molecular data represent a valuable step toward filling the data gaps that hinder effective conservation efforts and data-informed management of threatened Peruvian batoid lineages.

The total absence of reference sequences for five Peruvian batoid species (*A. frerichsi*, *B. aguja*, *B. chapmani*, *B. longicauda*, and *B. peruana*) highlights a persistent challenge in applying DNA barcoding to regional diversity research. These underrepresented taxa consist entirely of deep-sea skates, which face sampling constraints as they are mostly caught opportunistically during exploratory research cruises or as bycatch in deep-water fisheries, such as the Patagonian toothfish longline or trawling fleets (Ebert et al. 2022; Zavalaga et al. 2024). In this regard, our successful assembly and public repository release of the complete mitogenome for *N. martinezi* (GenBank BK075715) represents a significant contribution. By providing a robust set of mitochondrial markers, this resource establishes a critical genomic baseline to accelerate taxonomic resolution and evolutionary reconstructions within these poorly understood deep-sea lineages. Future efforts must target the sampling of these data-deficient batoids, as resolving these barcoding gaps will not only refine molecular identification frameworks but likely unveil cryptic regional diversity.

Despite a collapsed barcoding gap in the extended dataset caused by the shallow interspecific divergence between *G. afuerae* and *G. crebripunctata* alongside cryptic lineages within *R. velezi*, COI effectively identified Peruvian batoid diversity, resolving all assessed taxa into distinct monophyletic MOTUs (Fig. 6). However, exploratory reconstructions exhibited a single deep-node artifact due to limited taxon sampling, where Gymnuridae was nested within Myliobatidae (UFboot = 72). Notably, incorporating two related auxiliary lineages (*Plesiobatis* and *Urolophus*) disrupted this anomalous nesting, grouping these three lineages entirely outside Myliobatidae and recovering *Gymnura* and *Urolophus* as monophyletic sister groups with high support (UFBoot2 = 88; Fig. 6). This isolated misplacement likely stems from long-branch attraction (LBA), exacerbated by a lack of intermediate taxon coverage (Bergsten 2005). Ancient divergence typically causes mutational saturation and homoplasy at the mitochondrial COI locus (Naylor et al. 2012a), erroneously pulling *Gymnura* toward the highly derived clades. This is consistent with its status as an early-diverging lineage that split approximately at 59 mya during the Paleocene (Gales et al. 2024). These findings underline the importance of expanded taxon sampling in mitigating homoplasic noise to restore deep phylogenetic signals.

Notably, *G. crebripunctata* remains the sole taxon on the regional diversity baseline with an unresolved regional geographical occurrence, as it currently lacks both morphological and molecular verification from Peruvian specimens. To date, definitive genetic evidence for this species has only been verified from Mexican samples, leaving its true southern distribution boundaries to be determined (Ehemann et al. 2024b). Recent molecular frameworks including Peruvian butterfly ray specimens consistently assign local samples only to *G. afuerae* (Marín et al. 2018; Ehemann et al. 2024b; Gales et al. 2024; this study). Although the absolute number of genetically analyzed samples remains low, the fact that all Peruvian sequences form a single, independent clade (see Fig. 6) strongly supports *G. afuerae* as the dominant—if not exclusive—lineage in the region. Consequently, the absolute lack of local molecular data and verified morphological records suggests that the presence of *G. crebripunctata* in Peru should be treated with caution, warranting urgent verification to avoid overestimating regional biodiversity.

On the other hand, future taxonomic revisions should target newly discovered Peruvian batoid lineages revealed by recent molecular frameworks. These potentially new regional species include *Narcine* cf. *entemedor* (Rodrigues-Filho et al. 2026), *Rhinoptera* cf. *steindachneri* (Cunha et al. 2026), and *Rostroraja* cf. *velezi* (this study). If further research corroborates the molecular discoveries supporting the validation of these three candidate species, the regional baseline could increase up to 40 species. These findings underscore the utility of molecular tools in revealing hidden batoid diversity, which will ultimately inform and refine local conservation and management strategies.

Our molecular analyses revealed two distinct genetic lineages for *R. velezi*: haplogroup 1 (BIN BOLD:AAX9825; Costa Rica and Tumbes) and haplogroup 2 (BIN BOLD:ADR8057; Piura and Tumbes) (Fig. 7). Pinpointing which lineage represents the original nominal species poses a significant taxonomic challenge, as genetic data are currently unavailable for the historical type material. Notably, a private barcode from Piura (BOLD ID PMFSH1053-21) is assigned to BIN BOLD:ADR8057 (haplogroup 2) (Fig. 7C), matching the exact type locality of the holotype IMARPE: 1213 utilized by Chirichigno (1973). Consequently, this lineage likely represents the true nominal *Rostroraja velezi sensu stricto*, leaving haplogroup 1 as an undescribed cryptic taxon. Alternatively, if direct analysis of the holotype reveals a match with haplogroup 1, then that lineage would retain the historical name, rendering haplogroup 2 as the cryptic lineage. This uncertainty is critical in Tumbes, where both lineages occur in strict sympatry. Ultimately, an exhaustive morphological re-examination coupled with ancient DNA (aDNA) sequencing of the historical IMARPE holotype material remains indispensable to unambiguously assign the historical name.

The critical role of Peruvian marine ecosystems in sustaining manta and devil ray species is well-documented. Peru hosts five *Mobula* species (Alfaro-Cordova et al. 2023; Campos-León et al. 2026), accounting for 50% of the genus’s known global diversity. In addition, Peru and Ecuador sustain the world’s largest population of *Mo. birostris*, which is supported by an essential, shared migratory corridor (Harty et al. 2022). Furthermore, mobulids represent one of the most commercially important Peruvian batoid groups (Alfaro-Cordova et al. 2017, 2023; González-Pestana et al. 2022; Rojas-Perea et al. 2025). Consequently, Peru has been classified as both a priority conservation nation (Palacios et al. 2025) and a highest-risk hotspot (Laglbauer et al. 2026). The recent IUCN uplisting of *Mo. mobular*, *Mo. tarapacana*, and *Mo. thurstoni* to the CR category, which was driven primarily by severe declines in global landings (Jabado et al. 2025) including Peru (Rojas-Perea et al. 2025), demands that molecular conservation initiatives be urgently accelerated and standardized.

This study’s addition of the first Peruvian public COI barcodes for *Mo. munkiana* and *Mo. thurstoni*, alongside additional *Mo. mobular* entries, represents a significant step forward in documenting regional biodiversity. Building on these new data, this research offers the first population genetic assessments for *Mo. mobular* and *Mo. munkiana* in the Eastern Pacific. Our preliminary results revealed a pattern of genetic connectivity between populations from the Northern (Mexico) and Southern (Peru) hemispheres in both species. However, results from *Mo. munkiana* should be taken with caution due to the small sample size of available barcodes. A matching pattern of high genetic connectivity across ocean basins has been observed in related species, including *Mo. birostris* (Humble et al. 2025), suggesting that this widespread gene flow is likely driven by the high migratory capacity of manta and devil rays (Stewart et al. 2018). This high degree of gene flow emphasizes the pivotal role of marine corridors for these transboundary stocks, demanding coordinated international conservation efforts and governance (Harty et al. 2022; Humble et al. 2025).

Interestingly, the presence of private peripheral haplotypes in *Mo. mobular* samples from Mexico and Ecuador (Fig. 8A), diverging by a single mutational step from the ancestral haplotype, points to a potential contemporary female philopatry. In a precautionary management framework for highly threatened elasmobranchs, even shallow maternal segregation warrants attention. Under the contemporary female philopatry scenario, localized depletion in heavily impacted zones like Peru (Laglbauer et al. 2026) is unlikely to be rapidly rescued by immigration from northern feeding grounds. Consequently, further investigations using additional nuclear markers, genomic frameworks, and satellite tracking are urgently warranted to test this hypothesis in order to design effective, localized conservation strategies.

The historical landing data analysis conducted herein indicates that four taxonomic groups still dominate Peruvian landings, which aligns with the findings reported by González-Pestana et al. (2022). These heavily harvested batoid species include *Mo. mobular*, *Myliobatis* spp., *P. planiceps*, and *H. brevis* (Fig. 10 and Fig. 11). Overall, landing records have consistently increased on a yearly basis (Fig. 10), but this does not imply that these are solely attributable to increases in fishing effort in the same fishing areas. There may be other reasons for these increments, including improvements in IMARPE’s capacity to collect data on artisanal fisheries and the expansion of artisanal fleets to new fishing grounds (e.g., spatial-temporal data from the mobulids indicate a relatively recent expansion of the fleet dynamics into oceanic waters; see Fig. S5). It is likely that a mixed effect of different causes could occur at the same time, and that should be assessed for each species with more granular data than the available data for the exploratory analyses presented in this research.

Our findings show a positive linear trend of overall batoid landings across the 29 years of data. However, if the time series is limited to 2015 onwards, i.e., when the amount of annual data made a leap that seems to be structural and has been maintained over time, we do not find such a steep linear trend. Instead, we found a non-significant linear trend in the aggregate batoid landings data (P-value ∼0.461), as well as in the *Myliobatis* (P-value ∼0.609) and *Mobula* groups (P-value ∼0.692). The lack of a significant trend in recent years could signal that landings have stabilized after a period of sharp increases in fishing effort and/or data-collection capacity. Access to unaggregated fishing data at the finest scale (e.g., by fishing trip and/or cast) and by single species, in conjunction with biological and molecular information, would allow researchers to make further inferences regarding the population status of these species and assess whether the current information effectively communicates risks that threaten the long-term viability of batoid species.

The research priority framework implemented herein reveals an alarming trend where intense fishing pressure directly overlaps with high risk extinction and severely depleted genetic diversity. Although the Tier 1 assessment successfully segregated species into three distinct categories (Low Priority = 37.8%, Medium Priority = 48.6%, and High Priority = 13.6%), the integration of Tier 2 molecular metrics elevated an additional 31 species from low or medium status, ultimately consolidating 36 of the evaluated species into the High Priority domain. This systematic consolidation underscores that molecular data voids represent critical blind spots in traditional, landings-based fishery management frameworks.

The sole exception to this predominant High Priority group was *U. chilensis*, whose *π* estimate (0.00213) marginally exceeded the GDRI low-genetic diversity threshold (*π* ≤ 0.0021; Fig. S6), thereby retaining its Medium Priority classification. This moderate diversity level in *U. chilensis* is likely sustained by the shorter generation times and multi-annual reproductive cycles typical of small round stingrays (Mejía-Falla et al. 2012; Alvarez-Fuentes et al. 2023). Such life history traits provide a unique axis of resilience for this taxonomic group within the generally vulnerable batoid lineage, approaching diversity patterns more typical of teleost fishes with high fecundity and large population sizes (Ravi and Venkatesh 2018; Ferragut-Perello et al. 2026).

Most analyzed batoid taxa with available molecular data (*n* = 14) fell within the low genetic diversity tier (COI-*π* ≤ 0.0021). This critical scenario in Peruvian rays was recently highlighted by Marín et al. (2026), who reported a single COI haplotype (*H_d_* = 0, *π* = 0) for *H. brevis* (*n* = 27) across Peru. Our nine new specimens from Pisco confirm the persistence of this unique haplotype, reinforcing the severity of this genetic depletion trend for the Peruvian diamond stingray. Similarly, our findings revealed that the severely exploited *My. peruviana* (*n* = 10) and *P. planiceps* (*n* = 11) exhibited zero COI variation between geographically distant locations (Piura and Ica regions). This alarming monomorphic pattern was also observed in other commercially exploited batoids (*Mo. birostris*, *Mo. tarapacana*, *My. longirostris*, *N. entemedor*) albeit across more localized areas (Tumbes and/or Piura regions) and with limited sample sizes (5 ≤ *n* ≤ 14) (Fig. S6).

This pervasive pattern of either null COI variability (observed across seven species) or low diversity (COI-*π* ≤ 0.0021; detected in six species) likely signals historical population bottlenecks or ongoing overexploitation within these regional batoid lineages. Intense, unmanaged fishing pressure removes a significant portion of the breeding biomass, eroding mitochondrial diversity and reducing the representation of unique maternal lineages (Petit-Marty et al. 2022; Sadler et al. 2023). A clear example of this phenomenon is the critical mitogenomic erosion observed in *P. pristis*, where anthropogenic impacts have extirpated local populations across its global geographic range, erasing 29 of the 35 historical maternal lineages (Fearing et al. 2026). Consequently, these long-lived, slow-reproducing elasmobranchs are left highly vulnerable and genetically weakened when coping with shifting environmental regimes or rapid climate changes (Dulvy et al. 2021).

These findings are of particular concern given that Peruvian batoids are episodically subjected to severe environmental stress driven by major oceanographic anomalies such as El Niño-Southern Oscillation (ENSO) events. While our preliminary genetic screening underscores an urgent need for expanded geographic and taxonomic sampling to determine if genetic depletion is uniform across regional populations, concrete management actions must be implemented now to mitigate further genetic erosion within Peruvian batoids.

For the remaining 23 taxa, an absolute lack of data or insufficient sample sizes (*n* < 5) prevented the estimation of molecular diversity metrics altogether. Consequently, these species were classified as High Priority under a precautionary “data gap” designation. This widespread data deficiency highlights the urgent need for targeted field sampling campaigns and molecular exploration to properly assess these highly vulnerable, unmonitored populations. Although our threshold (*n* ≥ 5) represents a limited, preliminary sample size due to local fishery and database constraints, it successfully captured the first molecular indicators for data-deficient taxa. The two-tiered priority system established herein provides a clear, standardized path forward for batoid management by successfully integrating ecological, fishery, and molecular data, a multi-criteria framework that remains completely unprecedented in the history of Peruvian fisheries management.

During the current golden age of integrative and molecular taxonomy, the BATOseq-PE library emerges as a valuable open-access genetic dataset, covering 56% (14 species) of the 25 commercially important Peruvian batoid species (Zavalaga et al. 2021), along with the complete mitogenome sequence of *N. martinezi*. As the first self-funded, regional initiative of its kind, this novel dataset establishes a public DNA reference library designed to spearhead batoid conservation efforts and drive informed fisheries management. Furthermore, we developed the first specialized tool to assess the genetic health (via the GDRI approach) and research priorities of Peruvian batoids. It was accomplished by combining our BATOseq-PE dataset with publicly available records, conservation status, endemism metrics, and fishery statistics, which aims to elevate national fisheries management to an unprecedented standard. We expect that our standardized protocols will serve as a foundational blueprint to scale these conservation efforts to other threatened or heavily exploited fish groups.

## 5. Future research and recommendations

The ecological significance and vulnerability of Peruvian batoids demand a swift integration of modern molecular frameworks into fisheries management. While the BATOseq-PE library significantly expanded public Peruvian sequences, additional sampling efforts must target remaining underrepresented lineages to achieve a taxonomically complete DNA reference library. Moving forward, our preliminary COI diversity indicators, which suggest a high incidence of low genetic diversity, should be supplemented with multilocus nuclear data, genome-wide SNP markers, and complementary functional molecular approaches (see Table 3). Integrating advanced molecular tools with traditional fisheries metrics will provide crucial baseline needed to advance from aggregated landing statistics into accurate species-specific population assessments.

**Table 3.**
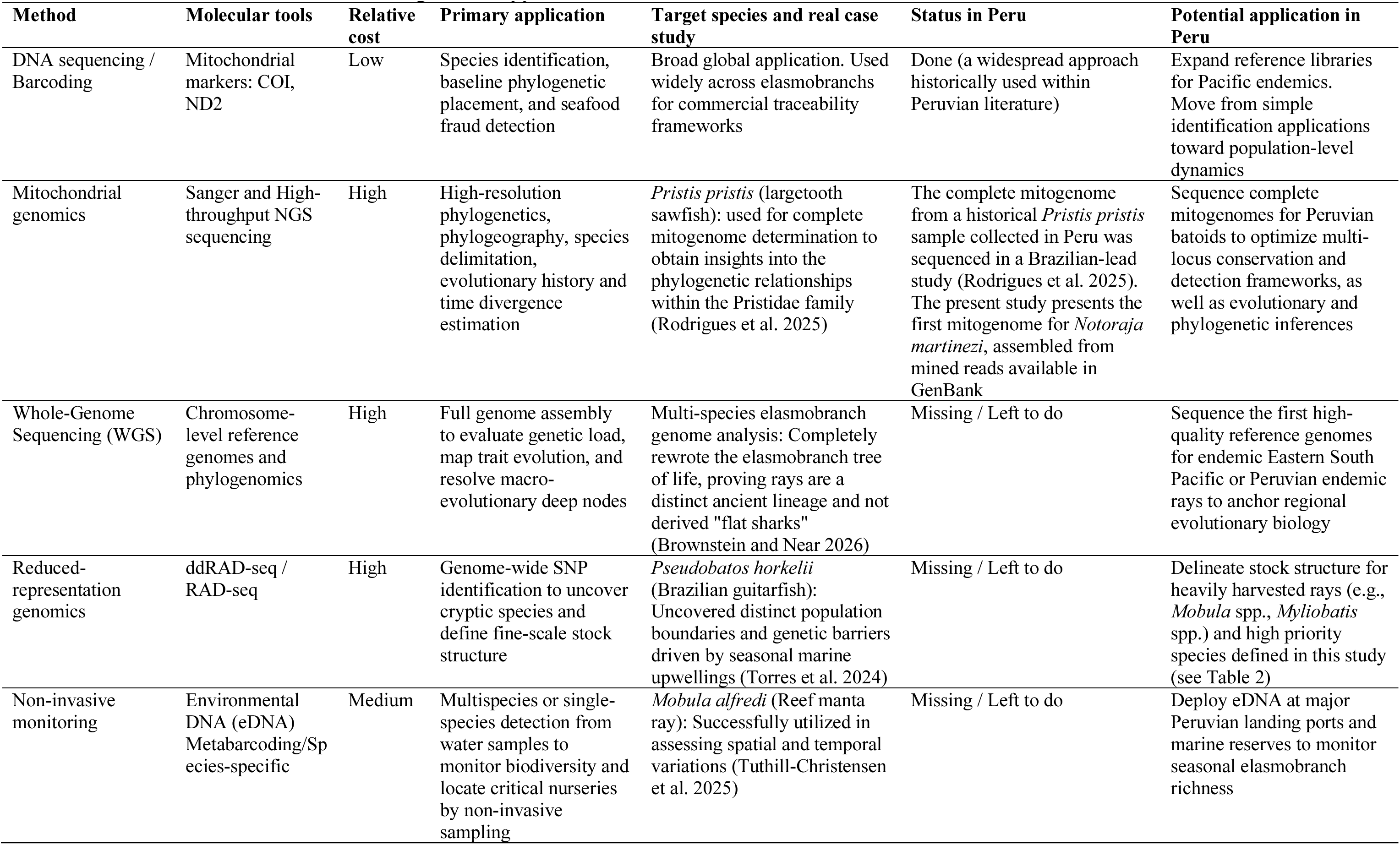
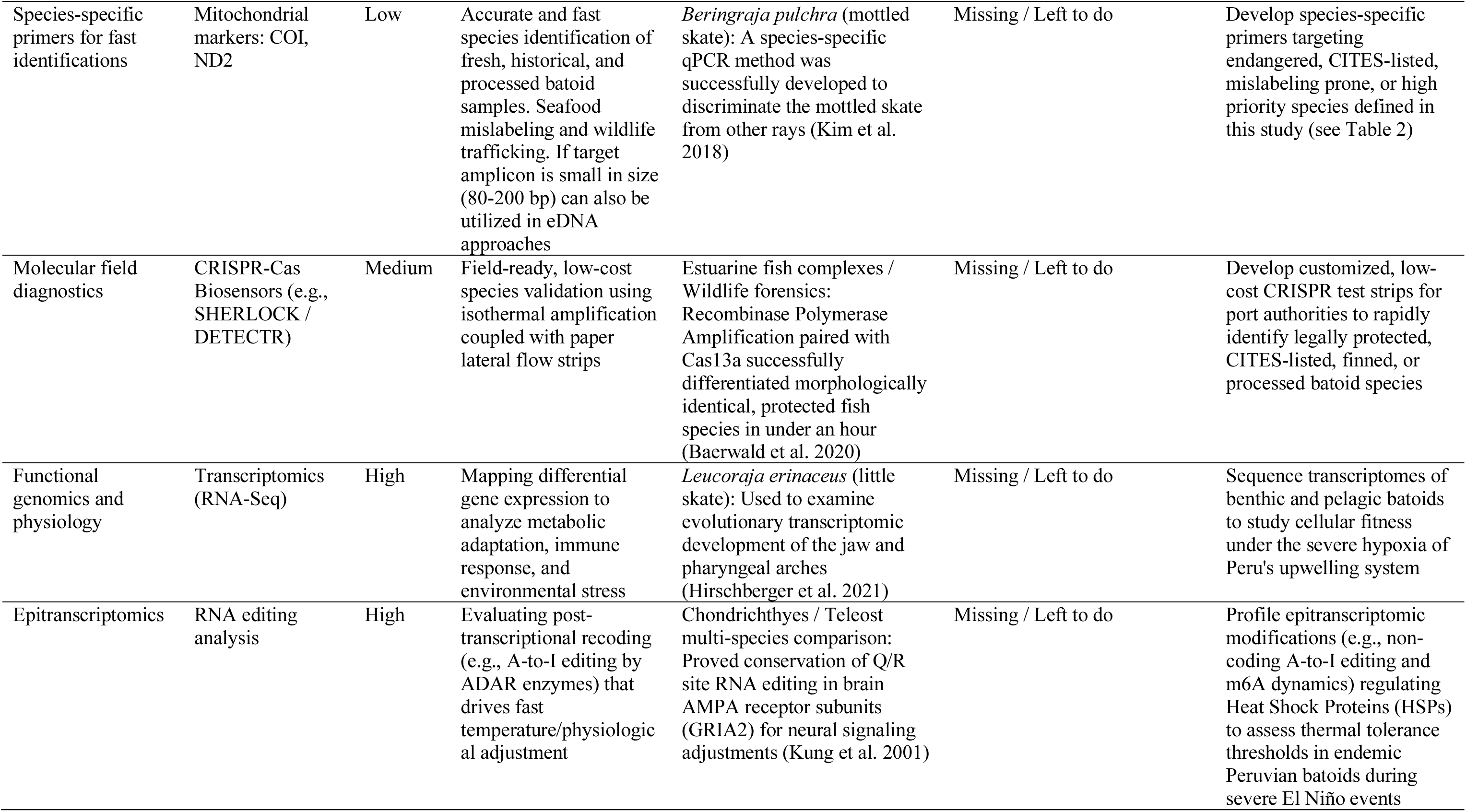
Potential molecular methods and genomic applications for future research on Peruvian batoids

Artisanal fishing is likely the primary driver of batoid population impacts. Moreover, with some exceptions, these species are practically unmanaged within Peruvian waters. For heavily exploited commercial taxa displaying signs of genetic depletion (*H. brevis*, *My. peruviana*, and *P. planiceps*) we recommend the establishment of urgent, immediate species-specific precautionary conservation and management measures, such as catch limits and/or seasonal closures. Conversely, for data-deficient species, we encourage the rapid development of genetic health baselines by implementing regular tissue sampling and molecular monitoring at major landing ports.

To complement this baseline molecular dataset, researchers are encouraged to release private regional barcodes, correct misidentifications, and update provisional assignments to improve database reliability and taxonomic coverage for priority batoids. Ultimately, this refined molecular library must translate into field-level compliance. The chronic lack of taxonomic resolution for major species groups should be addressed by implementing an official list of standardized commercial names, derived from eliminating ambiguous catch-all labels like “*raya*” to secure supply-chain traceability.

Finally, policymakers and governance frameworks must expand to match the geographic scale of highly migratory transboundary stocks, such as *Mobula* species. Peru is well-positioned to promote bilateral monitoring frameworks and coordinated international governance (e.g., with Ecuador and Mexico) to safeguard critical shared marine corridors, while simultaneously closing legal loopholes, such as those permitting the incidental exploitation of non-*birostris* mobulids. Synthesizing IMARPE’s valuable three-decade historical landing datasets with modern, open-access molecular repositories will allow policymakers to move past a state-centric model, establishing a modern, data-driven, and participatory national and localized batoid management plan.

## Author contributions

**AM, LESR, RGW, EZM**: conceptualization.

**AM**: wrote the original draft, data curation, formal analysis, developed figures and tables, software, mitogenome assembly and genetic characterization, validation, supervision.

**AM, LESR, RGW, SRP, CVL, SPM, ELL, LERF, EZM**: investigation, sampling design and logistics.

**LESR, AYU, SRP, CVL, SPM, ELL, LERF, EZM**: field work.

**LESR, AYU, SVL, LERF**: molecular experiments, data analysis.

**AM, LESR, LERF**: phylogenetic analyses.

**RGW**: data curation, fisheries analysis, developed figures, wrote fisheries section.

**EZM**: administered and supervised the project.

All authors reviewed, contributed, and approved the submitted version.

**Funding:** This work received no funding.

**Data availability:** All genetic sequences generated herein are available in the GenBank database under accession numbers PZ899860 to PZ899931, PZ901813, PZ901814, and BK075715.

**Conflict of interest:** The authors declare no competing interests.

## Acknowledgements

We are grateful to N. Ehemann for his valuable comments and insightful discussions regarding the taxonomic status, regional distribution, and life-history of Eastern Pacific *Gymnura* and *Urotrygon* species. The authors extend their gratitude to the *Instituto Público de Investigación de Acuicultura y Pesca* (IPIAP) and AMAREA for providing the illustrations of five *Mobula* species.

## Supplementary files

**Appendix S1** *Notoraja martinezi*: technical metadata, genetic characterization, mitogenome map, BOLD identification tree. Note: this file contains Table S3, Table S4, Fig. S1 and Fig. S2

**Appendix S2** Detailed analysis of the historical trends, fishing gear contributions, and expansion associated with the *Myliobatis* and *Mobula* fisheries. Note: this file contains Fig. S4 and Fig. S5

**Appendix S3** Automated Excel file containing the MCDA-research priority framework

**File S1** PCR amplification protocols

**Fig. S1** Circular map of the complete mitochondrial genome of *Notoraja martinezi*

**Fig. S2** Phylogenetic tree identification of our query COI sequence of *Notoraja martinezi* (highlighted in red) using the BOLD Taxon ID engine

**Fig. S3** IUCN conservation status of 37 Peruvian marine batoid species

**Fig. S4** Differences in average annual catch between Period 1 (1996-2014) and Period 2 (2015-2025) for *Myliobatis* spp., categorized by 0.2° by 0.2° grid cells, fishing gear, and the classification of fishing zones (traditional or newly established)

**Fig. S5** Differences in average annual catch between Period 1 (1996-2014) and Period 2 (2015-2025) for *Mobula* spp., categorized by 0.2° by 0.2° grid cells, fishing gear, and the classification of fishing zones (traditional or newly established)

**Fig. S6** Intraspecific nucleotide diversity (*π*) of the cytochrome c oxidase subunit I gene (COI) across 14 Peruvian marine batoid species. The horizontal dotted line represents the low genetic diversity threshold (COI-*π* ≤ 0.0021) based on the Genetic Diversity Risk Indicator (GDRI) approach. Sample sizes (*n*) are indicated in parentheses along the x-axis

**Fig. S7** Timeline of Peruvian taxonomic and molecular research for marine batoids. Type locality: TL

**File S1** PCR amplification protocols

**a)** PRIMER SET by Folmer et al. (1994)

Primer F: LCO1490 (5’-GGTCAACAAATCATAAAGATATTGG-3’) Primer R: HCO2198 (5’-TAAACTTCAGGGTGACCAAAAAATCA-3’)

Fragment size after primer removal: 658 bp

**Target taxa**: Hypanus brevis, Pseudobatos planiceps, Zapteryx xyster

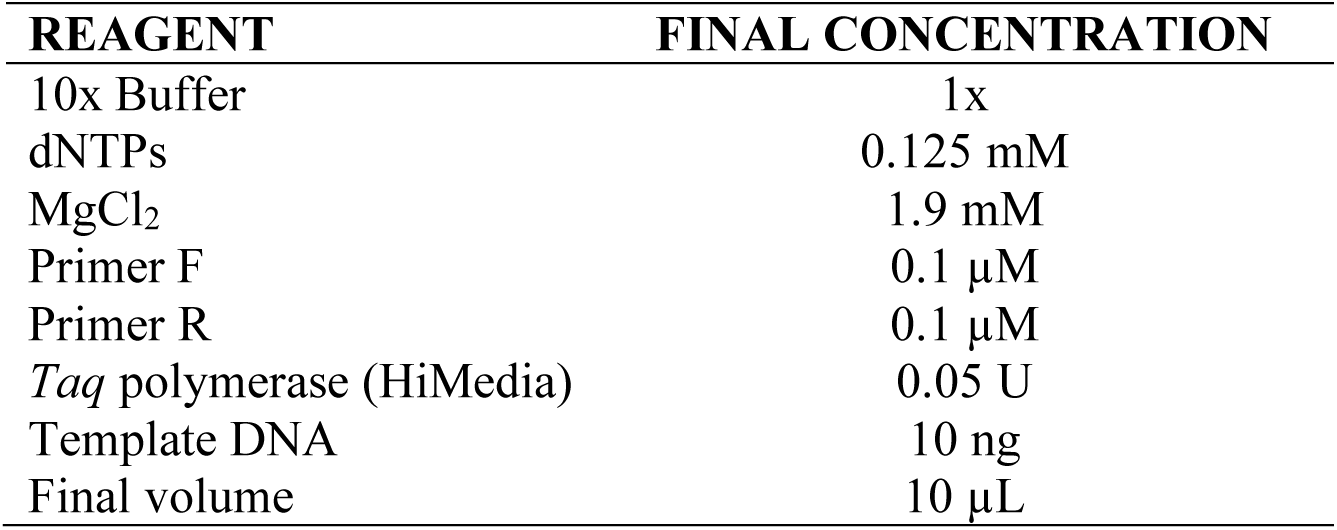

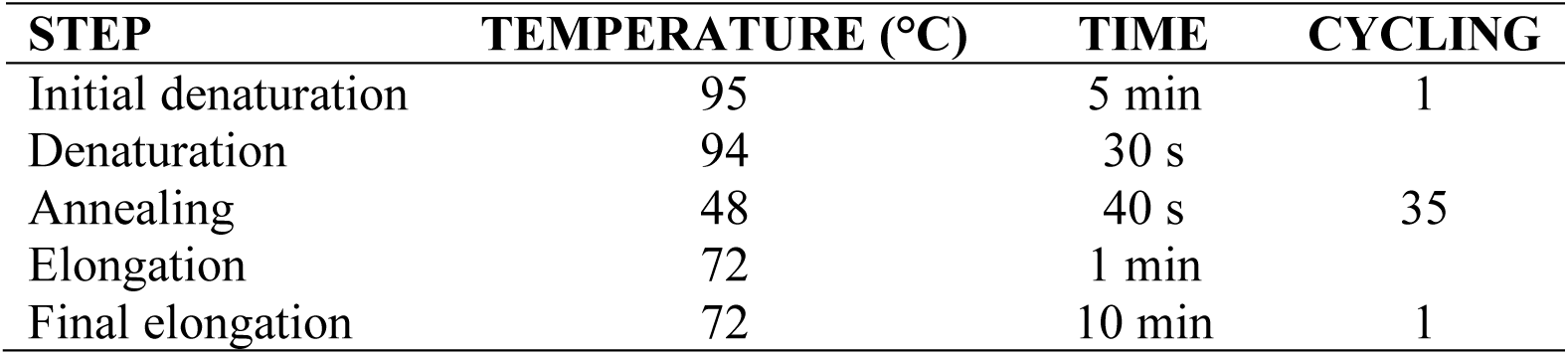

| STEP | TEMPERATURE (°C) | TIME | CYCLING |
| --- | --- | --- | --- |
| Initial denaturation | 95 | 5 min | 1 |
| Denaturation | 94 | 30 s |  |
| Annealing | 48 | 40 s | 35 |
| Elongation | 72 | 1 min |  |
| Final elongation | 72 | 10 min | 1 |

**b)** PRIMER SET by Ward et al. (2005):

Primer F: FishF1 (5’-TCAACCAACCACAAAGACATTGGCAC-3’) Primer R: FishR1 (5’-TAGACTTCTGGGTGGCCAAAGAATCA-3’)

Fragment size after primer removal: 655 bp

**Target taxa**: Aetobatus laticeps, Gymnura afuerae, Hypanus brevis, Mobula spp., Myliobatis spp., Urotrygon chilensis

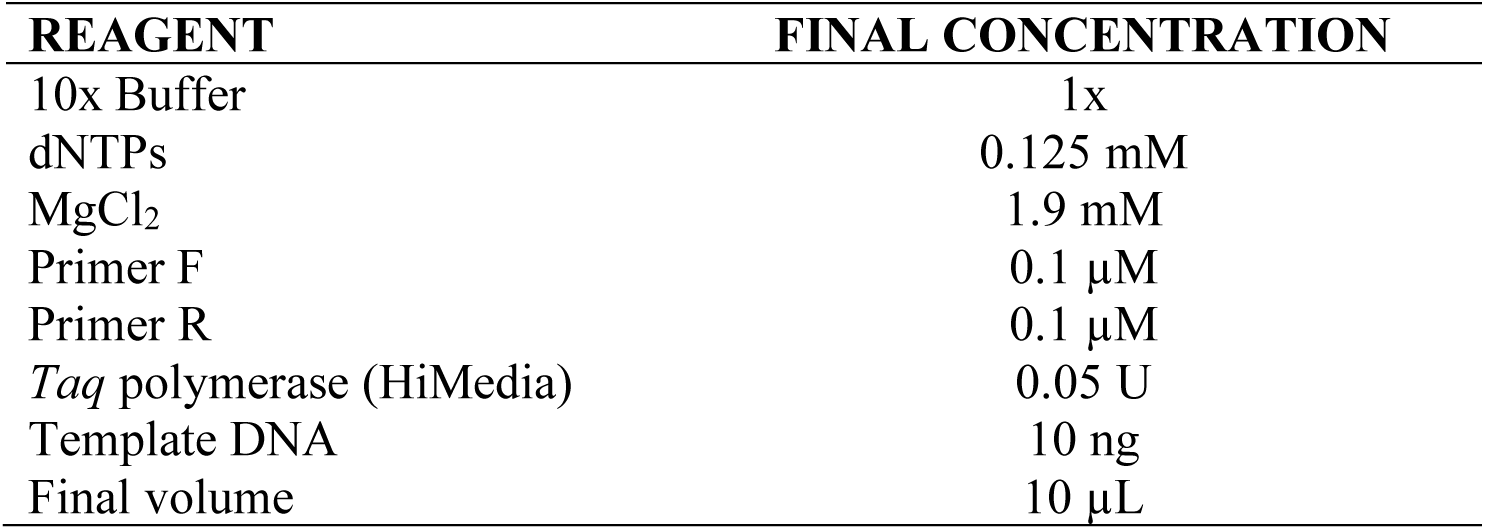

| STEP | TEMPERATURE (°C) | TIME | CYCLING |
| --- | --- | --- | --- |
| Initial denaturation | 95 | 5 min | 1 |
| Denaturation | 94 | 30 s |  |
| Annealing | 54 | 40 s | 32 |
| Elongation | 72 | 1 min |  |
| Final elongation | 72 | 10 min | 1 |

**c)** PRIMER SET COI-1 by Ivanova et al. (2007):

Primer F: FF2d (5’-TTCTCCACCAACCACAARGAYATYGG-3’) Primer R: FR1d (5’-CACCTCAGGGTGTCCGAARAAYCARAA-3’)

Fragment size after primer removal: 655 bp

**Target taxon**: Rostroraja velezi

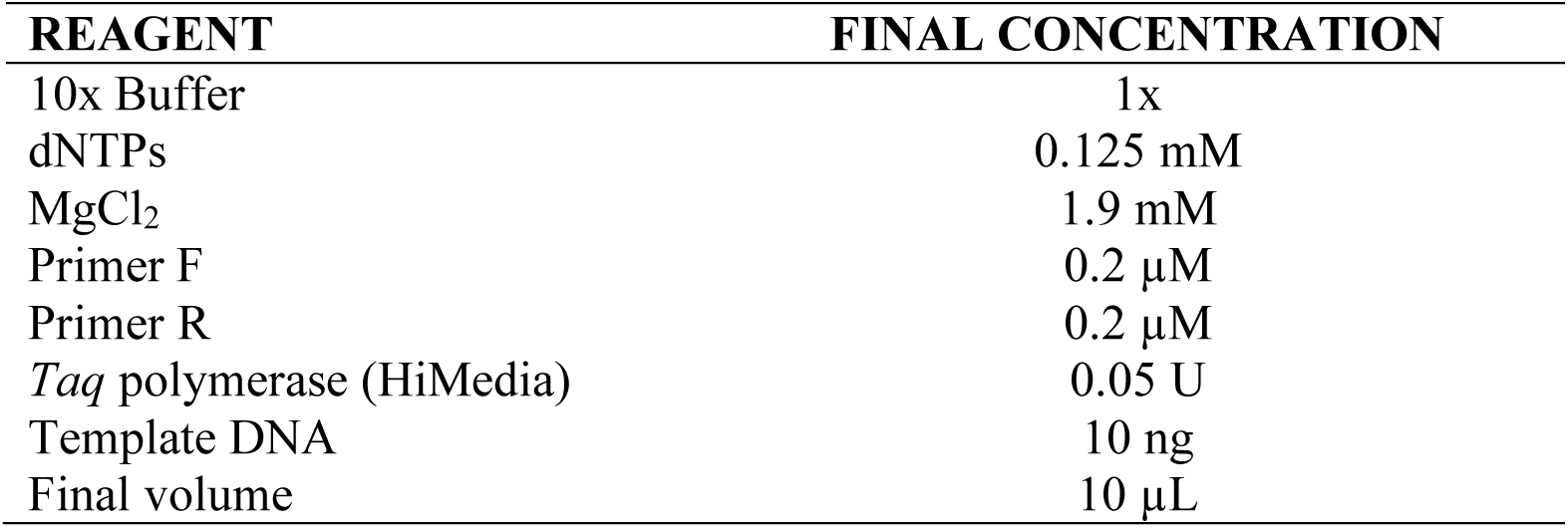

| REAGENT | FINAL CONCENTRATION |
| --- | --- |
| 10x Buffer | 1x |
| dNTPs | 0.125 mM |
| MgCl<sub>2</sub> | 1.9 mM |
| Primer F | 0.2 µM |
| Primer R | 0.2 µM |
| *Taq* polymerase (HiMedia) | 0.05 U |
| Template DNA | 10 ng |
| Final volume | 10 µL |

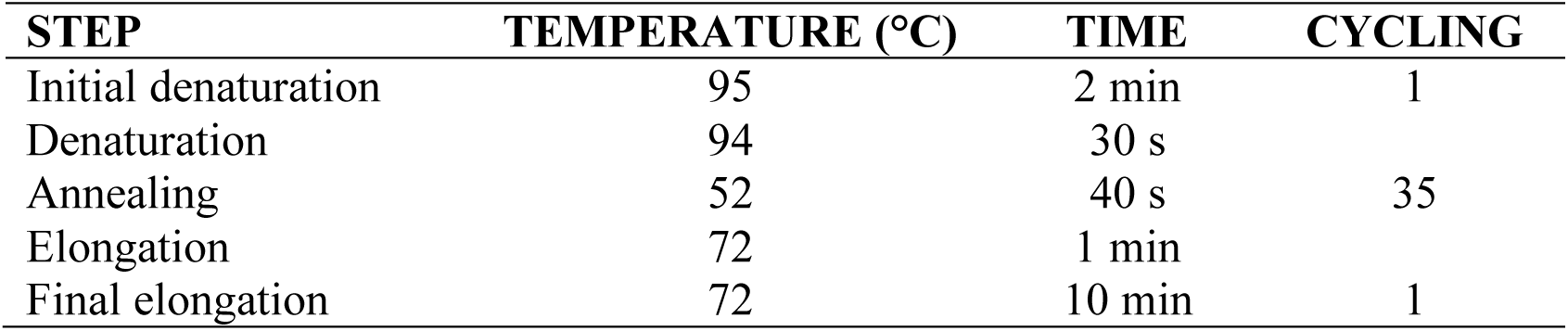

### **d)** PRIMER SET COI-3 (”cocktail”) by Ivanova et al. (2007)

Primer F1: VF2_tl (5’-tgtaaaacgacggccagtCAACCAACCACAAAGACATTGGCAC −3’) Primer F2: FishF2_tl (5’-tgtaaaacgacggccagtCGACTAATCATAAAGATATCGGCAC-3’) Primer R1: FR1d_tl (5’-caggaaacagctatgacACCTCAGGGTGTCCGAARAAYCARAA-3’) Primer R2: FishR2_tl (5’-caggaaacagctatgacACTTCAGGGTGACCGAAGAATCAGAA-3’)

Fragment size after primer removal: 652 bp

**Target taxa**: Hypanus brevis, Mobula spp., Myliobatis spp., Narcine entemedor, Rostroraja velezi

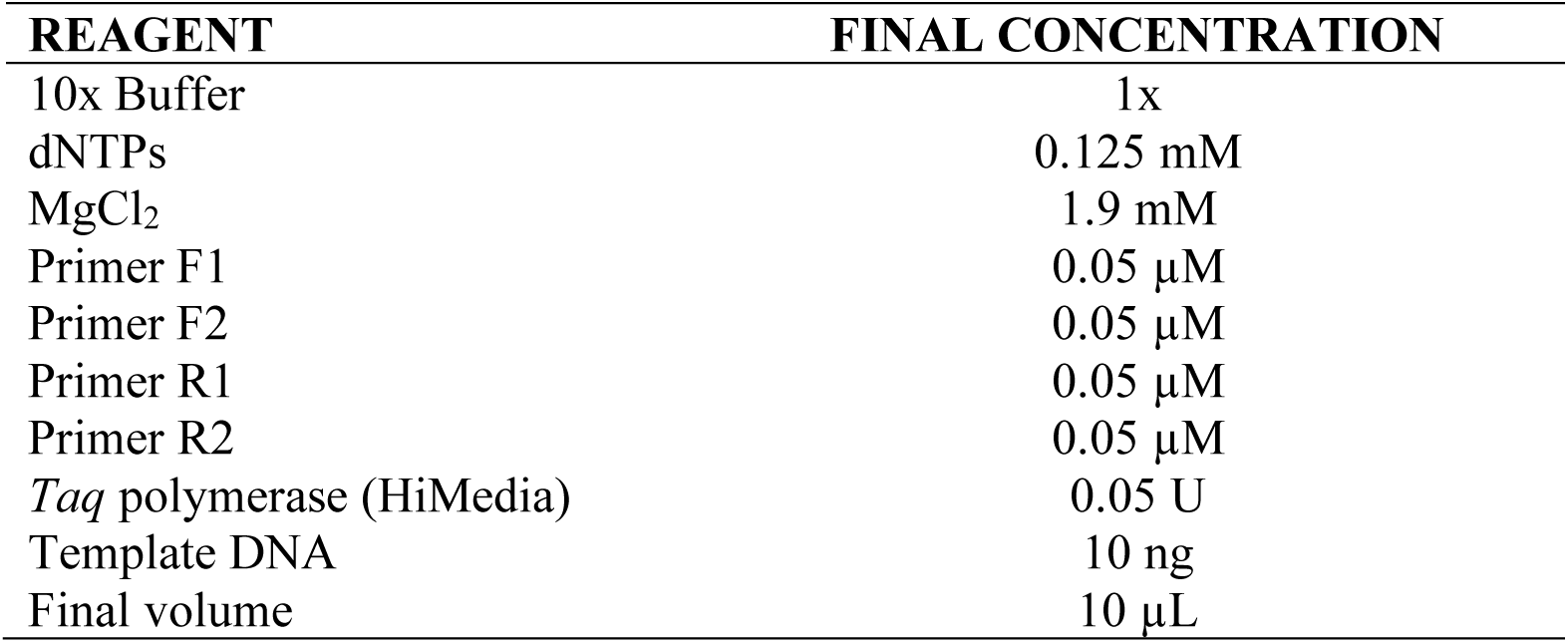

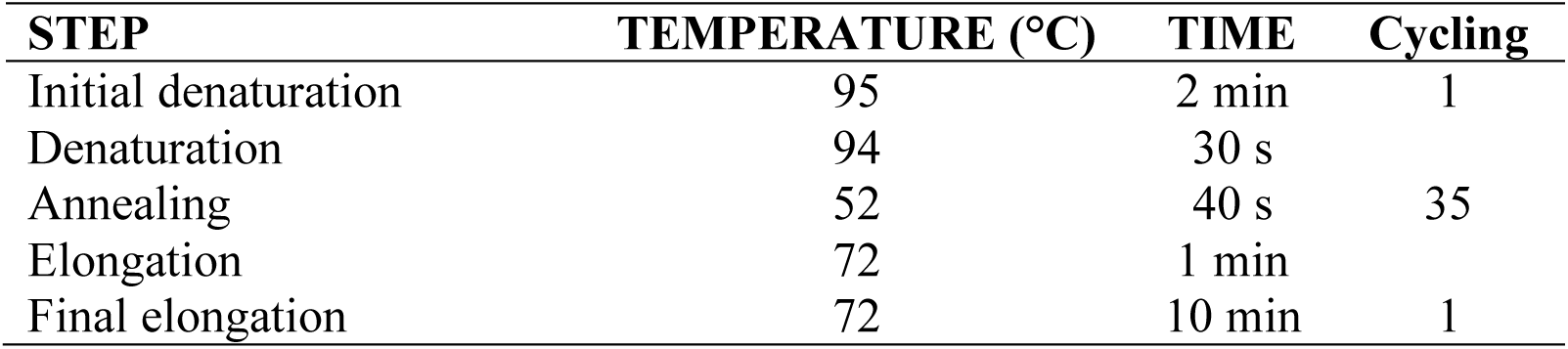

**Appendix S1** Notoraja martinezi: technical metadata, genetic characterization, mitogenome map, BOLD identification tree

Note: this file contains Table S3, Table S4, Fig. S1, and Fig. S2

**Appendix S2:** Detailed analysis of the historical trends, fishing gear contributions, and expansion associated with the *Myliobatis* and *Mobula* fisheries.

Note: this file contains Fig. S4 and Fig. S5

### Landing trends and spatial analysis for the eagle rays (*Myliobatis* spp.) fishery

Landings from *Myliobatis* spp. group averaged 365 tons·year^-1^ (SD = 58) in Period 1 and doubled this magnitude from 2015 onwards (mean = 778, SD = 146), and the whole-time series fit relatively well with a linear regression (Fig. 11B; ∼24 tons·year^-1^; adjusted R^2^ = 0.712). In particular, average annual landings with gillnets exceeded those in Period 2 by double that of Period 1 (P-value ∼0.000), rising from 325 tons·year^-1^ (SD = 162) to 733 tons·year^-1^ (SD = 139). However, fishing with gillnets has maintained a similar relative magnitude across both periods, rising from 89% of total landings in Period 1 to 94% in the next. In total, 466 distinct 0.2° by 0.2° fishing zones were covered by gillnetters during Period 1, predominantly concentrated in the center and northern areas of the country, where some significant increments in the average catches can also be seen between periods (see the top-right Panel of Fig. S4). In Period 2, the reported areas expanded to 190 additional zones (top-left Panel of Fig. S4). Reports of purse-seine fishing have remained at the same magnitude (p-value ∼0.866), at 23 tons·year^-1^ (SD = 12) in Period 1 and 27 tons·year^-1^ (SD = 25) in Period 2 (bottom Panels of Fig. S4).

### Landing trends and spatial analysis for the devil rays (*Mobula* spp.) fishery

Landings from the species group of the *Mobula* genus averaged 160 tons·year^-1^ (SD = 152) in Period 1 and exceeded double this in the next period (mean = 378, SD = 72). The time series fit acceptably with a linear regression (Fig. S5; ∼13 tons·year^-1^; adjusted R^2^ = 0.480). In particular, average annual landings with gillnets increased almost fivefold from Period 1 to Period 2 (i.e., 2015-2025), rising from 66 tons·year^-1^ (SD = 53) to 326 tons·year^-1^ (SD = 68). In total, 76 distinct fishing zones were utilized by gillnetters during Period 1, predominantly concentrated in the northernmost areas of the country (see the top-right Panel of Fig. S5). Then, in Period 2, reported areas expanded significantly, covering eight times more area, notably extending into more oceanic waters and central regions (top-left Panel of Fig. S5). Although this fishing gear attained considerable significance in Period 2, accounting for 86% of the total genus landings during that interval of time, the relative importance of purse seine gear declined, decreasing from 56% to only 7% of the total annual landings (shifting from a mean ± SD of 56 ± 124 to 28 ± 30; P-value ∼ 0.134). Consequently, the reported fishing zones of artisanal purse seiners did not experience such expansion as the one presented for gillnetters (bottom-left Panel of Fig. S5), and even in some very coastal areas showed a diminishment of the average landing per year (bottom-right Panel of Fig. S5)

---

**Fig. S1.**
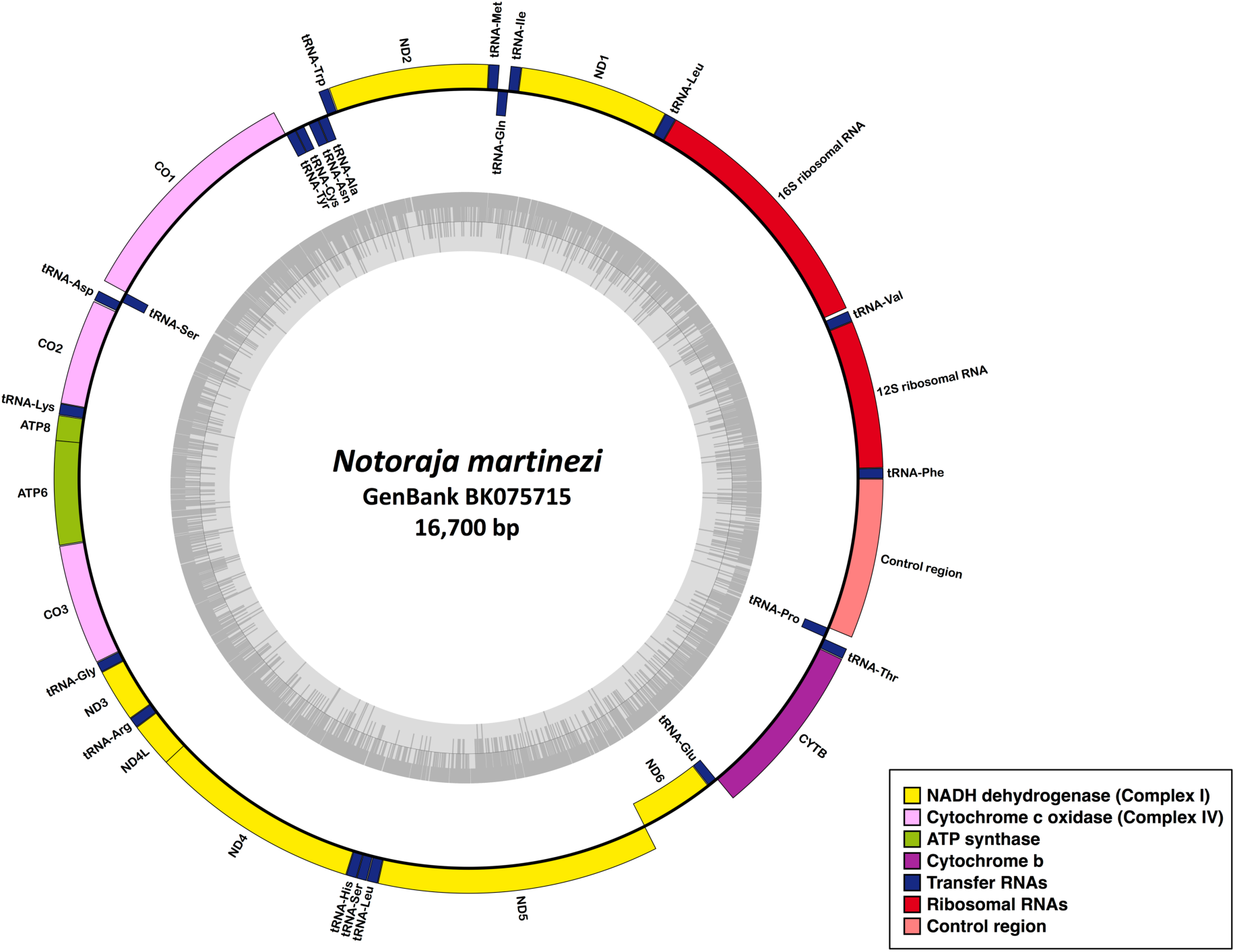
Circular map of the complete mitochondrial genome of *Notoraja martinezi*

**Fig. S2.**
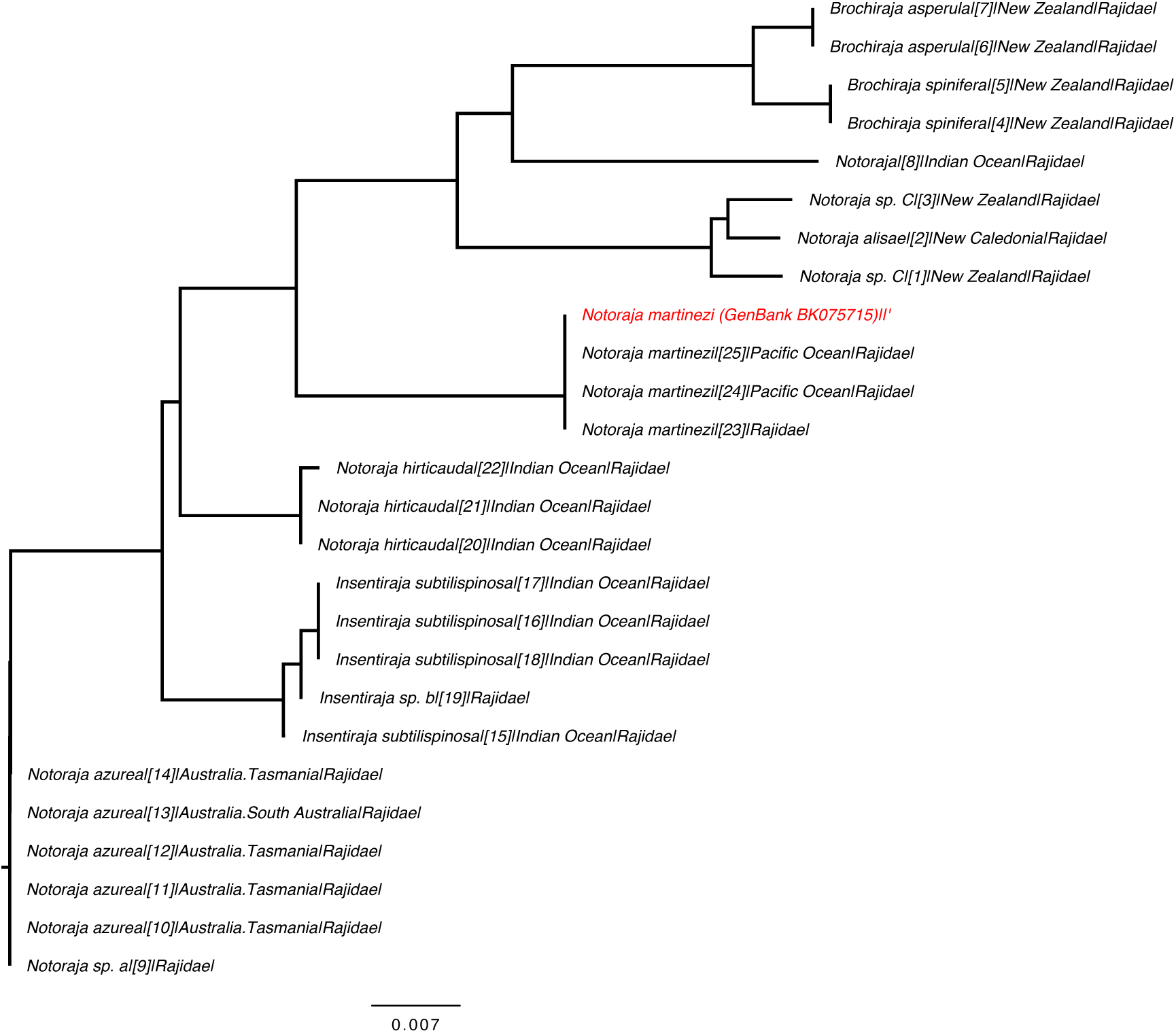
Phylogenetic tree identification of our query COI sequence of *Notoraja martinezi* (highlighted in red) using the BOLD Taxon ID engine

**Fig. S3.**
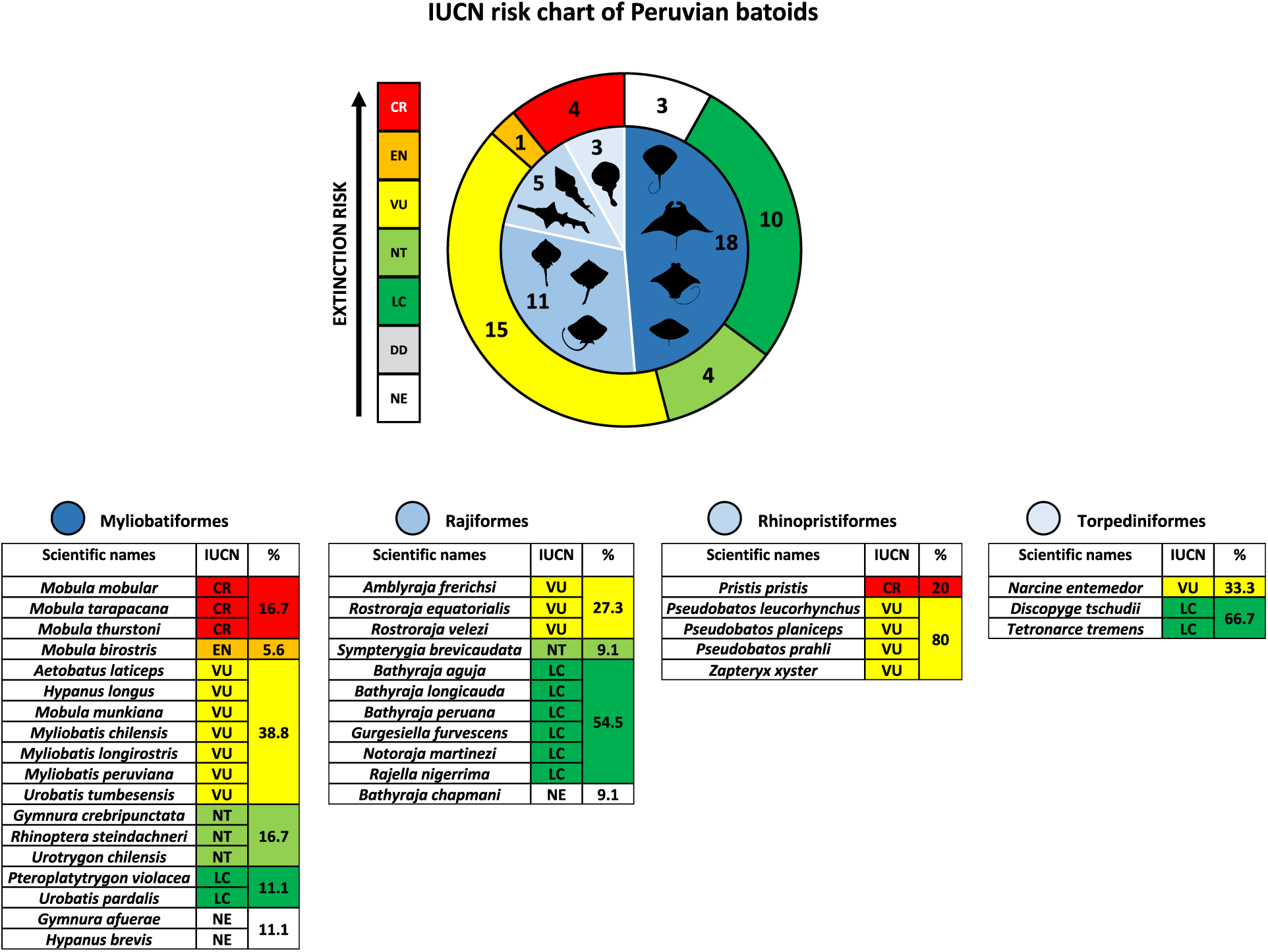
IUCN conservation status of 37 Peruvian marine batoid species

**Fig. S4.**
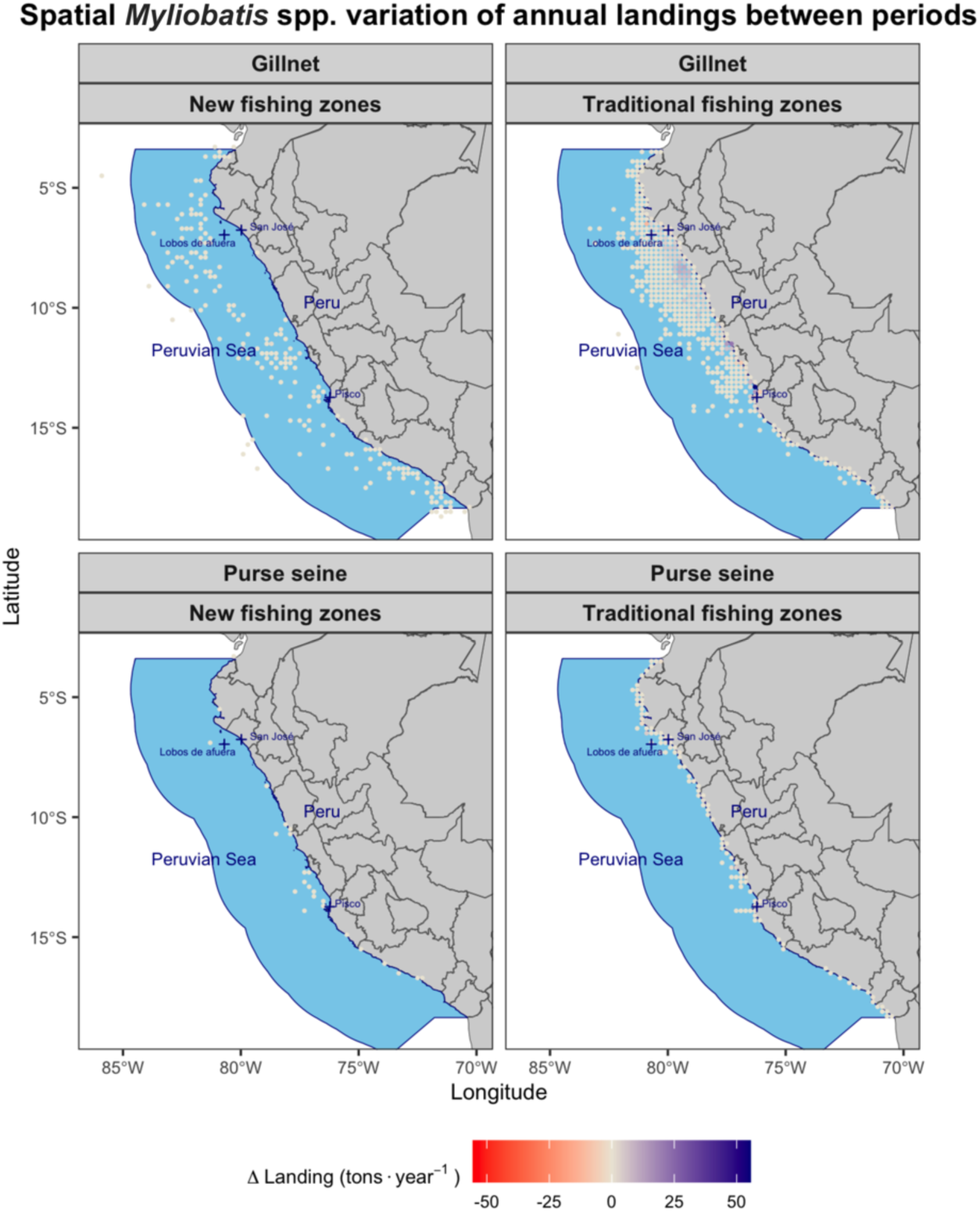
Differences in average annual catch between Period 1 (1996-2014) and Period 2 (2015-2025) for *Myliobatis* spp., categorized by 0.2° by 0.2° grid cells, fishing gear, and the classification of fishing zones (traditional or newly established)

**Fig. S5.**
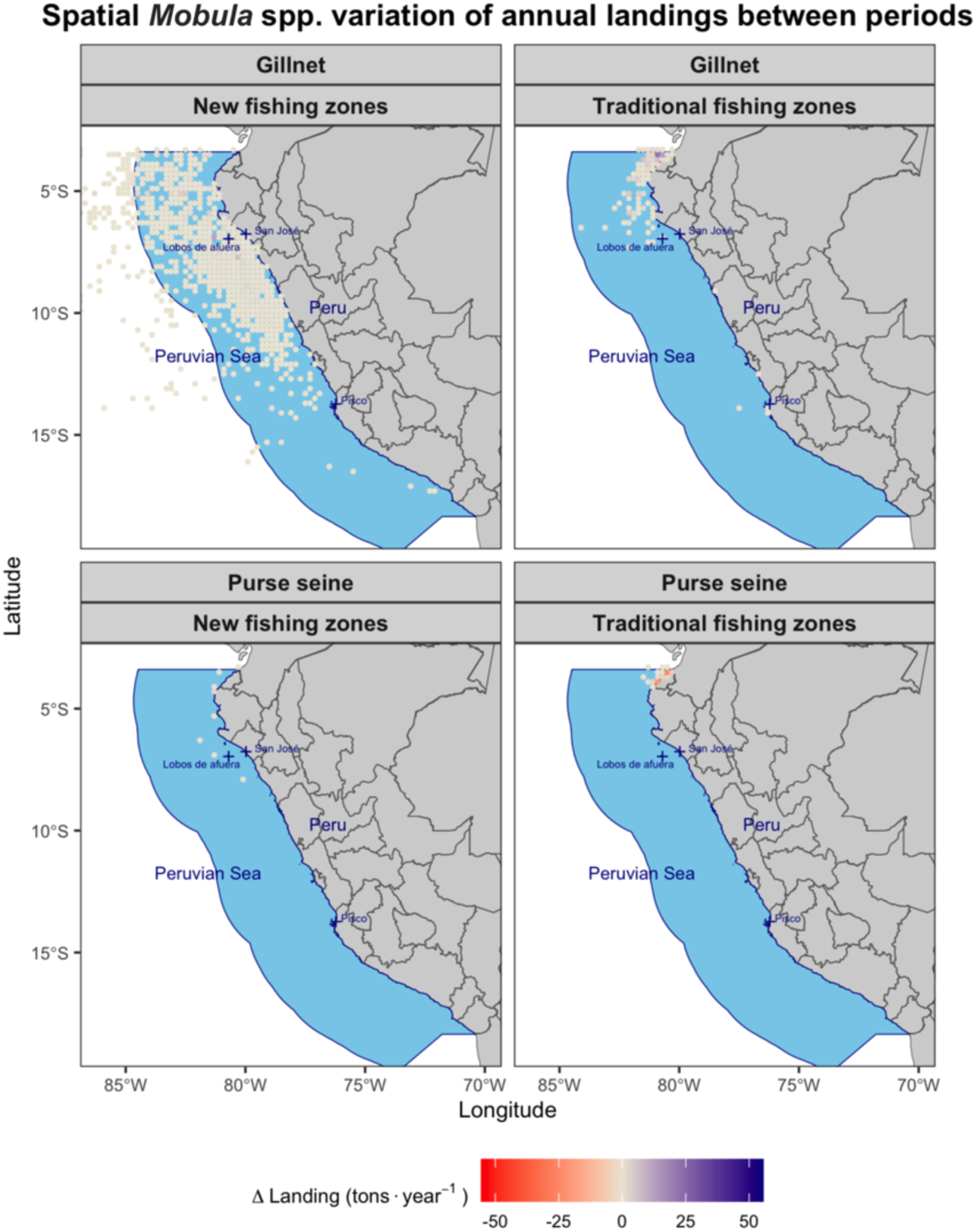
Differences in average annual catch between Period 1 (1996-2014) and Period 2 (2015-2025) for *Mobula* spp., categorized by 0.2° by 0.2° grid cells, fishing gear, and the classification of fishing zones (traditional or newly established)

**Fig. S6.**
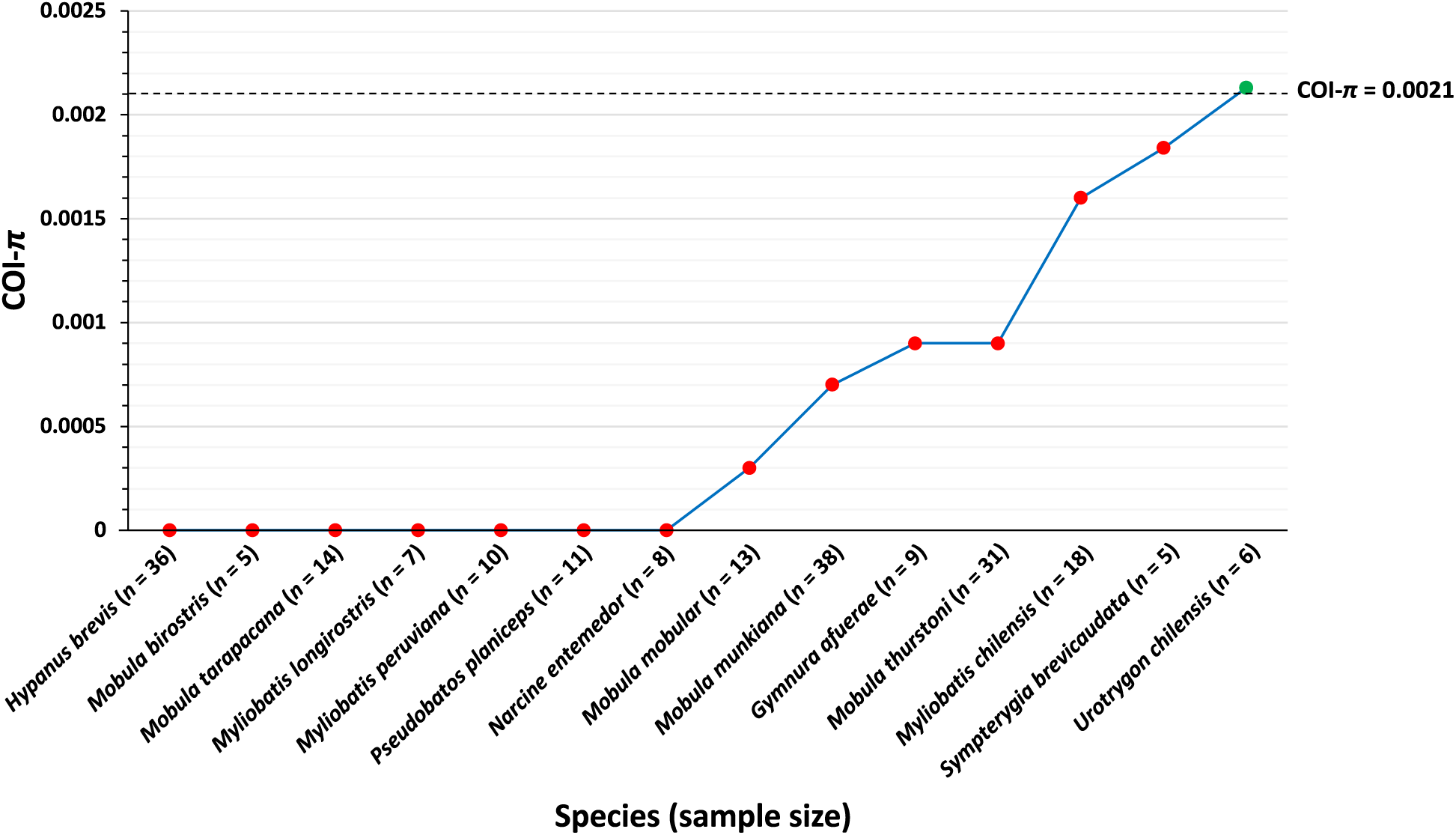
Intraspecific nucleotide diversity (*π*) of the cytochrome c oxidase subunit I gene (COI) across 14 Peruvian marine batoid species. The horizontal dotted line represents the low genetic diversity threshold (COI-*π* ≤ 0.0021) based on the Genetic Diversity Risk Indicator (GDRI) approach. Sample sizes (*n*) are indicated in parentheses along the x-axis

**Fig. S7.**
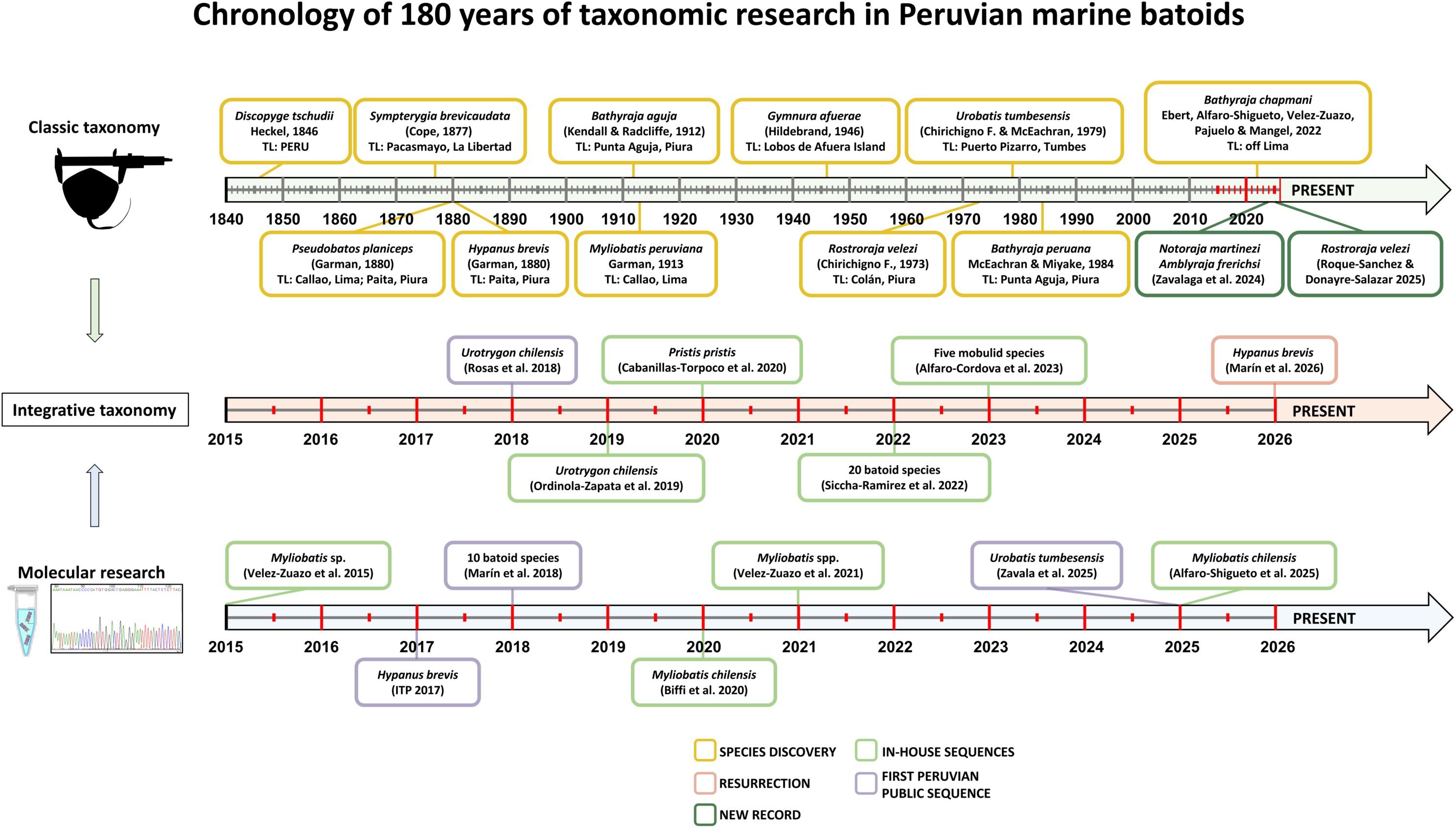
Timeline of Peruvian taxonomic and molecular research for marine batoids. Type locality: TL

**Table S1.** Multi-criteria decision analysis (MCDA) scoring rubric, parameters, and conditional threshold used for the prioritization of Peruvian batoid species

| Tier / Criteria | Parameter | Condition / Category | Assigned score | Framework application |
| --- | --- | --- | --- | --- |
| Tier 1: Core priority | IUCN conservation status (IUCN score) | CR | 5 | Baseline score calculated for all 37 species |
|  |  | EN | 4 |  |
|  |  | VU | 3 |  |
|  |  | NT | 2 |  |
|  |  | LC, DD, NE | 1 |  |
|  | ESPO endemic? (Endemism score) | Restricted to ESPO (YES) | 3 | Higher weight due to heightened vulnerability |
|  |  | Not restricted (NO) | 1 |  |
|  | Fisheries pressure (Fisheries score) | Primary commercial / High-volume landings | 3 | Scaled by landing volume and economic pressure |
|  |  | Secondary commercial / Low volume and low commercial value | 2 |  |
|  |  | No commercial value / Protected taxa | 1 |  |
| Tier 2: Genetic modifier | Nucleotide diversity ($\pi$) | Critically low diversity ($\pi$ = 0 to 0.0021) | +1 to final tier 1 rank | Triggers an immediate upgrade by one full category due to potential genetic vulnerability |
| | | Adequate diversity ($\pi$ > 0.0021) | 0 (no change) | Retains original tier 1 score |
| | | Missing sequence data (sample size ≤ 4) (GenBank / BOLD gap) | Flag as "High Priority (data gap)" | Highlights critical knowledge gaps for genetic data |
|  |  | IUCN status: Critically Endangered (CR); Endangered (EN); Vulnerable (VU); Near Threatened (NT); Least Concern (LC); Data Deficient (DD); Not Evaluated (NE)<br>Eastern South Pacific Ocean (ESPO) |  |  |

**Table S3.** Technical metadata and assembly statistics of the complete mitochondrial genome of *Notoraja martinezi*.

| Sample information |  |
| --- | --- |
| Species | *Notoraja martinezi* |
| BioSample | SAMN48911751 |
| Collection date | 2010-11-25 |
| Geographic location | Pacific Ocean: Costa Rica<br>9.3589° N, 85.44463° W |
| Submitted by | Smithsonian National Museum of Natural History |
| Study | NOAA Genome Skimming of Marine animals<br>inhabiting the US Exclusive Economic Zone |
| Run | SRR33849666 |
| Strategy | Whole-Genome Sequencing |
| Platform | Illumina NovaSeq X Plus |
| Assembly statistics |  |
| Total reads | 48,714,906 |
| Filtered reads | 20,000,000 |
| Reads mapped | 77,232 |
| Consensus assembled sequence size (bp) | 16,700 |
| Mean coverage | 524.9X |
| GenBank accession | BK075715 |

**Table S4.** Genetic characterization of the mitochondrial genome of *Notoraja martinezi*

| Gene | Start | End | Size (bp) | Strand | Start/Stop |
| --- | --- | --- | --- | --- | --- |
| trnF(gaa)PHE | 1 | 69 | 69 | + | — |
| 12S rRNA | 70 | 1,034 | 965 | + | — |
| trnV(tac)VAL | 1,035 | 1,106 | 72 | + | — |
| 16S rRNA | 1,127 | 2,784 | 1,658 | + | — |
| trnL2(taa)LEU | 2,785 | 2,859 | 75 | + | — |
| ND1 | 2,860 | 3,834 | 975 | + | ATG/TAA |
| trnI(gat)ILE | 3,835 | 3,904 | 70 | + | — |
| trnQ(ttg)GLN | 3,904 | 3,975 | 72 | - | — |
| trnM(cat)MET | 3,976 | 4,044 | 69 | + | — |
| ND2 | 4,045 | 5,088 | 1,044 | + | ATG/TAA |
| trnW(tca)SEC | 5,091 | 5,159 | 69 | + | — |
| trnA(tgc)ALA | 5,161 | 5,229 | 69 | - | — |
| trnN(gtt)ASN | 5,230 | 5,302 | 73 | - | — |
| trnC(gca)CYS | 5,336 | 5,400 | 65 | - | — |
| trnY(gta)TYR | 5,401 | 5,470 | 70 | - | — |
| CO1 | 5,472 | 7,028 | 1,557 | + | GTG/TAA |
| trnS2(tga)SER | 7,029 | 7,099 | 71 | - | — |
| trnD(gtc)ASP | 7,100 | 7,168 | 69 | + | — |
| CO2 | 7,174 | 7,864 | 691 | + | ATG/T-- |
| trnK(ttt)LYS | 7,865 | 7,938 | 74 | + | — |
| ATP8 | 7,940 | 8,107 | 168 | + | ATG/TAA |
| ATP6 | 8,098 | 8,781 | 684 | + | ATG/TAA |
| CO3 | 8,784 | 9,569 | 786 | + | ATG/TAA |
| trnG(tcc)GLY | 9,571 | 9,639 | 69 | + | — |
| ND3 | 9,640 | 9,988 | 349 | + | ATG/T-- |
| trnR(tcg)ARG | 9,989 | 10,057 | 69 | + | — |
| ND4L | 10,058 | 10,354 | 297 | + | ATG/TAA |
| ND4 | 10,348 | 11,728 | 1,381 | + | ATG/T-- |
| trnH(gtg)HIS | 11,729 | 11,797 | 69 | + | — |
| trnS1(gct)SER | 11,798 | 11,864 | 67 | + | — |
| trnL1(tag)LEU | 11,866 | 11,937 | 72 | + | — |
| ND5 | 11,938 | 13,773 | 1,836 | + | ATG/TAA |
| ND6 | 13,770 | 14,288 | 519 | - | ATG/TAA |
| trnE(ttc)GLU | 14,290 | 14,358 | 69 | - | — |
| CYTB | 14,362 | 15,504 | 1,143 | + | ATG/TAA |
| trnT(tgt)THR | 15,507 | 15,577 | 71 | + | — |
| trnP(tgg)PRO | 15,587 | 15,655 | 69 | - | — |
| Control region | 15,656 | 16,700 | 1,045 | + | — |

**Table S5.** GenBank/[BOLD] records used in the species delimitation analysis for *Rostroraja velezi* and the phylogeographic/population assessments for *Mobula mobular*, *Mobula munkiana*, *Sympterygia brevicaudata*, and *Urotrygon chilensis*

| Species | GenBank/[BOLD] | Geographic origin | Reference |
| --- | --- | --- | --- |
| **COI sequences used in species delimitation analyses for *Rostroraja velezi*** |  |  |  |
| *Rostroraja velezi* | **PZ899916** | **Tumbes (Peru)** | **This study** |
|  | **PZ899917** | **Tumbes (Peru)** | **This study** |
|  | **PZ899918** | **Tumbes (Peru)** | **This study** |
|  | **PZ899919** | **Tumbes (Peru)** | **This study** |
|  | [RDFCA219-05] | Costa Rica | — |
| *Bathyraja spinicauda* | KF604208 | Norway | Lynghammar et al. (2014) |
|  | KF604209 | Norway | Lynghammar et al. (2014) |
|  | KF604211 | Norway | Lynghammar et al. (2014) |
|  | KF604215 | Norway | Lynghammar et al. (2014) |
|  | KF604217 | Norway | Lynghammar et al. (2014) |
| *Raja asterias* | KT307374 | Algeria | — |
|  | KT307377 | Algeria | — |
|  | KT307384 | Sicily (Italy) | — |
|  | GU805133 | Sicily (Italy) | — |
|  | GU805135 | Sicily (Italy) | — |
|  | GU805389 | Marches (Italy) | — |
|  | GU805813 | Marches (Italy) | — |
| *Raja clavata* | KF604269 | Norway | Lynghammar et al. (2014) |
|  | KJ128596 | Sweden | — |
|  | JQ775100 | Portugal | Costa et al. (2012) |
|  | GU805373 | Marches (Italy) | — |
|  | GU805405 | Croatia | — |
| *Raja maderensis* | MZ661342 | Madeira (Portugal) | Crobe et al. (2021) |
|  | MZ661343 | Madeira (Portugal) | Crobe et al. (2021) |
|  | HM043185 | Portugal | Serra-Pereira et al. (2011) |
| *Raja straeleni* | JF494349 | South Africa | — |
|  | JF494353 | South Africa | — |
|  | GU805709 | Antarctica | — |
|  | GU805710 | Antarctica | — |
|  | GU805711 | Antarctica | — |
| *Rostroraja alba* | KT307419 | Italy | — |
|  | KT307420 | Italy | — |
|  | JF494387 | South Africa | — |
|  | JF494388 | South Africa | — |
|  | JF494389 | South Africa | — |
|  | JF494390 | South Africa | — |
|  | JF494391 | South Africa | — |
| *Rostroraja eglanteria* | MT455719 | Virginia Beach County (USA) | — |
|  | MT455929 | Atlantic Ocean (USA) | — |
|  | MT456247 | Atlantic Ocean (USA) | — |
|  | MH379043 | New Jersey (USA) | — |
|  | KT075315 | New Jersey (USA) | — |
|  | JF895098 | USA | Coulson et al. (2011) |
| *Tetronarce tremens* | KU737923 | Chile | — |
|  | KU737924 | Chile | — |
|  | KU737925 | Chile | — |
|  | KU737926 | Chile | — |
| **COI sequences used in phylogeographic analyses (haplotype network and AMOVA) for *Mobula mobular*** |  |  |  |
| *Mobula mobular* | **PZ899871** | **Piura (Peru)** | **This study** |
|  | **PZ899872** | **Piura (Peru)** | **This study** |
|  | **PZ899873** | **Tumbes (Peru)** | **This study** |
|  | **PZ899874** | **Lambayeque (Peru)** | **This study** |
|  | MH194451 to MH194453 | Tumbes (Peru) | Marín et al. (2018) |
|  | MH194455 | Tumbes (Peru) | Marín et al. (2018) |
|  | MH194457 | Tumbes (Peru) | Marín et al. (2018) |
|  | MH194461 | Tumbes (Peru) | Marín et al. (2018) |
|  | MH194464 | Tumbes (Peru) | Marín et al. (2018) |
|  | MH194469 | Tumbes (Peru) | Marín et al. (2018) |
|  | MH194482 | Lambayeque (Peru) | Marín et al. (2018) |
|  | KP175596 to KP175618 | Ecuador | — |
|  | KP175628 to KP175648 | Baja California Sur<br>(Mexico) | — |
| **COI sequences used in phylogeographic analysis (haplotype network) for *Mobula munkiana*** |  |  |  |
| *Mobula munkiana* | **PZ899875** | **Tumbes (Peru)** | **This study** |
|  | **PZ899876** | **Tumbes (Peru)** | **This study** |
|  | **PZ899877** | **Tumbes (Peru)** | **This study** |
|  | **PZ899878** | **Tumbes (Peru)** | **This study** |
|  | **PZ899879** | **Tumbes (Peru)** | **This study** |
|  | **PZ899880** | **Tumbes (Peru)** | **This study** |
|  | KM364898 to KM364900 | Baja California Sur<br>(Mexico) | Poortvliet et al. (2015) |
|  | KM364990 | Baja California (Mexico) | Poortvliet et al. (2015) |
|  | KX151645 | Baja California Sur<br>(Mexico) | White et al. (2018) |
| **COI sequences used in phylogeographic analysis (haplotype network) for *Sympterygia brevicaudata*** |  |  |  |
| *Sympterygia brevicaudata* | MH194468 | Tumbes (Peru) | Marín et al. (2018) |
|  | PP390310 | Piura (Peru) | Zavala et al. (2025) |
|  | PP390335 | Piura (Peru) | Zavala et al. (2025) |
|  | PP390355 | Piura (Peru) | Zavala et al. (2025) |
|  | PP390380 | Piura (Peru) | Zavala et al. (2025) |
|  | OQ472386 | Chile | Dufflocq et al. (2026) |
|  | KU737922 | Chile | Unpublished |
| **COI sequences used in phylogeographic analysis (haplotype network) for *Urotrygon chilensis*** |  |  |  |
| *Urotrygon chilensis* | **PZ899912** | **Lambayeque (Peru)** | **This study** |
|  | **PZ899913** | **Lambayeque (Peru)** | **This study** |
|  | **PZ899914** | **Lambayeque (Peru)** | **This study** |
|  | **PZ899915** | **Lambayeque (Peru)** | **This study** |
|  | KY572908 | Lima (Peru) | Rosas et al. (2018) |
|  | PP390357 | Piura (Peru) | Zavala et al. (2025) |
|  | PP236862 to PP236864 | Ecuador | Ehemann et al. (2024a) |

**Table S6.**
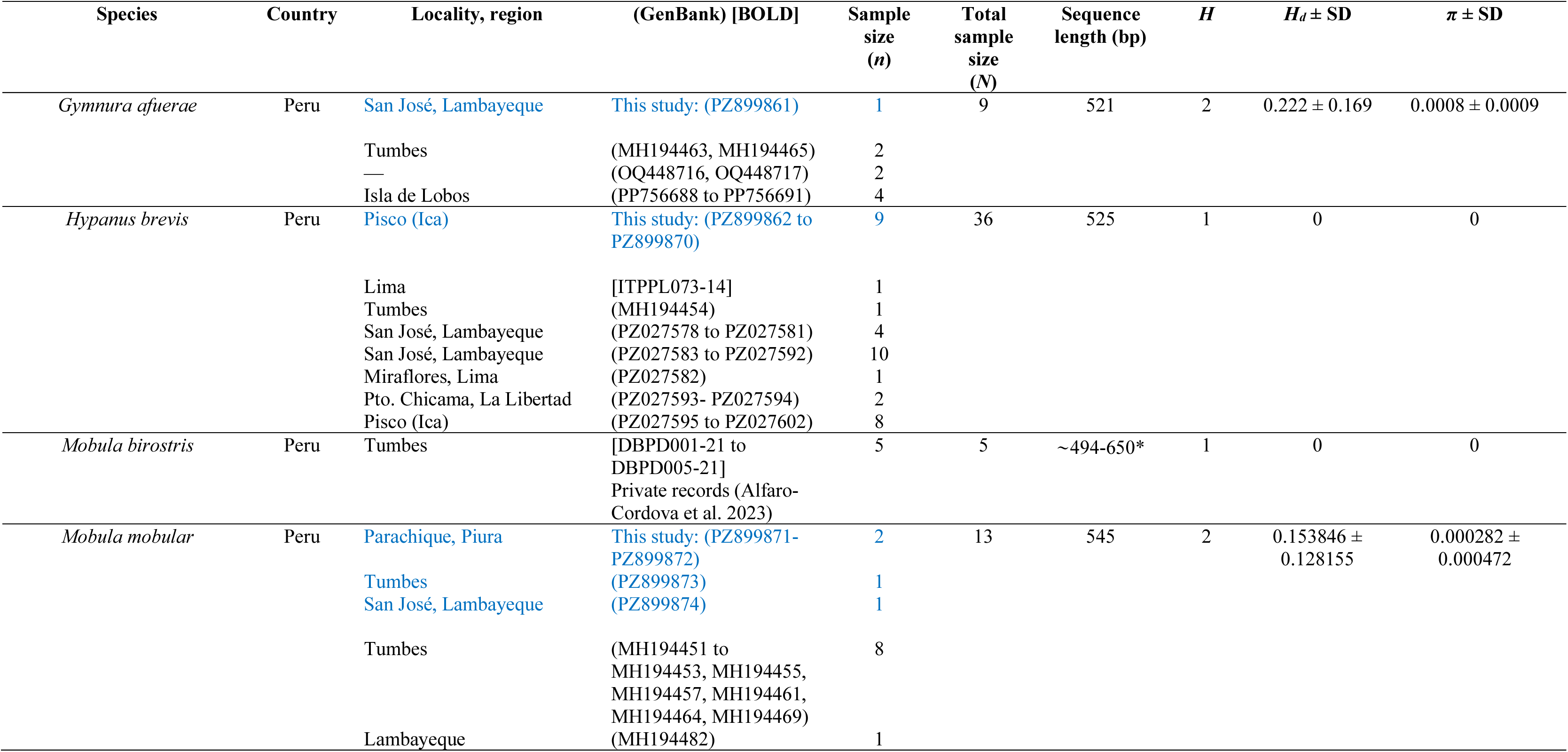
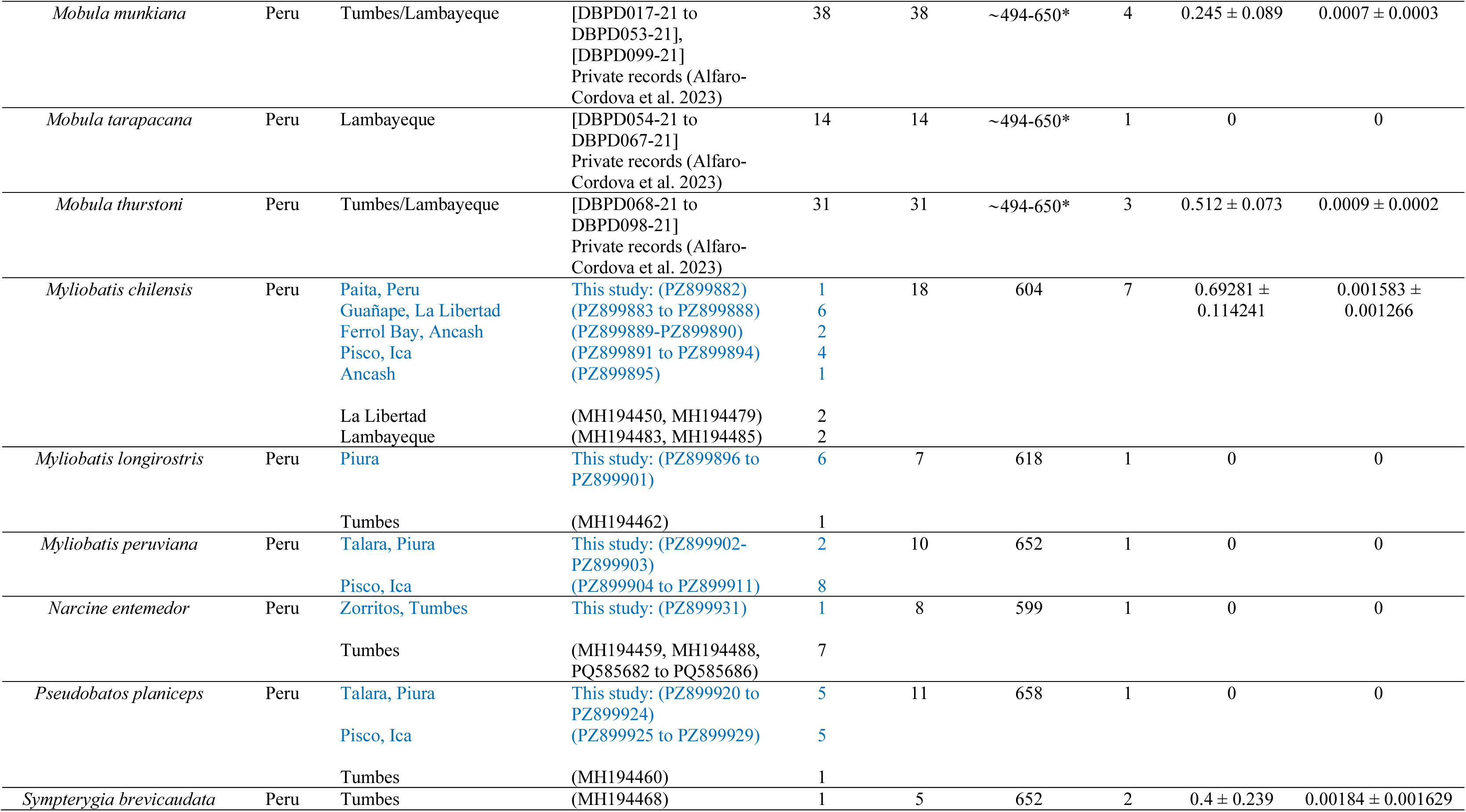
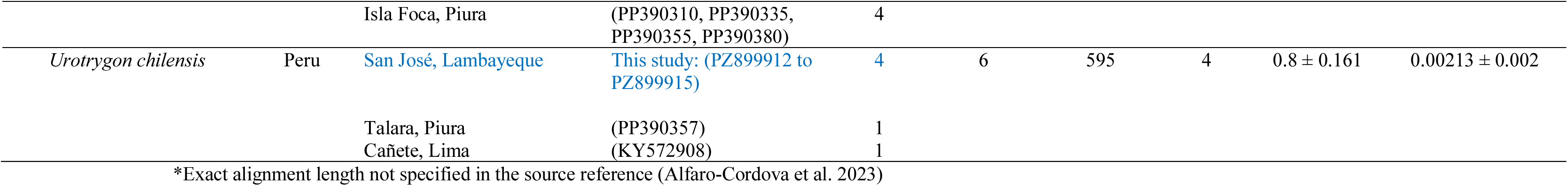
Diversity metrics for Peruvian batoid species based on barcodes from the cytochrome c oxidase subunit I gene (COI), including sample size (*n*), total sample size (*N*), number of haplotypes (*H*), haplotype diversity (*Hd*), and nucleotide diversity (*π*). Samples analyzed in the present study (BATOseq-PE library) are written in blue

## Notes

### Competing Interest Statement

The authors have declared no competing interest.

### Summary of Updates

This version of the manuscript has been revised to correct typos.

